# A Modular Genetic Code Expansion Approach to Site-Specific Lysine Acylations

**DOI:** 10.1101/2025.10.03.680225

**Authors:** Pascal Knecht, Tuan-Anh Nguyen, Kathrin Lang

**Affiliations:** Laboratory for Organic Chemistry, Department of Chemistry and Applied Biosciences, ETH Zurich, Zurich, Switzerland; CeMM Research Center for Molecular Medicine, Austrian Academy of Sciences, Vienna, Austria

## Abstract

Lysine acylations are a diverse class of post-translational modifications (PTMs) that dynamically regulate protein function. Access to homogenous, site-specifically modified proteins is essential for dissecting their molecular roles, yet remains challenging with traditional methods. Here we present a modular strategy that combines genetic code expansion (GCE) with a chemoselective amide bond-forming reaction to install a wide range of lysine acylations directly onto folded proteins. Key to this approach is the genetic encoding of *N*^ε^-methoxylysine, which reacts efficiently with *N*-methyliminodiacetyl (MIDA) acylboronates to generate diverse acyl lysine modifications from a single protein precursor, including several acyl PTMs not previously accessible through GCE. This mild and broadly applicable methodology enables systematic studies of lysine acylation on multiple target proteins. We demonstrate its utility by probing the effects of defined acyl modifications on enzymatic activity, protein-RNA interactions and deacylase-mediated regulation. Together, this platform establishes a versatile strategy to interrogate the functional consequences of this wide-spread class of PTMs.

## MAIN

Lysine acylations encompass a family of chemically diverse post-translational modifications (PTMs) in which the ε-amino group of lysine is linked via a stable amide bond to moieties ranging from short or long alkyl chains to hydroxy moieties, carboxylic acids, aromatic rings, and even entire proteins, thereby altering the charge, sterics, and chemical functionality of the protein scaffold.^1–3^ These modifications are installed either enzymatically, by dedicated writer enzymes, or non-enzymatically through reaction with activated metabolic intermediates. To dissect how individual lysine acylations influence protein function and to interrogate the activities of cognate reader and eraser proteins, it is essential to generate homogenous, site-specifically modified proteins. Genetic code expansion (GCE) provides a direct route to access defined modified proteins via site-specific incorporation of lysine derivatives bearing the respective acyl group. To date, hundreds of non-canonical amino acids (ncAAs) have been introduced into proteins, including many lysine acylations, mainly those bearing small and uncharged acyl moieties^4^. The direct installation of large, or negatively charged acyl lysine derivatives remains challenging due to cell permeability limitations, ncAA stability and the steric constrains of orthogonal aminoacyl tRNA synthetases (aaRSs).^5^

To overcome these limitations, we envisioned a ‘tag-and-modify’ approach, in which a chemoselective handle is genetically encoded and subsequently converted into the desired acyl PTM on purified proteins^6^. This required identifying an amide bond-forming reaction that operates under physiological conditions, at low micromolar protein concentrations and with near-quantitative conversions. Such a reaction would provide a modular route to diverse lysine acylations from a single ncAA precursor, eliminating the need to evolve a dedicated orthogonal aaRS for each target PTM, a particular advantage for charged and bulky acylations that proved to be very challenging aaRS-substrates^5^. Amide bonds are among the most fundamental linkages in organic chemistry and are typically formed by coupling amines with activated carboxylic acids. In small-molecule and peptide synthesis, this is achieved using coupling agents, often in the presence of catalysts or additives to improve yields and suppress side reactions. While highly effective in synthetic settings, these methods rely on protected substrates, strong activating agents, high reactant concentrations and organic solvents – conditions incompatible with folded proteins. Bioorthogonal reactions yielding amide bonds remain however rare. One example, the traceless Staudinger ligation, has been applied to react a genetically encoded azidonorleucine with acyl-modified phosphinothioesters to generate acetylated and succinylated proteins.^7^ The reaction is however accompanied by substantial side-product formation from azide reduction and suffers from slow kinetics leading to yields typically below 50%. Inspired by peptide ligation strategies pioneered by the Bode laboratory, we instead focused on the reaction of hydroxylamine derivatives with activated carboxylates to form amide bonds under mild aqueous conditions (Supplementary Fig. 1a)^8,9^. We envisioned to site-specifically incorporate a hydroxylamine-modified lysine derivative that could react on-protein with appropriately activated acyl donors to access endogenous acyl-PTMs (Fig. 1a). The α-ketoacid-hydroxylamine (KAHA) ligation, while powerful in peptide ligation, requires high reactant concentrations (ca. 5 mM) and acidic conditions, incompatible with most native proteins.^10^ In contrast, reaction of hydroxylamine derivatives with potassium acyltrifluoroborates (KAT ligation), particularly in the quinoline-activated form, yields amides at neutral pH and micromolar concentrations, but its demonstrated scope has been largely confined to activated quinoline-derived acyl donors, limiting its suitability for installing naturally occurring lysine acylations.^11^ An interesting modification of the KAT ligation involves N-methyliminodiacetyl (MIDA) acylboronates, enabling rapid, chemoselective amide bond formation with stable alkyl substituted hydroxylamines (Supplementary Fig. 1a).^12^

**Fig. 1.**
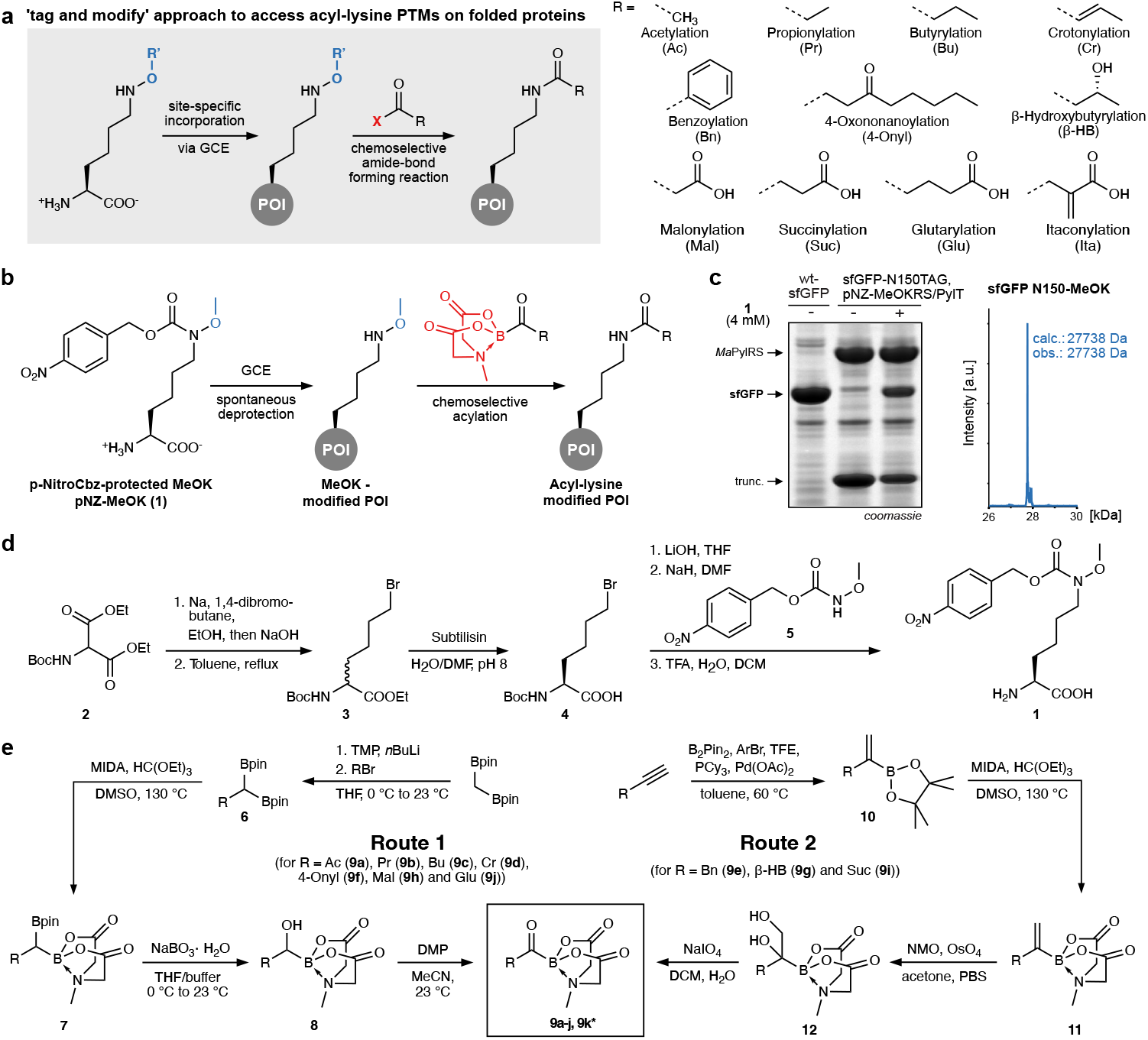
Synthesis and site-specific installation of MeOK via GCE. **a) ‘**Tag and modify’ approach to access acyl lysine PTMs on folded proteins. A hydroxylamine containing ncAA is incorporated into proteins via GCE, which is reacted with a suitable activated acyl donor. A variety of endogenous lysine acylations is displayed on the right. An *O-*methylhydroxylamine derivative of lysine bearing a *p*-nitrobenzyloxycarbonyl (pNZ) cage **1** is incorporated into proteins via GCE. The pNZ group is removed by endogenous *E. coli* nitroreductases after incorporation. Reaction of the MeOK-modified POI with a MIDA acylboronate yields the desired acyl-lysine modified POI, including **c)** SDS-PAGE and MS analysis confirming incorporation of pNZ-MeOK into sfGFP in response to an amber stop codon. Full-length MS analysis confirms cleavage of the pNZ cage. Consistent results were obtained over three distinct replicate experiments. **d)** Synthesis scheme to access pNZ-MeOK **1. e)** Schematic representation of the two routes, which were used for the preparation of various acylboronates. *MIDA itaconylboronate **9k** was prepared following a different route (depicted in Scheme S3).

In this work, we present a general and modular platform for installing a broad spectrum of lysine acylations, including bulky and negatively charged PTMs, directly onto folded proteins from a single genetically encoded precursor. Site-specific incorporation of *N*^ε^-methoxylysine (MeOK) introduces a stable hydroxylamine functionality into a protein of interest (POI). Subsequent reaction with synthetically accessible MIDA acylboronates proceeds under mild, aqueous conditions, yielding diverse lysine acylations, several of which have not previously been accessible by genetic code expansion. We demonstrate the scope of this chemistry by generating eleven distinctly acylated variants of ubiquitin (Ub), probing the functional consequences of lysine acetylation on isocitrate dehydrogenase (IDH) activity, and examining the impact of acylation on RNA binding of glyceraldehyde-3-phospate dehydrogenase (GAPDH). Furthermore, we show that these site-specifically acylated proteins serve as substrates for deacylating enzymes, underscoring their biological relevance. Our approach provides a versatile route to homogenous, site-specifically acylated proteins and offers a powerful tool for dissecting the mechanistic roles of lysine acylations in protein regulation.

## RESULTS AND DISCUSSION

### Design and site-specific encoding of a hydroxylamine-bearing lysine derivative

To access site-specific lysine acylations on folded proteins via the ‘tag and modify’ approach, we initially sought to employ KAT ligation, a method known for its efficiency in ligating unprotected peptide fragments under aqueous conditions.^13^ Although KAT ligation has only rarely been applied to folded proteins, previous studies demonstrated its feasibility by first attaching a cysteine-reactive *O*-carbamoylhydroxylamine handle to a target protein, followed by conjugation with quinolyl or pyridyl KATs at acidic or neutral pH (Supplementary Fig. 1b)^11,14^. Inspired by this work, we designed and synthesized a series of *O*-carbamoylhydroxylamine-bearing lysine derivatives (Supplementary Fig. 1c). However, these lysine derivatives proved to be unstable under physiological conditions, and we were unable to identify specific orthogonal aaRS variants for their genetic encoding.

Consequently, we turned to more stable *O*-alkylhydroxylamine lysine derivatives that are expected to undergo amide bond formation with *N*-methyliminodiacetyl (MIDA) acylboronates (Supplementary Fig. 1a)^12^. Based on preliminary data, we identified *O*-methylhydroxylamine as a moiety that is stable in the presence of millimolar concentrations of reactive cellular metabolites such as glutathione and pyruvate, suggesting its compatibility with standard conditions used for recombinant protein expression and purification. To enable access to native lysine acylations via reaction with MIDA acylboronates, we therefore envisioned to synthesize and genetically encode N^ε^-methoxy-L-lysine (MeOK) into target proteins (Fig. 1b). Given that the methoxy substitution on the ε-amino group of lysine represents only a minor structural alteration, we anticipated that orthogonal aaRS variants might struggle to distinguish this ncAA from lysine. To address this challenge, we synthesized derivative **1** that besides the methoxy group bears a p-nitrobenzyloxy-carbonyl (pNZ) protecting group on the ε-amino group of lysine (Fig. 1b). We and others have previously shown that this protection group is cleaved spontaneously after protein expression, triggered by reduction of the aromatic nitro group, followed by fragmentation, thereby restoring the reactive hydroxylamine functionality on the target protein.^15–17^ We synthesized caged hydroxylamine ncAA **1** (pNZ-OMeK) by first alkylating diethyl 2-((tert-butoxycarbonyl)amino)malonate (2) with 1,4-dibromobutane (Fig. 1d and Scheme S1).^18^ Subsequent ester hydrolysis and decarboxylation led to a racemic mixture of bromoalkyl amino acid ester **3**. Incubation with subtilisin allowed separation of the enantiomers by only hydrolysing the ethyl-ester of the L-amino acid leading to 6-bromo-*N*-boc-L-norleucine (4), which was reacted with 4-nitrobenzyl methoxycarbamate (5) that was prepared from commercially available 4-nitrobenzyl chloroformate and methoxyamine hydrochloride. Final Boc deprotection afforded ncAA **1** at multigram scale (Fig. 1d and Supplementary Information).

To genetically encode ncAA **1** we screened the ability of > 200 pyrrolysine tRNA synthetase (PylRS) mutants available in the lab for suppressing an amber codon introduced into superfolder green fluorescent protein (sfGP-N150TAG) in the presence of the corresponding tRNA_CUA_ (PylT). Excitingly, a few PylRS mutants from *Methanomethylophilus alvus* (*Ma*) showed weak activity for aminoacylating PylT with ncAA **1**. In order to improve incorporation yields, we created - based on mutation sites in these variants - a cassette mutagenesis library of *Ma*PylRS via randomization of five residues (L125, Y126, M129, H227, Y228) and subjected this library to alternating rounds of positive and negative selection in *Escherichia coli* in the presence of **1**.^19,20^ Gratifyingly, a novel *Ma*PylRS variant (Y126W, M129C, H227S and Y228F) led to good incorporation yields of **1** (Fig. 1c). Purification of amber suppressed sfGFP and mass spectrometric (MS) analysis revealed the correct mass for MeOK-modified sfGFP, confirming that the pNZ cage was indeed cleaved upon protein expression (Fig. 1c). This was corroborated by recombinant expression, purification and LC-MS analysis of other amber-containing target proteins expressed in the presence of **1** (Ub-K63MeOK, Ub-K48MeOK, SUMO-K11MeOK and SUMO-K45MeOK (Supplementary Fig. 2)).

With MeOK-bearing POIs in hand, we were well-positioned to explore the reaction between hydroxylamines and MIDA acylboronates for site-specific installation of acyl-lysine PTMs on folded proteins.

### Synthesis of suitable MIDA acylboronates

To access MIDA acylboronates suitable for site-specific installation of acyl lysine PTMs on MeOK-modified POIs (Fig. 1a), we employed one of the two synthetic routes outlined in Fig. 1e.^21–23^ For route 1, we first alkylated bis[(pinacolato)boryl]methane with the appropriate substituent corresponding to the desired acylation.^24^ Subsequently one of the geminal pinacol boronic ester (BPin) groups was transformed into a B(MIDA) moiety by heating in the presence of methyliminodiacetic acid. Oxidation of the remaining C-BPin bond with sodium perborate and further oxidation of the resulting alcohol afforded the desired MIDA acyl boronates **9**.^21^ In some instances, it was more feasible to synthesize the MIDA reagents according to a different procedure, depicted as route 2 in Fig. 1e.^22,23^ For this, a suitably substituted terminal alkyne was first subjected to hydroboration to afford the corresponding alkenyl-2-pinacol boronic ester **10**. Subsequently, the BPin moiety was exchanged for B(MIDA) under the same conditions as in route 1. The alkene functionality was then subjected to dihydroxylation, followed by oxidative cleavage using sodium periodate to yield the desired MIDA acyl boronates **9**. Following one of these two routes, we generated MIDA acylboronates with acetyl (Ac, **9a**), propionyl (Pr, **9b**), butyryl (Bu, **9c**), crotonyl (Cr, **9d**), benzoyl (Bn, **9e**), 4-oxononanoyl (4-Only, **9f**), β-hydroxybutyryl (β-Hb, **9g**), malonyl (Mal, **9h**), succinyl (Suc, **9i**) and glutaryl (Glu, **9j**) moieties. To access MIDA itaconylboronate **9k**, we pursued an alternative synthetic approach (see Scheme S3). In the key step, we followed a literature procedure describing the attachment of *in situ* generated MIDA formylboronate by organometallic nucleophiles.^25^ Details on synthesis, yields and characterization for all MIDA acylboronates can be found in the Supplementary Information.

### Installing site-specific lysine acylations

We next tested and optimized conditions for reacting MeOK-modified POIs with the synthesized MIDA acylboronates. As first test protein we selected ubiquitin (Ub), bearing MeOK at position 63, which resides in a flexible loop before the C-terminal ß-sheet and is surface exposed and thereby well accessible^26^. Incubation of Ub-K63MeOK (20 µM) in aqueous buffer (50 mM citrate buffer, 150 mM NaCl) with 100 equivalents MIDA acetylboronate (**9a**, 2 mM) afforded fully acetylated protein (Ub-K63AcK) within two to three hours at pH 7.0 as judged by LC-MS (Fig. 2a and Supplementary Fig. 3 and 4). Excitingly under similar conditions (typically 10-20 µM POI, 2 mM MIDA acylboronate, 50 mM citrate, 150 mM NaCl, pH 6.5-7.0, 3-17 hours), Ub-K63MeOK at position 63 reacted with all synthesized MIDA acylboronates (**9a-k**), giving straight-forward and modular access to eleven different site-specific acyl lysine PTMs starting from one single MeOK-modified POI (Fig. 2b, Supplementary Fig. 5 and 6). Importantly, this includes PTMs that could not be directly accessed previously, such as negatively charged acylations (e.g. malonylation, succinylation, glutarylation or itaconylation) as the corresponding ncAAs are poorly cell permeable^27^ and no orthogonal aaRS/tRNA pairs could be evolved for their direct incorporation.^5,7^

**Fig. 2.**
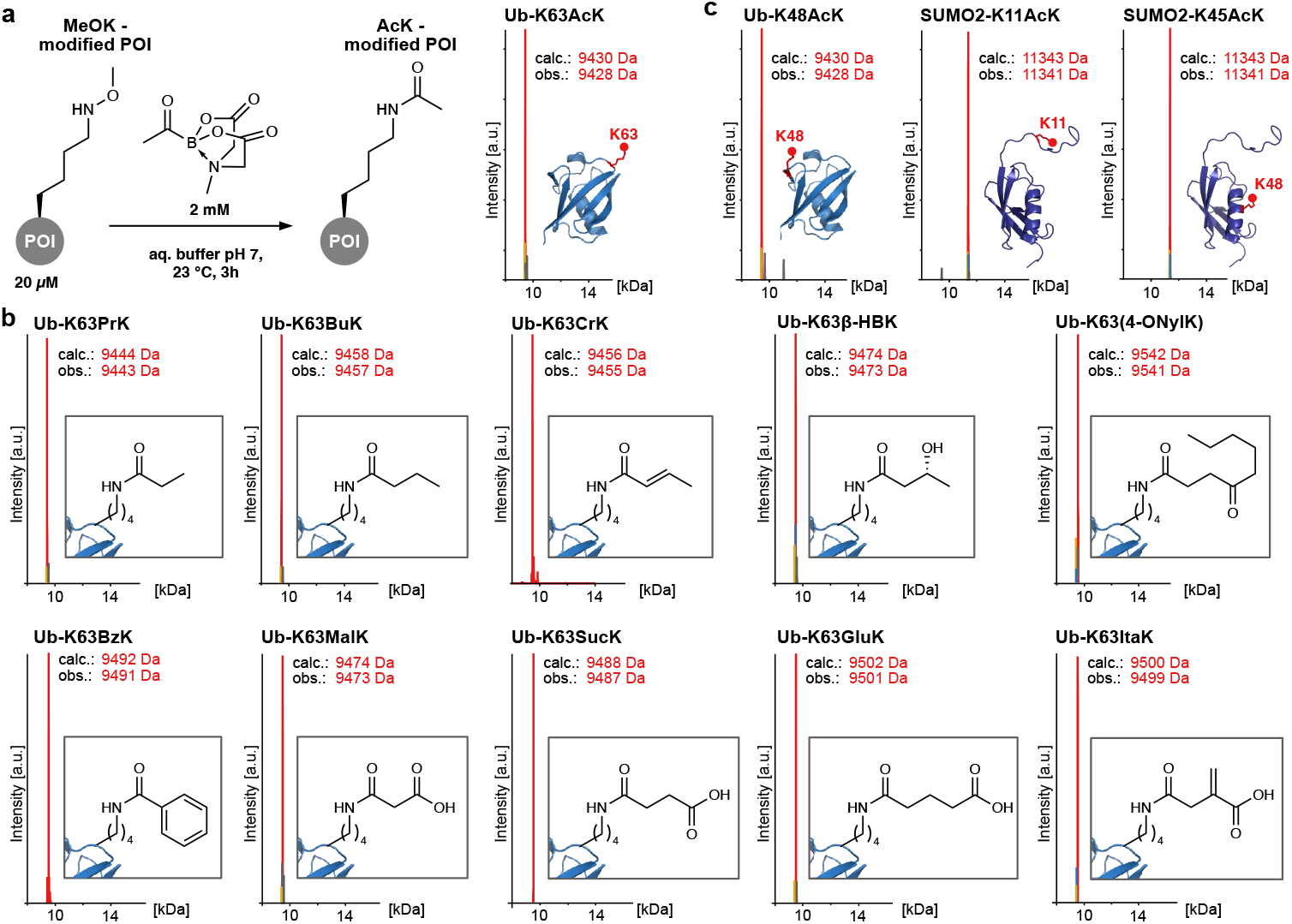
Installing site-specific lysine acylations. **a)** Schematic representation for installing acyl lysine modification on a MeOK-modified POI with applied conditions on the left. Full-length MS analysis of Ub-K63AcK after installation. **b)** Full-length mass spectra of various installed lysine acylations via the described method. For reaction conditions see Supplementary Fig. 5 and 6. **c)** Full-length masses of Ub-K48AcK, SUMO2-K11AcK and SUMO2-K45AcK installed under reaction conditions displayed in a). The modified residue is highlighted in red on the protein structure. The mass of the desired acylated protein is displayed in red, the hydroxylamine starting material in blue and the mass of unmodified lysine in yellow. PDB codes: 1UBQ and 2N1W.^26^ Consistent results were obtained over three distinct replicate experiments.

To further show the generality and modularity of our approach, we examined the MIDA acyl-boronate-mediated acylation at other positions within Ub (e.g. Ub-K48MeOK) and on other POIs bearing site-specifically encoded MeOK (e.g. SUMO2-K11MeOK and SUMO2-K45MeOK). Under the conditions described above, all tested POI-MeOK variants were efficiently and near-quantitatively acetylated using the corresponding MIDA acetylboronate **9a** (Fig. 2c, Supplementary Fig. 3 and 4). In all cases, MS analysis revealed a single dominant peak corresponding to successful and selective formation of the desired acetylated products. In some instances, a minor mass signal, attributable to lysine was observed. This suggests a high degree of selectivity and efficiency, with minimal side products for our acylation approach. While Ub lacks cysteine residues, the efficient and near-quantitative modification of SUMO2 variants bearing MeOK at either position 11 or 45, demonstrates that the MIDA acetylboronate-mediated reaction proceeds effectively even in the presence of nearby cysteine residues. Notably, SUMO2 contains a cysteine residue at position 48, proximal to one of the modification sites, yet the reaction efficiency remains uncompromised. This indicates that neither the MIDA acetylboronate reagent nor its reaction intermediates are adversely impacted by proximal nucleophilic side chains, underscoring the chemoselectivity and versatility of the method in complex protein environments. Importantly, the same holds true for the crotonyl and itaconyl boronate reagents. Despite featuring potential Michael acceptor motifs, both reagents show no or only minor reactivity with SUMO2-wild type (wt) under the employed conditions (20 µM SUMO2, 1 mM acylboronate, pH 5.5). In contrast, near-quantitative crotonylation or itaconylation are observed when SUMO2 displays a site-specifically incorporated MeOK at position 11 or 45 (Supplementary Fig. 7).

### Caged hydroxylamine lysine derivatives for small-molecule activation

Upon incorporation of ncAA **1** into various target proteins, we observed that the efficiency of pNZ deprotection varied depending on POI, the specific incorporation site, and the conditions used for expression and purification. In particular, when ncAA **1** was genetically encoded at more occluded and less solvent-exposed sites within a protein structure (e.g. position K27 in Ub) or into proteins prone to forming inclusion bodies during recombinant protein expression in *E. coli* (e.g. histones), the resulting purified protein typically consisted of a mixture of both the intact caged form of **1** and the decaged MeOK form (Fig. 3a and Supplementary Fig. 8a). The exact process by which the pNZ group is cleaved upon protein expression remains poorly understood. It is generally proposed that the nitro group of the incorporated ncAA **1** is reduced by cellular factors to its corresponding hydroxylamine or amine counterpart, which triggers a 1,6-elimination reaction, resulting in the release of carbon dioxide and uncaging of ncAA **1** (Supplementary Fig. 8b).^28^ Several *E. coli* nitroreductases (NTRs) have been implicated in the reduction of aromatic nitro groups in peptides and proteins.^29,30^ Particularly, the NAD(P)H-dependent NTRs NsfA and NsfB have demonstrated high activity toward the reduction of nitro groups.^28–31^ Amber suppression experiments in the double knockout Δ*nsfA*Δ*nsfB* K12 strain in the presence of **1** resulted in only partial pNZ decaging, even for POIs and positions that showed quantitative decaging to MeOK when expressed in wt-K12 cells (e.g. sfGFP-N150TAG, Ub-K63TAG, Supplementary Fig. 8c). This indicates that NsfA and/or NsfB are indeed involved in nitro group reduction of **1** and are essential for efficient decaging to MeOK. In vitro incubation of these partially decaged protein variants with either purified NsfA or NfsB resulted in decaging (Supplementary Fig. 8d). Unfortunately, this was not the case for proteins where incorporation sites for **1** resided in more occluded areas such as position 27 in Ub. Expression of Ub-K27TAG in presence of **1** in wt-K12 yielded a Ub species in which the majority of the pNZ cage remained intact with only a small fraction of the protein fully decaged to Ub-K27MeOK (Fig. 3a). Subsequent incubation of this protein mixture with purified NsfA or NsfB did not result in complete decaging to Ub-K27MeOK, likely due to restricted access of NsfA and NsfB to the occluded modification site (Fig. 3b).

**Fig. 3.**
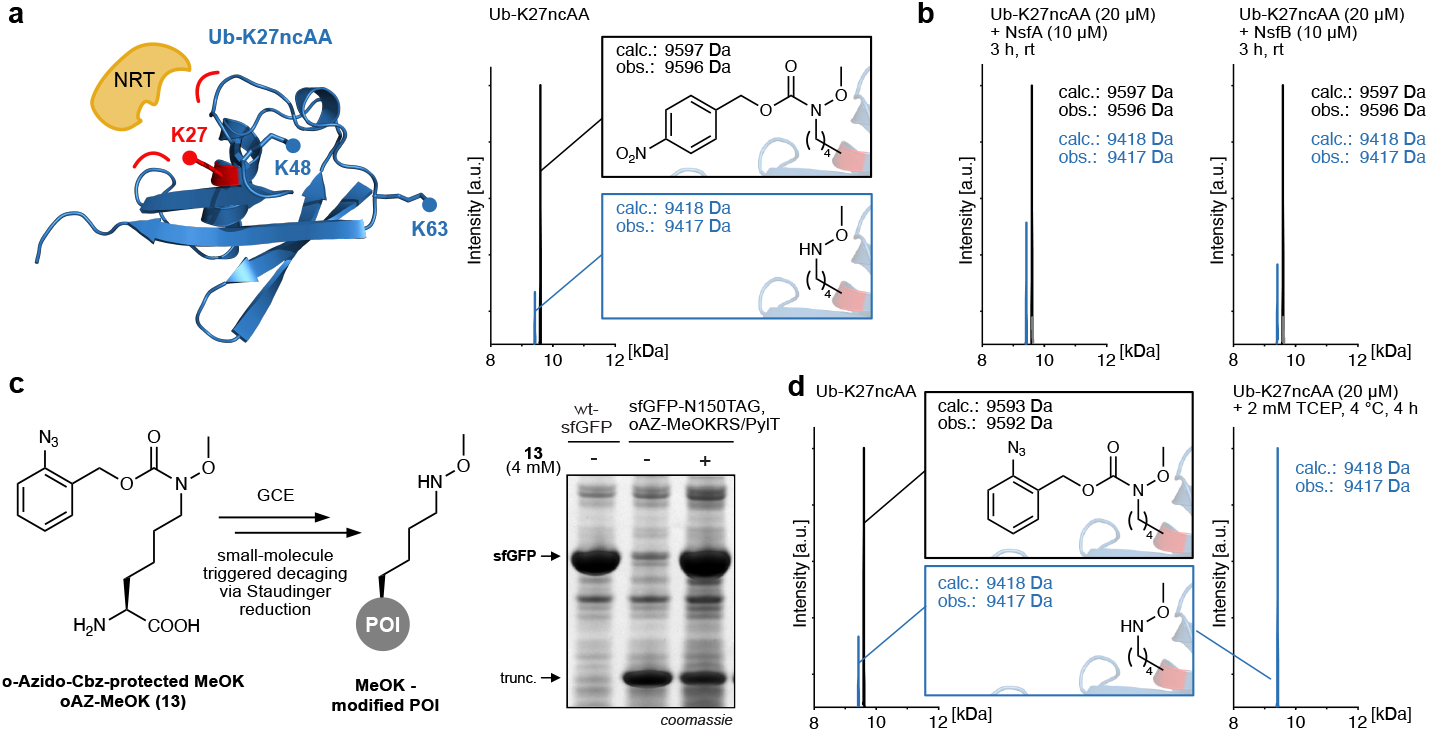
A MeOK derivative that can be decaged by a small molecule. **a)** Left: Schematics showing that lysine 27 in Ub is more sterically occluded than e.g. K48 and K63, thereby likely hampering access of nitroreductases (NRTs) to ncAAs installed at this position (PDB: 1UBQ)^26^. This leads to only partially decaged MeOK-bearing Ub. Right: full-length mass spectrum of Ub-K27TAG expressed in the presence of pNZ-MeOK. The peak with the mass corresponding to the ncAA with the intact pNZ cage is coloured in black, the desired MeOK-modified POI in blue. **b)** MS analysis show that incubation of Ub-K27pNZ-MeOK with recombinantly expressed *E. coli* nitroreductases NfsA or NfsB does not lead to full pNZ decaging. **c)** Small-molecule triggered decaging approach to access MeOK-bearing POIs. ncAA 13 bearing an oAZ-cage is site-specifically incorporated via GCE, followed by Staudinger reduction leading to MeOK-modified POI. SDS-PAGE analysis shows very efficient incorporation of oAz-MeOK (13) into sfGFP. **d)** Full-length MS analysis shows that oAZ-MeOK at K27 in ubiquitin stays mainly intact after Ub expression and purification. Treatment of purified Ub variant with 2 mM TCEP for 4 hours at 4 °C leads to complete deprotection and clean MeOK-bearing Ub. Consistent results were obtained over three distinct replicate experiments.

To broaden the scope of target proteins amenable to our modular acylation approach, we sought of developing a MeOK-derivative that can be unmasked by small molecules. Inspired by reports on phosphine-induced Staudinger reduction of aryl azides^32,33^, we designed ncAA **13**, an o-azidobenzyloxycarbonyl-caged MeOK derivative (oAZ-MeOK (13), Fig. 3c). Synthesis proceeded in analogous fashion to synthesis of pNZ-MeOK by reacting 6-bromo-N-boc-L-norleucine (4) with the corresponding o-azidobenzyl methoxycarbamate followed by Boc-deprotection to yield **13** (see Scheme S2). Screening of our in-house library of over 200 PylRS variants for incorporation efficiency of **13** into sfGFP-N150TAG identified several positive hits. Notably, the *Methonosarcina mazei* (*Mm)* PylRS variant carrying the Y306A and Y384F mutations - previously reported as highly promiscuous PylRS variant for a broad range of ncAAs^34^, exhibited particularly high incorporation efficiency for **13** (Fig. 3c). Purification and MS analysis of amber codon-containing proteins in the presence of **13** revealed site-specific incorporation of **13** and - depending on POI and position - partial cleavage to MeOK due to azide reduction in living *E. coli* (Fig. 3d). Excitingly, incubation with low concentrations of tris(2-carboxyethyl)phosphine (TCEP, 2 mM), a common additive during protein purification, resulted in clean and quantitative decaging to MeOK-containing POI (Supplementary Fig. 9), even for sterically occluded positions, such as residue K27 within Ub (Fig. 3d).

### Modular acylation recapitulates the effects of genetically encoded PTMs

With a modular tool in hand to site-specifically introduce various lysine acylations at user-defined sites into diverse target proteins, we next sought to investigate whether the effects of PTMs introduced via our modular acylation strategy replicate those introduced through direct incorporation via genetic code expansion. As a first proof-of-principle target we selected IDH from *E. coli*, a NADP^+^-dependent enzyme in the tricarboxylic acid cycle that functions as a homodimer and catalyses the conversion of isocitrate to α-ketoglutarate, producing NADPH (Supplementary Fig. 10a)^35^. IDH is classically regulated through reversible phosphorylation of a serine residue in its active site, which modulates its enzymatic activity to balance metabolic flux between energy generation and biosynthesis.^36^ In addition to phosphorylation, IDH is known to undergo extensive lysine acetylation but the effects of these modifications are less studied.^37,38^ In order to see the compatibility and applicability of our modular acylation strategy on biologically relevant protein systems, we chose to prepare an IDH variant, bearing acetylated lysine at residue 242, and studying its influence on enzymatic activity. K242 is located on a surface-exposed helix near the dimer interface (Supplementary Fig. 10a)^39^. It is not directly involved in substrate or cofactor binding, but its acetylation may modulate enzymatic activity through structural and allosteric effects. We prepared IHD-K242AcK either by direct incorporation of AcK using the previously published *M. barkeri* (*Mb)* PylRS-variant AcKRS3^40^ or by site-specific acetylation of IDH-K242MeOK with the corresponding MIDA acetylboronate (10 µM IDH-K242MeOK, 3 mM MIDA acetylboronate, pH 6.5, 5 hours at rt). Purity and identity of the IDH variants were assessed by SDS-PAGE and LC-MS (Fig. 4 a,b and Supplementary Fig. 10b). Importantly, the IDH-wt variant treated under the same conditions used for site-specific acetylation of IDH-K242MeOK did not show any evidence of non-specific modification or degradation (Fig. 4b and Supplementary Fig. 10b).

**Fig. 4.**
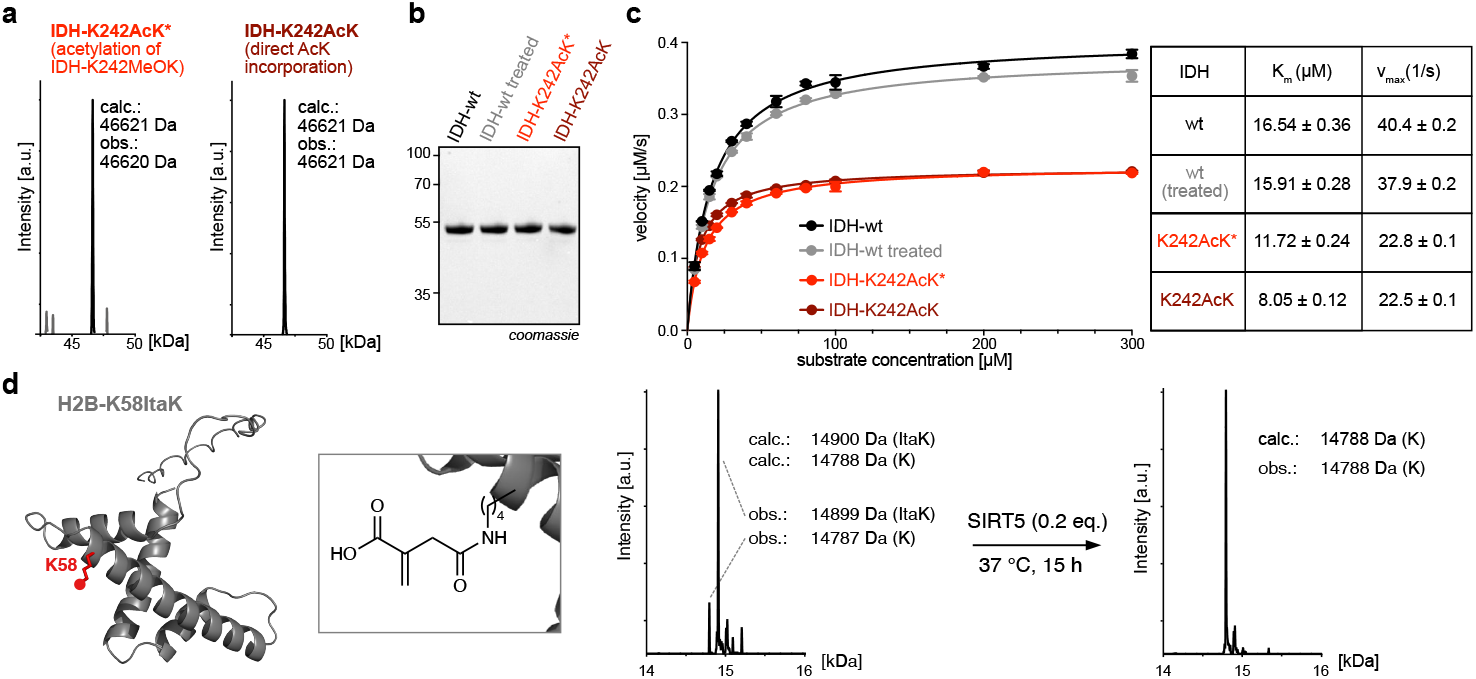
Site-specifically acylated POIs allow studying acylation effects and deacylase specificities. **a)** Full-length MS analysis of IDH-K242AcK prepared by either acetylation of OMeK-bearing IDH (IDH-K242AcK*, left)) or via direct AcK incorporation (IDH-K242AcK, right). **b)** SDS-PAGE analysis of different IDH variants. **c)** Michaelis-Menten kinetics of IDH-wt and IDH-242AcK variants. Both AcK variants show practically identical reduction in activity. IDH-wt treated denotes an IDH-wt sample treated under identical conditions as used for IDH-OMeK acylation and serves as control of preserving enzyme integrity showing barely any deviation from IDH-wt. The initial reaction velocity (*v*_*0*_) was plotted against isocitrate concentration and fitted with a Michaelis-Menten model to obtain values for *K*_*m*_ and *v*_*ma*x_. Average values and errors (±s.e.m.) were calculated from three biologically independent experiments (*n* = 3). Data was processed unsing GraphPad Prism 10 (GraphPad software.) **d)** Alphafold prediction of histone H2B with residue K58 highlighted in red with a zoom displaying K58 itaconylation (left)^41,42^. Full-length mass spectrum of itaconylated histone H2B before and after incubation with deacylase SIRT5, which completely removes the modification (right).

To evaluate the enzymatic activity of the different IDH variants, we employed an assay to monitor the consumption of D,L-isocitrate by measuring the increase in NADPH absorbance at 340 nm upon incubation of IDH with NADP^+^ and varying concentrations of D,L-isocitrate. This allowed us to determine the maximal velocity (v_max_) and the Michaelis-Menten constant (K_m_) of the IDH variants (Fig. 3c, Supplementary Fig. 11-12). IDH-wt exhibited kinetic parameters consistent with literature values, with K_m_ values in the low µM range.^35,43^ Notably, IDH-wt subjected to the acylation conditions displayed identical Michaelis-Menten parameters, confirming that the treatment itself does not impact proper enzyme folding and structural integrity. In contrast, the IDH-K242AcK variants, prepared either via direct AcK incorporation or via site-specific acetylation, showed a 60-70% reduction in both K_m_ and v_max_, likely by altering inter-subunit interactions and weakening homodimer stability, possibly through changes in conformational flexibility or disruption of allosteric communication. Importantly the AcK-variants, obtained either through direct AcK incorporation or through modular acylation perform nearly identical in the conducted enzymatic assay.

### Probing specificities of deacylases

Our modular acylation strategy enables the introduction of diverse acyl PTMs by reacting one POI bearing a site-specific MeOK residue with various MIDA acylboronates. The resulting acylated proteins faithfully replicate the structural and functional properties of their endogenous counterparts, making our approach a powerful and versatile platform for probing the activity, substrate scope and mechanistic preferences of deacylases. This is especially valuable for novel, under-studied acyl PTMs and for acylations that cannot be directly installed through GCE or more traditional enzymatic methods.

As a proof of concept, we prepared a panel of Ub variants featuring acetyl-, malonyl-, succinyl- or itaconyl-modified lysine at position 63 by incubating an Ub-K63MeOK variant with the corresponding acylation reagents (**9a, 9h, 9i** and **9k**, Fig. 1e). The modified Ub variants were purified and incubated with SIRT5, a member of the sirtuin family of the NAD^+^-dependent lysine deacylases. Unlike other sirtuins, such as SIRT1 and SIRT2, which primarily act as deacetylases, SIRT5 is best known for its role in regulating mitochondrial metabolism by removing non-acetyl lysine acylations such as succinylation, malonylation and glutarylation^44,45^. Indeed Ub-K63AcK remained unmodified after incubation with SIRT5. In contrast, the negatively charged acyl modifications (malonyl, succinyl, itaconyl) were either fully or partially removed under the tested conditions, as proved by LC-MS (Supplementary Fig. 13).

Lysine itaconylation is a recently discovered non-enzymatic PTM^46^, in which itaconyl-CoA, a metabolic derivative of itaconate, reacts with the ε-amino group of lysine residues. Itaconylation has been implicated in immunoregulation and quantitative proteomics identified itaconylation sites in multiple functional proteins, including glycolytic enzymes and histones. Whether this modification arises solely from non-enzymatic reactions in response to elevated itaconate/itaconyl-CoA levels or is also dynamically regulated by specific writer and eraser enzymes remains to be investigated. The identification of such regulators, as well as of potential reader enzymes requires however the development of reliable methods to access site-specifically itaconylated target proteins. Among the most heavily itaconylated proteins, histone H2B type1-B (H2B1B) was identified^46^. Given that H2B1B is a known substrate for several epigenetically relevant acylations, it is plausible that itaconylation could exert similar functional effects. To explore how itaconylation at this site might be regulated, we incorporated ncAA **13** into H2B bearing an amber codon at position 58, followed by deprotection with TCEP (1 mM) during purification, to obtain homogenous H2B-K58MeOK. Reaction with MIDA itaconylboronate **9k** gave access to site-specifically modified H2B-K58ItaK, as confirmed by LC-MS (Fig. 4d). Upon incubation of this variant with SIRT5, complete deacylation was observed, demonstrating that SIRT5 is active on native itaconylated substrates (Fig. 4d). We anticipate that access to site-specifically itaconylated target proteins will unlock new opportunities for in-depth functional and mechanistic studies of this novel PTM. This will aid in uncovering the roles of potential writer, reader, and eraser enzymes in regulating itaconylation and its broader biological functions.

### Succinylation influences RNA binding of GAPDH

To further study functional and structural consequences of acylation PTMs, we turned our focus to GAPDH, an NAD^+^-dependent enzyme that is central to glycolysis.^47^ Beside its metabolic function in the glycolytic pathway, GAPDH is implicated in a variety of cellular processes (so called moonlighting functions), including nucleic acid binding^48^, DNA replication and repair^49^, apoptosis and cytoskeletal interactions.^50^ There is growing evidence that this functional versatility is at least partly governed by PTMs, which modulate its catalytic activity^51^, interaction profile as well as subcellular localization.^52^ We have recently studied how glutarylation of a lysine residue (K194) close to the cofactor binding site influences its glycolytic activity.^5^ In the cytosol, GAPDH functions as a tetrameric enzyme (Fig. 5a), a structure essential to its enzymatic activity. Beyond metabolism, the tetrameric form is known to facilitate binding to mRNAs, particularly to AU-rich elements in the 3’ untranslated region (UTR) of inflammatory cytokines like IFN-γ and TNF-α, indicating GAPDH’s role in post-transcriptional regulation of immune responses.^53^ Electrostatic potential mapping of the tetrameric GAPDH structure reveals an extended positively charged groove on the enzyme’s surface, located at the interface between adjacent subunits (Fig. 5a). This groove has been proposed as a potential RNA-binding site, and AlphaFold-based docking with a 20-nucleotide poly(A) RNA (20A-RNA) confirms that the RNA strand winds smoothly into this groove on both sides of the tetramer (Fig. 5a)^41,42^.

**Fig. 5.**
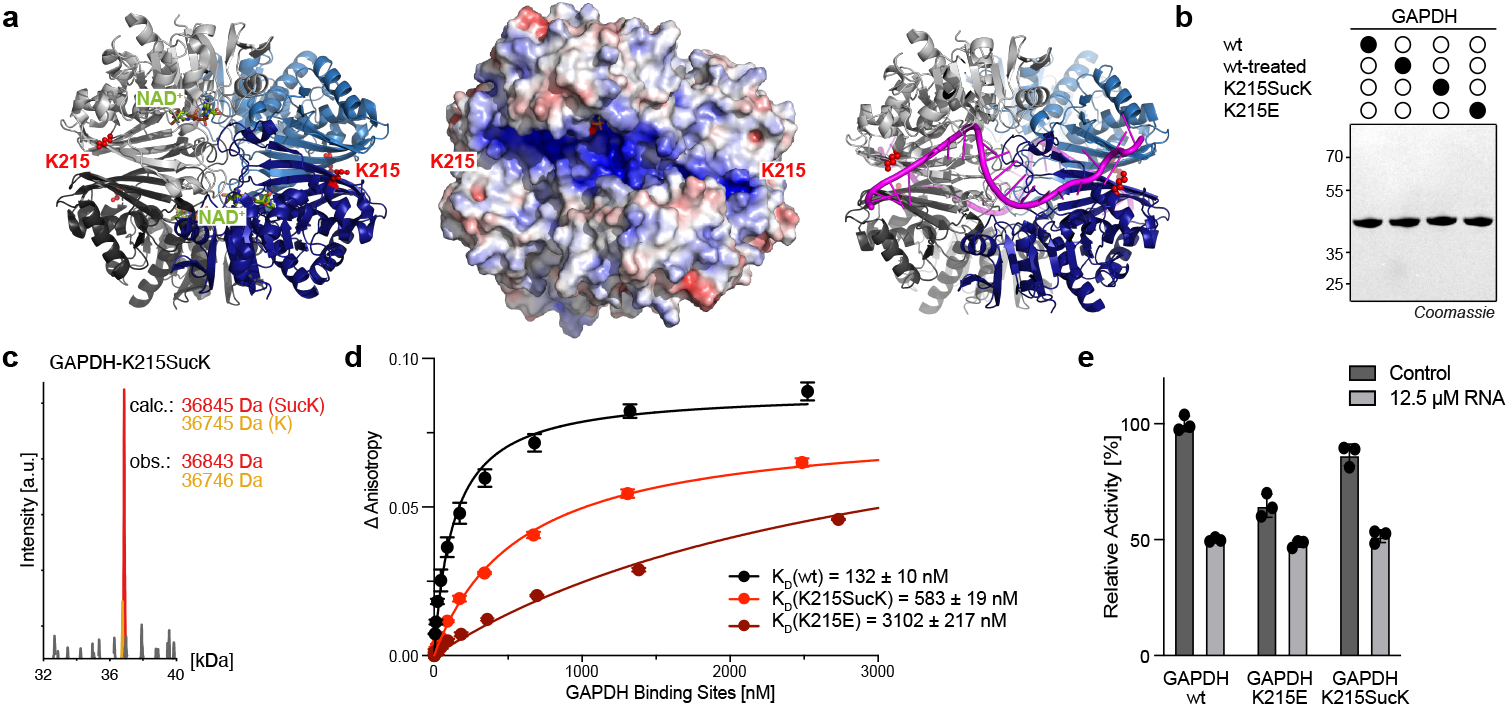
Succinylation of GAPDH at K215 modulates RNA binding. **a)** Left: Structure of the GAPDH tetramer with NAD^+^ and the K215 residue highlighted (red), (PDB: 1U8F)^54^. Middle: Electrostatic potential map of the GAPDH tetramer surface revealing a positively charged groove across the side of the tetramer. K215 residues are marked in red. Right: Alphafold prediction of GAPDH tetramer with two copies of 20A-RNA (pink), fitting into the positive groove on each side. K215 residues are highlighted in red^41,42^. Identical orientation of the GAPDH tetramer in all three presentations. **b)** SDS-PAGE analysis of GAPDH variants. **c)** Full-length mass spectrum of GAPDH-K215SucK installed via acylation of GAPDH-K215MeOK with **9i. d)** Fluorescence anisotropy results of binding event between fluorophore-labelled 20A-RNA and GAPDH variants. Change in anisotropy was plotted against the concentration of binding sites (two per tetramer) and fitted with single-site binding model to determine *K*_*D*_ values. The average values and standard deviations were calculated from three biologically independent experiments (*n* = 3). *K*_*D*_ values are given ± standard error. **e)** Comparison of relative activity of GAPDH variants without and in the presence of 20A-RNA. Activity of GAPDH-wt was normalized to 100%. Results from three biologically independent experiments (*n* = 3) are shown as black dots. Data was processed using GraphPad Prism 10 (GraphPad software).

Motivated by a recent study showing that lipopolysaccharide induced malonylation of a lysine residue within this groove regulated GAPDH binding of TNF-α mRNA, leading to its dissociation and thereby promoting translation and inflammation^55^, we set out to study effects of acylation introducing negative charge at this residue in in vitro settings. The respective residue, K215, sits at the ends of the positively charged groove and remains solvent-accessible in the tetrameric state. Proteomic data indicate that K215 can carry different PTMs, including acetylation, malonylation and succinylation.^56–58^ To probe the effect of introducing a negative charge at K215, we incorporated ncAA **13** at this position. In the presence of low concentrations of TCEP in the lysis buffer (1-2 mM), homogenous GAPDH-K215MeOK was obtained after standard affinity purification. Using our general workflow, we installed succinylation at K215. For comparison, we also expressed GAPDH-wt and a GAPDH-K215E mutant as a charge-mimic. The identity and purity of each variant was confirmed by SDS-PAGE and full-length MS analysis (Fig. 5b,c and Supplementary Fig. 14a).

To assess RNA binding affinities for the different GAPDH variants, we determined dissociation constants by fluorescence anisotropy using a fluorophore-labelled 20A-RNA probe. GAPDH-wt displayed strong RNA binding, with a K_D_ around 100 nM, whereas the K215E mutant exhibited a markedly reduced interaction, displaying a ca. 25-fold higher K_D_. The succinylated variant also showed diminished RNA binding, but to a lesser extent than GAPDH-K215E, with ca. 4-5-fold weaker affinity compared to GAPDH-wt (Fig. 5d, Supplementary Fig. 14b). This moderate reduction may be explained by the ability of a succinylated lysine side chain that is quite flexible to orient its negative charge away from the RNA backbone, thereby alleviating electrostatic repulsion. These findings point to a finely tuned interplay between the PTM state of K215 and RNA binding ability. RNA binding to the positively charged groove has been proposed to occlude or distort access to the active site and/or interfere with NAD^+^-binding.^48^ The AlphaFold model supports this, showing that the bound RNA sits directly in front of the NAD^+^-binding pocket, likely hindering proper co-factor association. To assess what effect RNA binding exerts on glycolytic activity, we measured the catalytic activity of each variant following the absorbance of NADH at 340 nm (Fig. 5e, Supplementary Fig. 15a,b). In the absence of 20A-RNA, GAPDH-K215SucK showed a modest reduction in activity (10-15%). For the glutamate variant the reduction was more pronounced, reaching ca. 60% of wt-activity. Addition of 20A-RNA reduced enzymatic activity for all variants to ca. 50% of wt-activity measured without RNA, with inhibitor effects more pronounced in variants with stronger RNA binding affinity. To see if succinylation at position K215 is regulated by SIRT5, we incubated GAPDH-K215SucK with this deacylase. However, we did not observe any notable desuccinylation (Supplementary Fig. 15c). Hence, we concluded that, at least in our *in vitro* setting, K215SucK is not reversed by SIRT5. It remains unknown whether the K215SucK modification is reversed by another eraser or the effect remains permanent.

## DISCUSSION and OUTLOOK

Lysine acylations constitute a diverse and rapidly expanding class of PTMs, yet many remain mechanistically and functionally unexplored due to the difficulty of generating homogenous, site-specifically modified target proteins^1-3,59^. Chemical protein synthesis is a powerful strategy to access proteins bearing diverse modifications, but becomes impractical for larger targets and is often hampered by refolding challenges.^60^ GCE offers an attractive alternative and has enabled the incorporation of many lysine acylations^4^. However, its efficiency can vary with the orthogonal aaRS/tRNA pair, and ncAAs can face limitations in cell permeability, chemical stability, or compatibility with the steric constraints of the aaRS active site, making the direct installation of bulky or charged acyl groups particularly challenging.

Here we introduce a modular ‚tag-and-modify’ platform for site-specific lysine acylation to circumvent these limitations. We designed and synthesized hydroxylamine-bearing ncAAs that upon incorporation into a POI can undergo a chemoselective amide bond-forming reaction with MIDA acylboronates, a relatively new class of stable acylboron compounds.^12,61^ To perform the described workflow, we rely on a single PylRS variant, which in the case of ncAA **13** leads to highly efficient amber suppression on various target proteins. We synthesized eleven different MIDA acylboronates and show the reliable, site-specific installation of a broad range of lysine acylations, including bulky and negatively charged species, directly on folded proteins under mild, aqueous conditions. The MIDA acylboronate-mediated acylation reaction leads to modified proteins that faithfully reproduce the structural and functional outcomes of endogenous PTMs. Furthermore, our approach enables access to acyl modifications that have remained inaccessible via direct genetic incorporation (e.g. malonylation, 4-oxononanoylation and itaconylation).

By requiring only a single PylRS/tRNA pair, our method lowers technical barriers and makes diverse acylations accessible to laboratories without specialized expertise in genetic code expansion. The synthetic routes to ncAAs **1** and **13** are straightforward and high yielding, ensuring ready access to ample material for GCE experiments. By contrast, MIDA acylboronates can be more challenging to obtain. As this compound class is relatively novel, only a limited number of synthetic procedures are available, often requiring greater synthetic expertise.^21,22,61^ Nevertheless, the broad range of MIDA acylboronates corresponding to naturally occurring acylations described here, together with the detailed procedures provided, substantially increases their accessibility.

With the MeOK-modified POI and the acylboronate for the desired modification at hand, their coupling reaction proceeds in a highly straightforward manner, at micromolar protein concentrations with an excess of acylboronate reagent. The reaction proved versatile across protein scaffolds and acyl types, and its high chemoselectivity typically yielded a single dominant product. A minor side product, which we frequently observed, can be attributed to an unmodified lysine residue, however this was usually present only in small amounts. As can be expected, the chemical environment of the hydroxylamine moiety in the POI influenced reaction rates, in rare cases product distribution. Overall, all tested proteins and sites gave very favourable outcomes, though buried residues inaccessible to small molecules will not be suitable to this modification chemistry. Furthermore, we only observed very sluggish yields for installing myristoylation (a C14 fatty acid), which we attribute to very poor solubility of the corresponding acylboronate in aqueous buffers, a limitation of the procedure to consider. The method’s versatility opens diverse future applications, from studying reader, writer and eraser specificities, to installing PTM analogues for direct crosslinking with interactors. Harnessing this modularity, in future work we aim to generate PTM libraries at single or multiple sites within a protein to probe optimal binders and study PTM cross-talk. In addition, future efforts will focus on enhancing reaction rate and selectivity with the long-term goal of enabling acylboronate-mediated acylations directly in cellulo. As the used MIDA acylboronates display a certain instability in aqueous environments, necessitating their use in 50-300-fold excess to achieve full conversion, future work will also be directed toward designing more stable, yet sufficiently reactive acylboronates. Such improvements would allow their use at near-equimolar ratios and better support in cellulo applications. Overall, the described acylation approach represents a significant advance for amide bond-forming reactions directly on folded proteins. We expect that this strategy will facilitate the study of lysine acylations, enabling exploration of PTMs that were previously out of reach. Furthermore, it may serve as inspiration for developing alternative chemoselective chemistries to access other elusive protein modifications.

## AUTHOR CONTRIBUTIONS

K.L. and T.A.N. envisioned the ‘tag-and-modify’-approach to lysine acylations. T.A.N. synthesized initial carbamoylhydroxylamines and performed attempts to genetically incorporate them into proteins. P.K. synthesized the MeOK ncAAs, evolved the corresponding pNZ-MeOKRS and conducted PylRS screens to identify oAZ-MeOKRS. P.K. expressed and purified Ub, SUMO2, IDH, histone H2B and GAPDH bearing site-specific MeOK. P.K. synthesized MIDA acylboronates, optimized reaction conditions with MeOK-bearing POIs and installed site-specific acylations. P.K. performed nitroreductase experiments, IDH activity assays, GAPDH assays and deacylation assays. P.K. and K.L. analysed the data, and K.L. supervised the study. K.L. and P.K. wrote the paper with input from T.A.N.

## Acknowledgements

This work was supported by funding from ETH Zurich and the European Research Council (ERC under the European Union’s Horizon 2020 research and innovation program, grant agreement no. 101003289–Ubl-tool to K.L.). We thank Dr. Maximilian Fottner for experimental guidance, in particular with protein purification, assistance in anisotropy experiments and general discussion. We also thank Lang group members for useful discussions and input. We thank Prof. Jeffrey Bode for insightful discussions.

