## Supplementary Information for "A Modular Genetic Code Expansion Approach to Site-Specific Lysine Acylations"

### Supplementary Figures 1 – 14

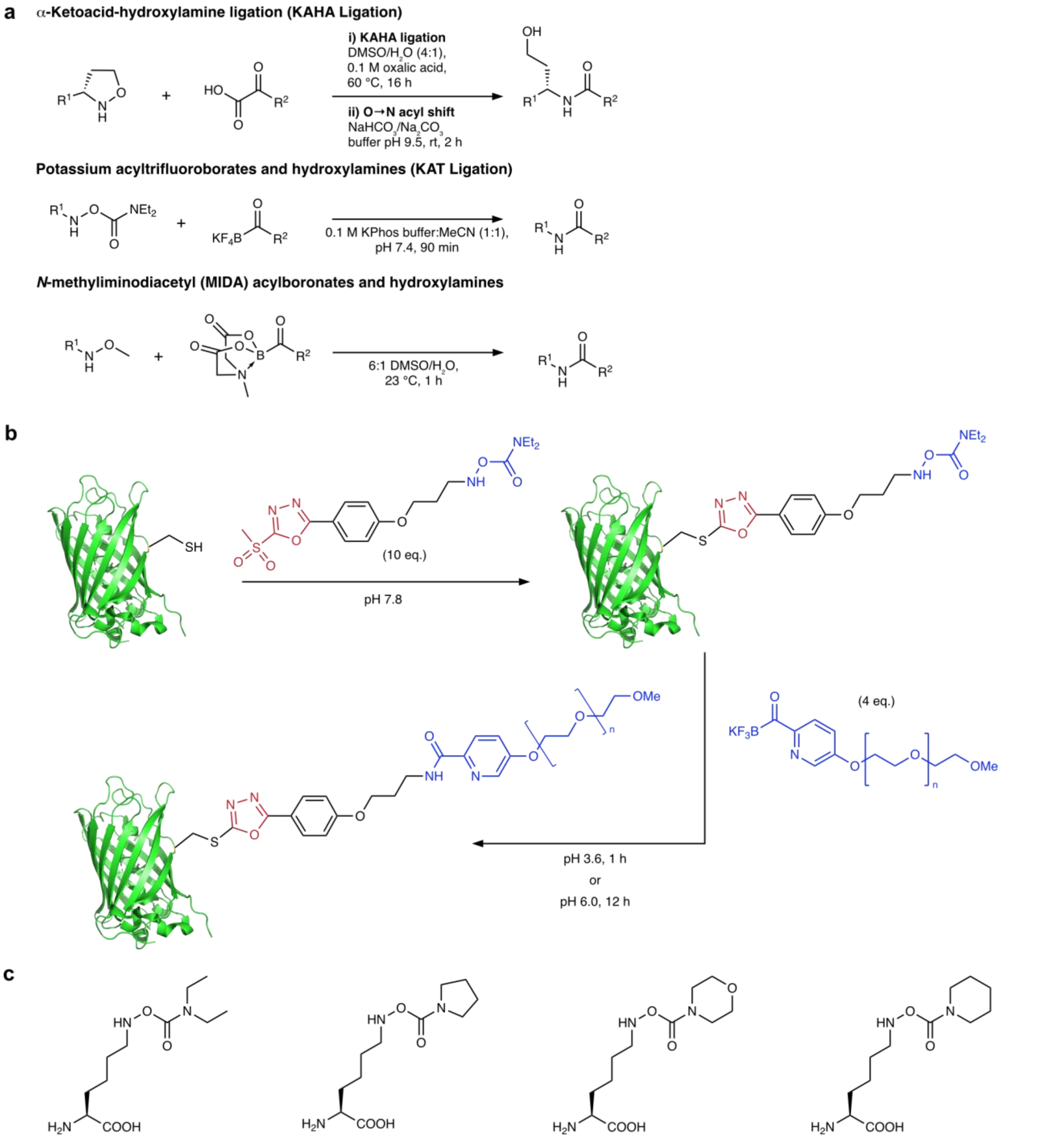

**Supplementary Figure 1 | Reaction of hydroxylamines with activated acyl donors and preliminary experiments**
**a)** Reactions of hydroxylamines with α-ketoacids[1], potassium acyltrifluoroborates[2] and *N*-methyliminodiacetyl (MIDA) acylboronates[3] under typical conditions. **b)** Modification of a super-folder green fluorescent protein (sfGFP) mutant (S147C) with a carbamoylhydroxylamine via a cysteine reactive moiety. Subsequently, a 2-pyridine-derived potassium acyltrifluoroboronate was used to PEGylate the protein.[4] **c)** Various carbamoylhydroxylamine-bearing ncAAs that were envisioned and synthesized in this study, but turned out not to be stable enough for incorporation into proteins via genetic code expansion.

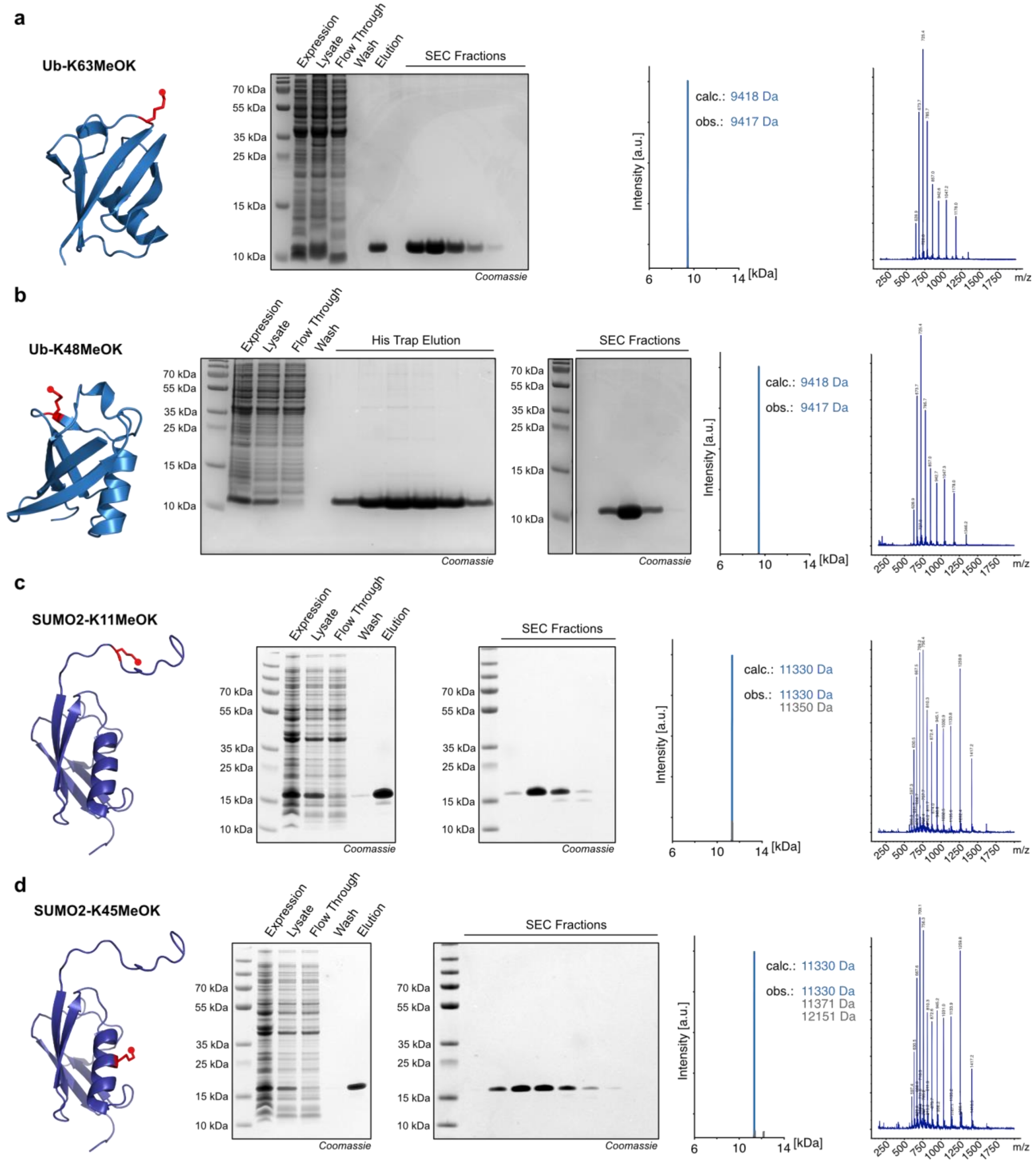

**Supplementary Figure 2 | Site-specific incorporation of MeOK into various proteins**
For each position, a graphical representation of the position at which the ncAA was installed is shown, together with the corresponding purification gels, the full-length MS analysis and the respective ionization pattern on which deconvolution was performed. Protein Data Bank files 1UBQ (ubiquitin)[5] and 2N1W (SUMO2) were used. **a)** Expression of Ub-K63MeOK. **b)** Expression of Ub-K48MeOK. **c)** Expression of SUMO2-K11MeOK. **d)** Expression of SUMO2-K45MeOK.

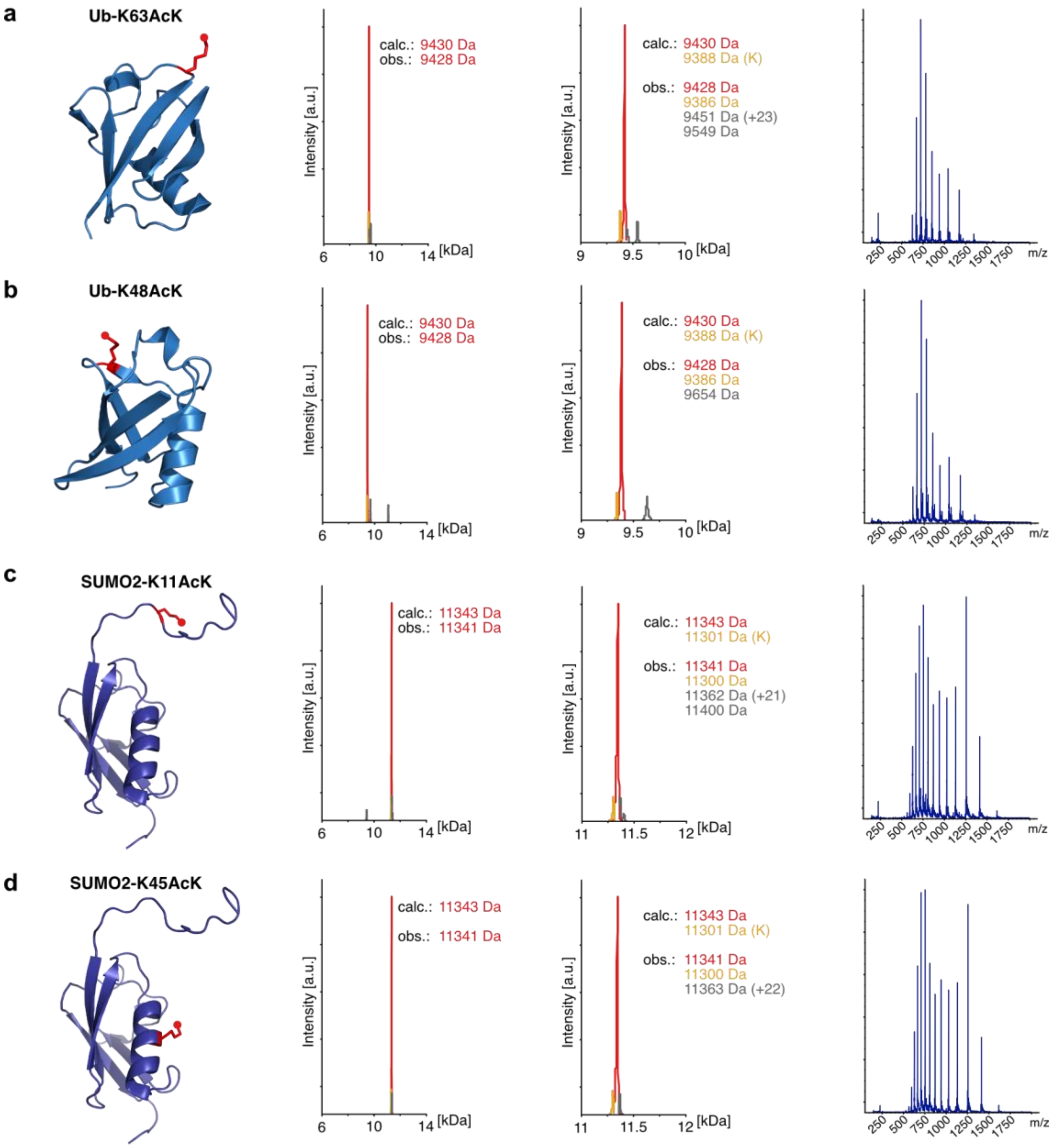

**Supplementary Figure 3 | Installation of acetylation on various positions of ubiquitin and SUMO2**
For each position, a graphical representation of the position at which the modification was installed, full-length MS analysis from 6 to 14 kDa as well as a zoomed version from 9 to 10 kDa and the corresponding ionisation pattern, which was used as basis for deconvolution are shown. Protein Data Bank files 1UBQ (ubiquitin)[5] and 2N1W (SUMO2) were used. Modifications were installed under the following conditions: 20 µM MeOK-modified protein, 2 mM MIDA acetylboronate **9a**, 50 mM citrate buffer pH 7.0, 2.5% DMSO, 23 °C, 3 hours. Peaks for MeOK-modified POIs are coloured in blue, acetylated POIs in red and wild-type POIs (bearing lysine) in yellow. **a)** Installation of acetylation on Ub-K63. **b)** Installation of acetylation on Ub-K48. **c)** Installation of acetylation on SUMO2-K11. **d)** Installation of acetylation on SUMO2-K45.

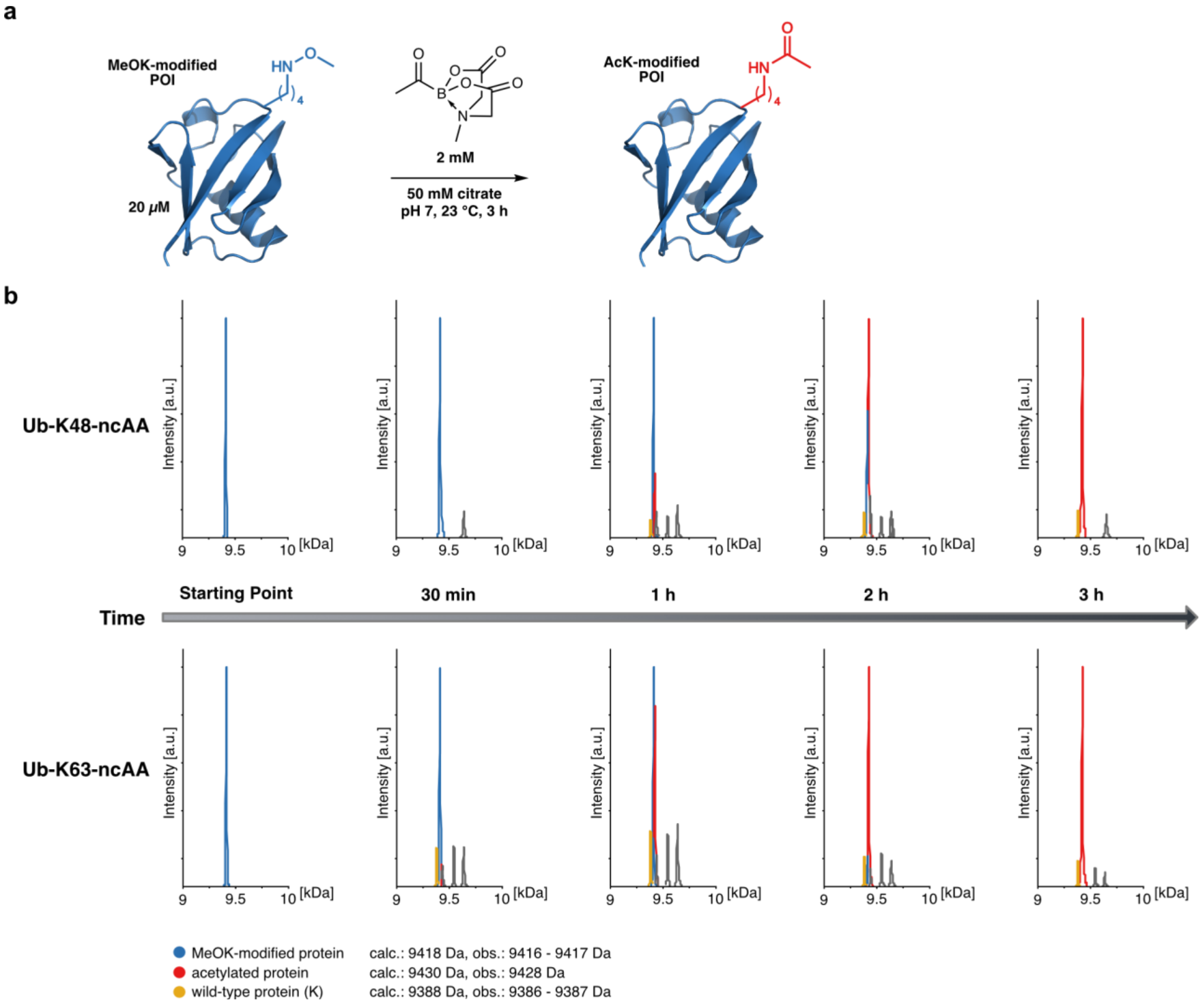

**Supplementary Figure 4 | Reaction monitoring for acetylation of Ub-K48MeOK and Ub-K63MeOK**
**a)** Schematic representation of the reaction performed on MeOK-bearing Ub with MIDA acetylboronate **9a** for accessing site-specifically acetylated Ub variants. Reaction conditions for the time courses depicted in b) are part of the scheme(PDB: 1UBQ)[5]. **b)** Time courses of the reaction of Ub-K48MeOK and Ub-K63MeOK with MIDA acetylboronate **9a**. The reactions were monitored by LC-MS. Observed masses are listed on the bottom.

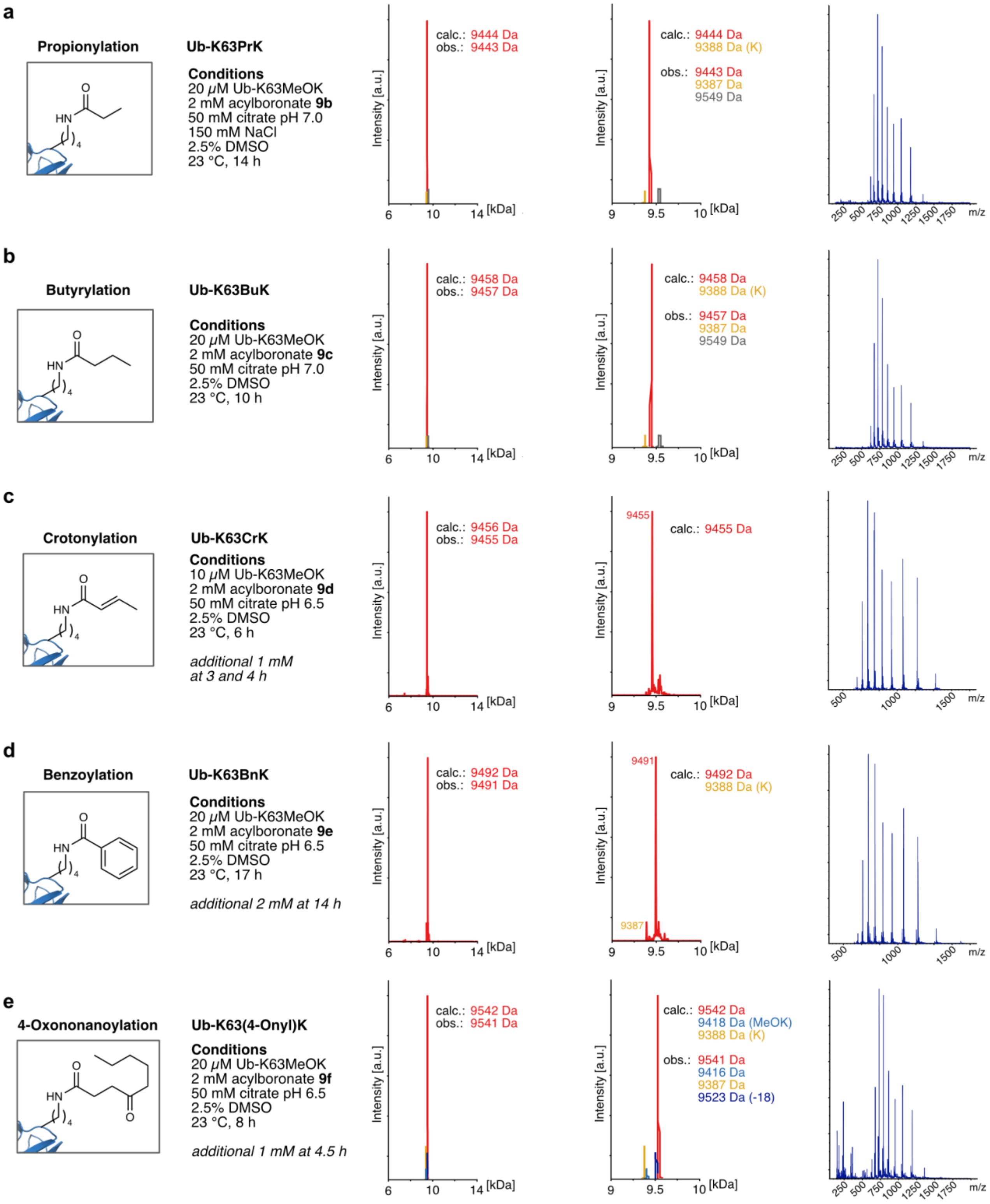

**Supplementary Figure 5 | Installation of various acylations 1**
Installation of lysine acylations using expressed Ub-K63MeOK and the corresponding acylboronates. All modifications were installed at position K63 on Ub. Respective conditions are listed next to the structural depiction of the acylation. Deconvoluted LC-MS analyses are displayed in the range of 6 to 14 kDa and 9 to 10 kDa (zoomed version). On the right, the obtained ionization pattern is displayed, which was used for deconvolution. Peaks for MeOK-modified Ub are coloured in blue, acylated Ub variants are depicted in red and wild-type Ub is shown in yellow. **a)** Installation of propionylation. **b)** Installation of butyrylation. **c)** Installation of crotonylation. **d)** Installation of benzoylation. **e)** Installation of 4-oxononanoylation.

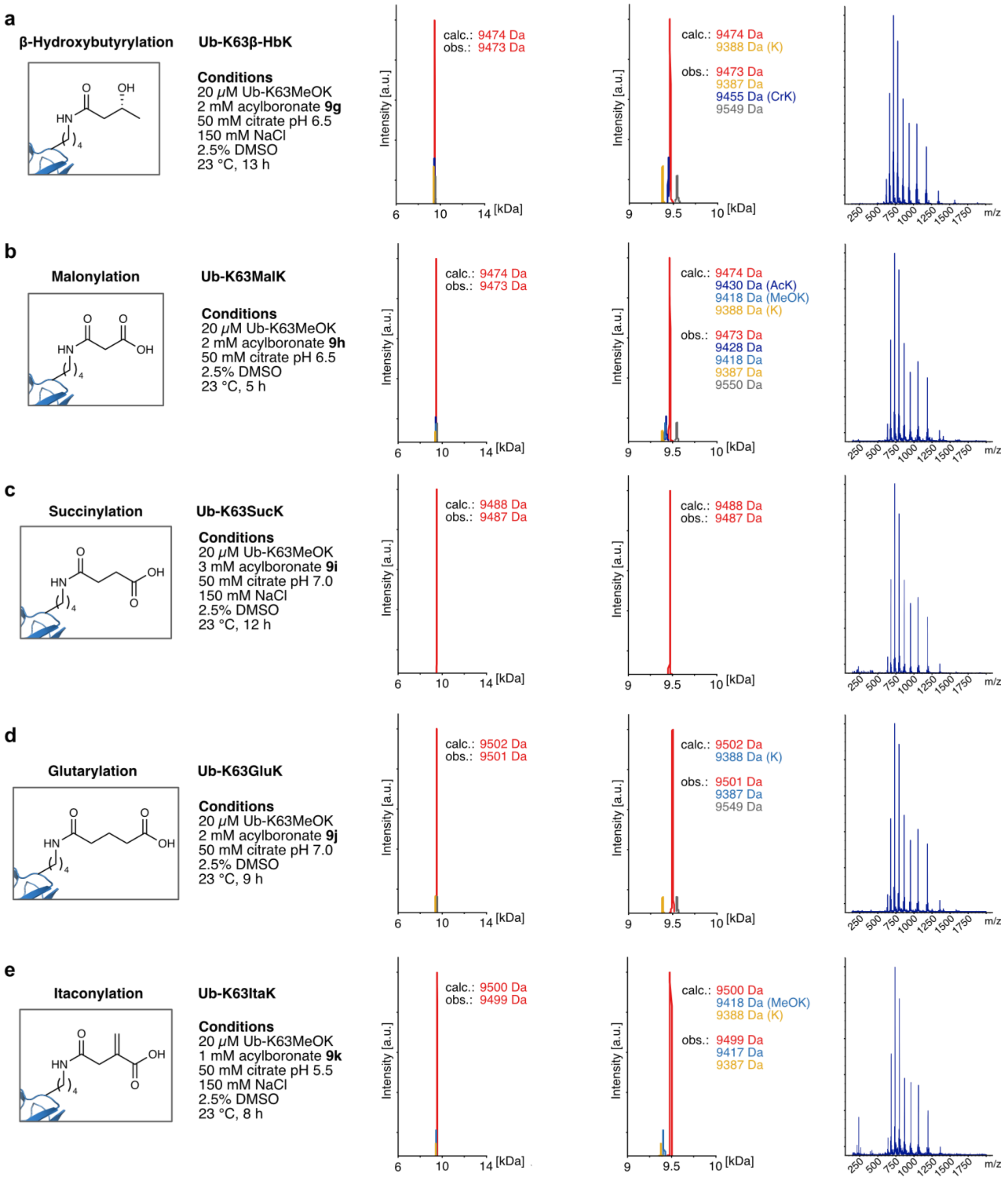

**Supplementary Figure 6 | Installation of various acylations 2**
Installation of lysine acylations using expressed Ub-K63MeOK and the corresponding acylboronates. All modifications were installed at position K63 on Ub. Respective conditions are listed next to the structural depiction of the acylation. Deconvoluted LC-MS analyses are displayed in the range of 6 to 14 kDa and 9 to 10 kDa (zoomed version). On the right, the obtained ionization pattern is displayed, which was used for deconvolution. Peaks for MeOK-modified Ub are coloured blue, acylated Ub variants are red and wild-type Ub is yellow. **a)** Installation of β-hydroxybutyrylation. **b)** Installation of malonylation. **c)** Installation of succinylation. **d)** Installation of glutarylation. **e)** Installation of itaconylation.

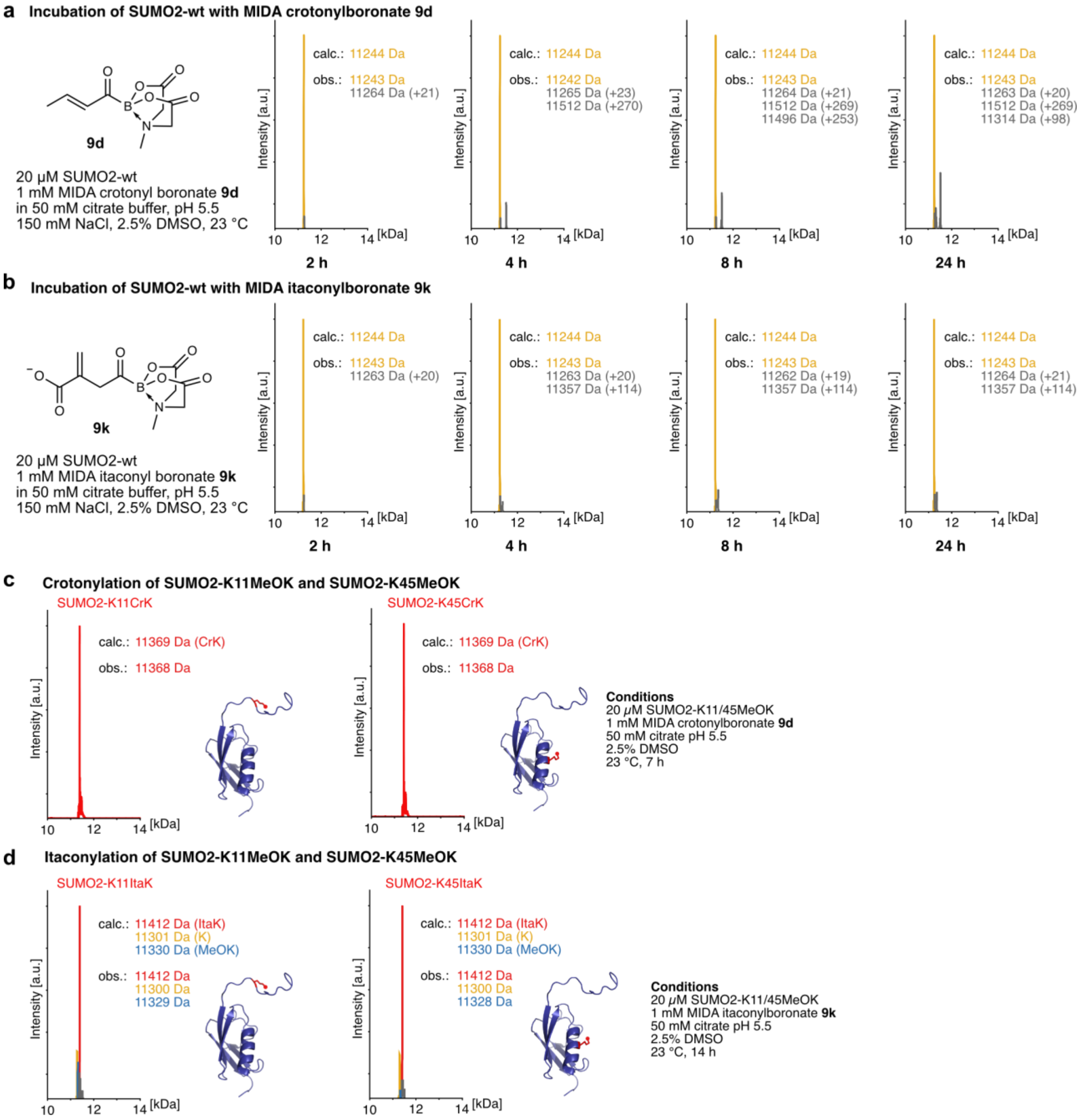

**Supplementary Figure 7 | Acylation with acylboronates containing electrophilic groups**

**a)** Incubation of SUMO2-wt with MIDA crotonylboronate **9d** to assess unspecific background modification. Over the course of 24 hours, the rise of a second peak can be observed. Its exact identity could however not be determined. Importantly, in the presence of MeOK-bearing protein a single dominant product peak was observed under the used conditions (see panel c). **b)** Incubation of SUMO2-wt with MIDA itaconylboronate **9k** to assess unspecific background modification. Over the course of 24 hours, no major unspecific reaction was observed. **c)** Installation of crotonylation at positions K11 and K45 of SUMO2 by reacting SUMO2-K11MeOK and SUMO2-K45MeOK via the displayed conditions. Full-length MS analyses show one fairly clean product peak for both SUMO2 variants. **d)** Installation of itaconylation at positions K11 and K45 of SUMO2 variants bearing MeOK at the corresponding positions via the displayed conditions. Full-length MS analyses show one dominant product peak with some side-product lacking the modification. Peaks for MeOK-modified SUMO2 are coloured blue, acylated SUMO2 variants are red and wild-type SUMO2 is yellow. Protein Data Bank 2N1W (SUMO2) was used.

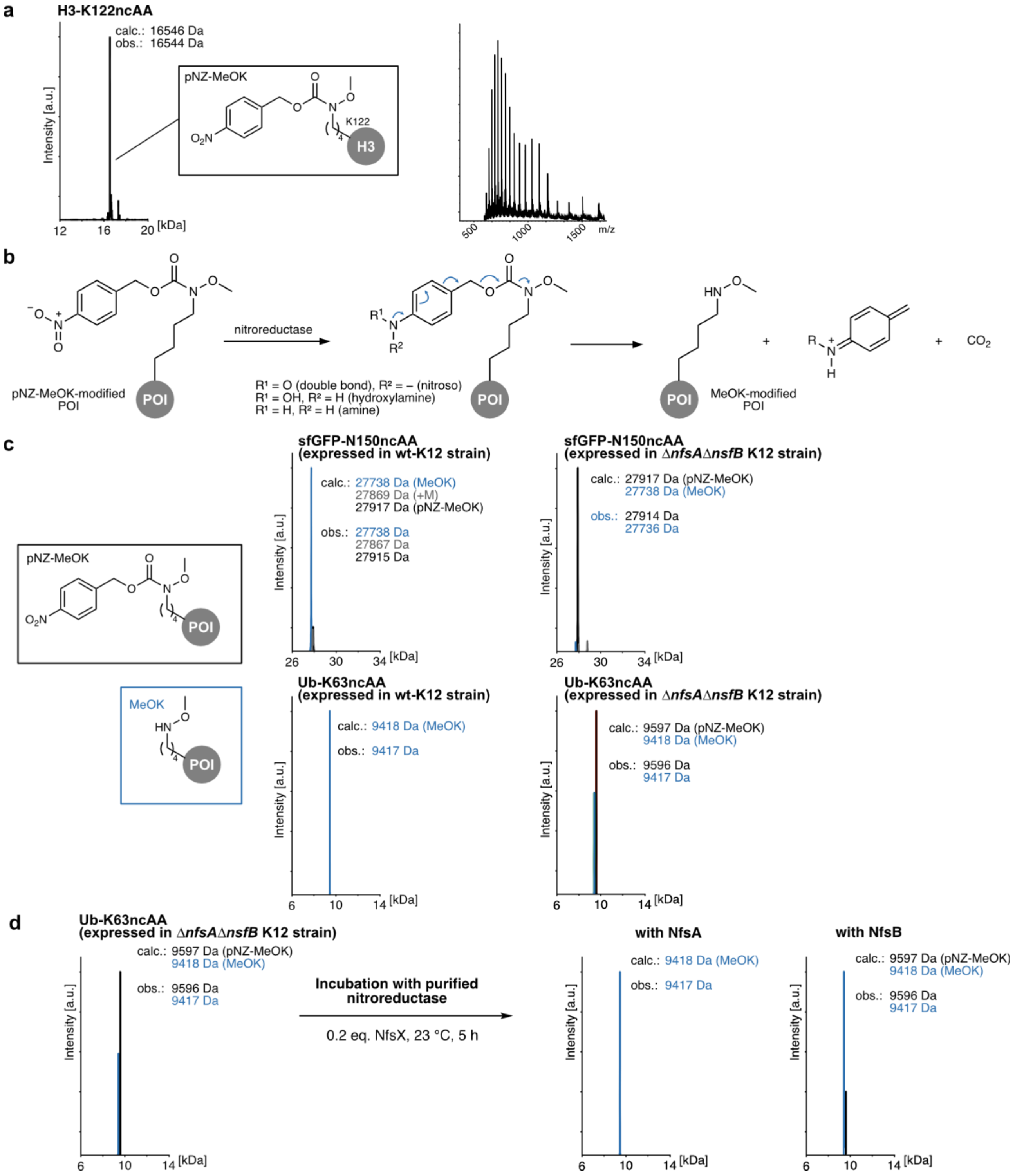

**Supplementary Figure 8 | Difficulties using nitroreductase-dependent decaging of pNZ-MeOK**

**a)** Left: Full-length mass spectrum of purified histone H3 (expressed from a H3-K122TAG plasmid in the presence of pNZ-MeOK) showing that the pNZ group is still intact after protein expression. Right: Ionization pattern on which deconvolution was performed to obtain the full-length mass spectrum. **b)** Proposed mechanism for on-protein reduction of pNZ-MeOK-bearing proteins by nitroreductases and liberation of MeOK-bearing protein.[6] **c)** Top: LCMS analysis of sfGFP-N150MeOK expressed in wt K12 cells or in the Δ*nfsA*Δ*nfsB* double knockout. In the latter, the pNZ protecting group remains almost completely intact. Bottom: LC-MS analysis of Ub-K63MeOK expressed in wt K12 cells or in the Δ*nfsA*Δ*nfsB* double knockout. In the latter, the pNZ protecting group is only partially unreduced. **d)** Incubation of Ub-K63MeOK expressed in the Δ*nfsA*Δ*nfsB* double knockout containing partially unreduced pNZ cage with expressed and purified *E. coli* nitroreductases. 20 µM Ub-K63pNZ-MeOK was incubated with 0.4 µM NfsA/B in 10 mM Tris pH 7.0, 100 mM NaCl, 2 mM NADH at 23 °C for 5 hours, then analysed by LCMS.

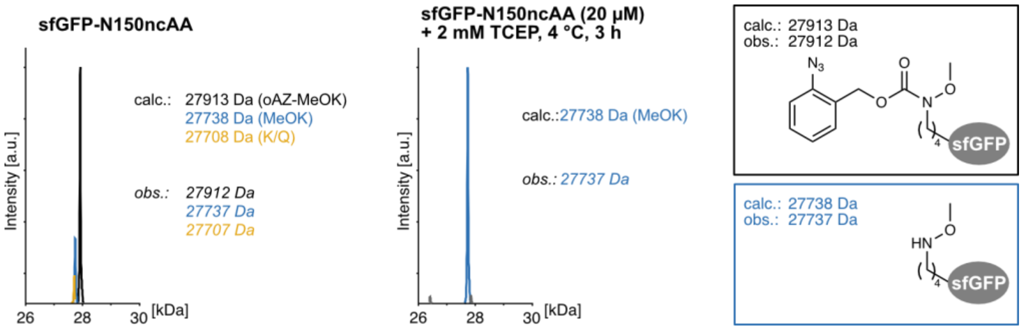

**Supplementary Figure 9 | Expression of MeOK-bearing sfGFP using oAZ-MeOK**

Left: Full-length mass spectrum of sfGFP expressed with amber suppression using oAZ-MeOK. After protein purification without using any reducing agents, the hydroxylamine is largely still protected with the oAZ group in place. Middle: Full-length mass spectrum showing that treatment of the purified protein with 2 mM TCEP for 3 hours at 4 °C results in complete removal of the oAZ group. Right: Depiction of the structures present at the targeted residue.

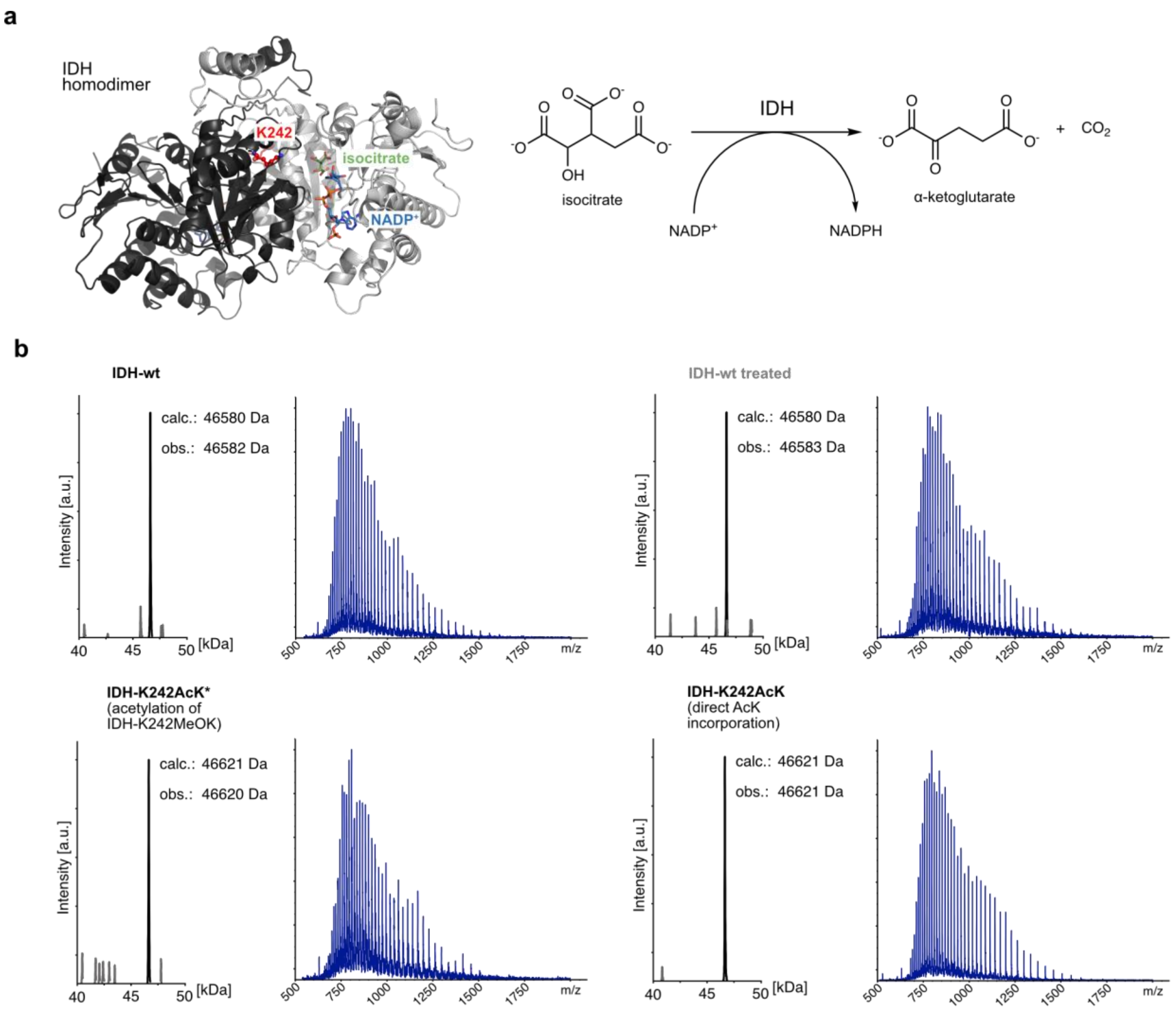

**Supplementary Figure 10 | Acetylation of IDH**

**a)** Left: Structure of the IDH homodimer (Protein Data Bank, 1AI2)[7]. The two subunits are coloured in black and grey. The targeted residue of the black subunit is highlighted in red, isocitrate in green and $NADP^+$ in blue. Right: Depiction of the reaction catalysed by IDH. **b)** Full-length mass spectra of IDH-wt, IDH-wt treated, IDH-K242AcK obtained through acetylation of incorporated MeOK and IDH-K242AcK obtained through direct incorporation of AcK using a previously published orthogonal synthetase.[8] IDH-wt treated received identical treatment as was used to install acetylation on MeOK. Ionization patterns on which deconvolutions were performed are displayed to the right of each mass spectrum.

**IDH-wt**

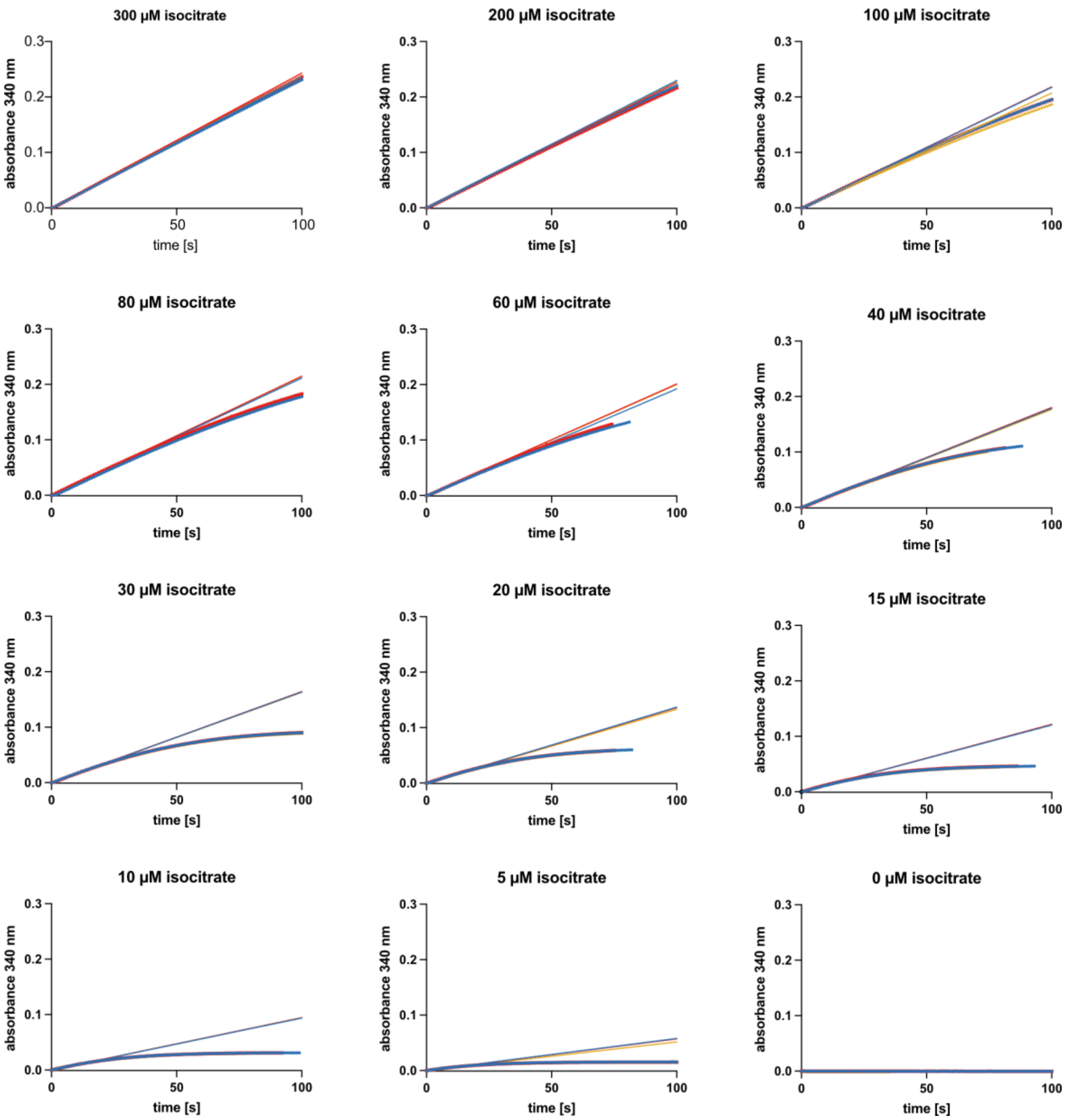

**Supplementary Figure 11 | IDH-wt kinetics**

Graphs displaying the linear regression of the initial reaction velocity of IDH-wt at substrate concentrations from 0 to 300 µM. NADPH absorbance was measured at 340 nM for ca. 100 seconds (dotted lines). The first 10 – 20 seconds (250 measurements/second) were used to perform a linear regression (solid lines) and determine the initial reaction velocity $v_0$. Each measurement and regression were performed three times (depicted in yellow, red and blue) from biologically independent reactions. Data was processed using GraphPad Prism 10 (GraphPad software).

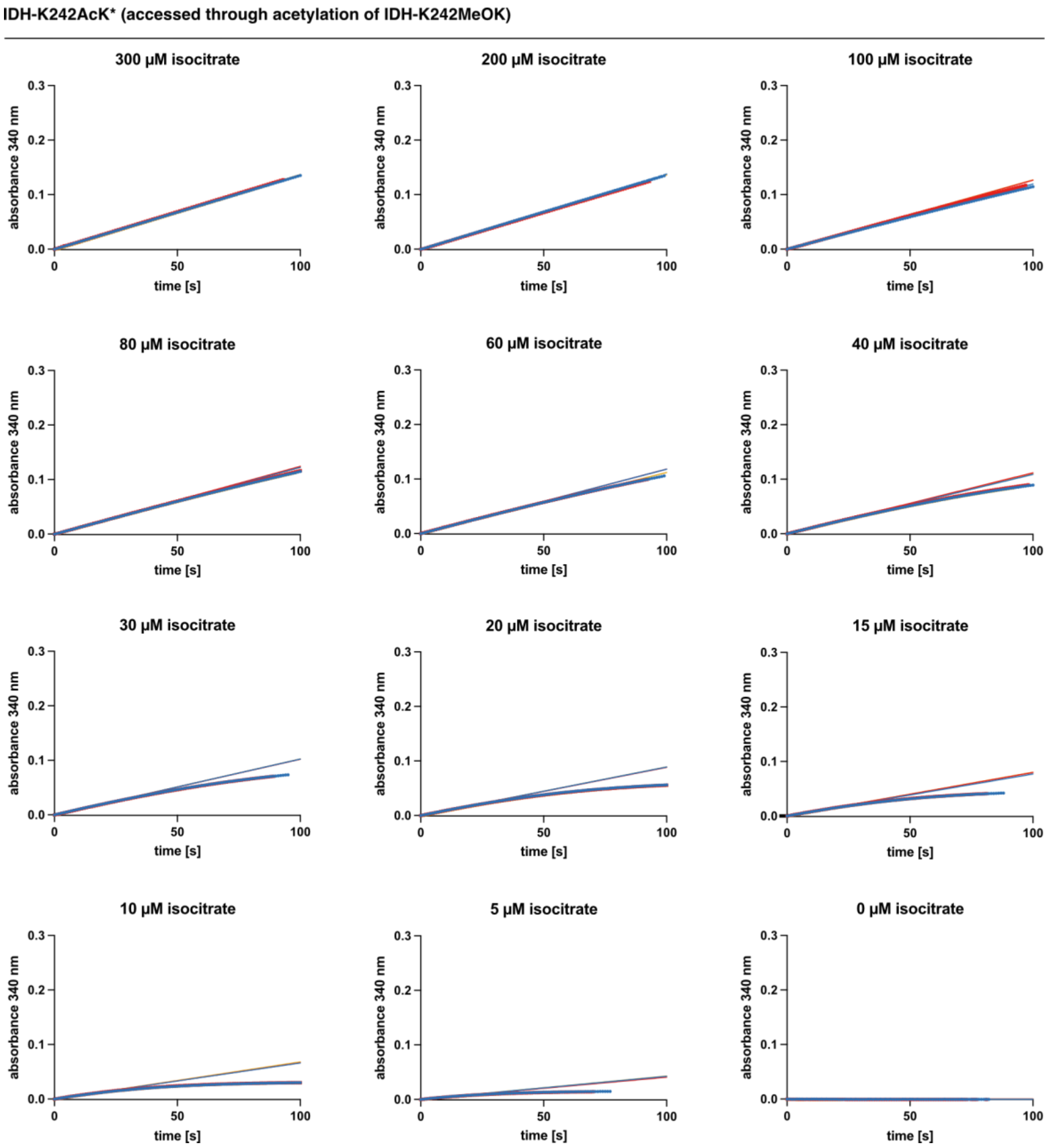

**Supplementary Figure 12 | IDH-K242AcK kinetics**

Graphs displaying the linear regression of the initial reaction velocity of IDH-K242AcK (accessed through acetylation of K242MeOK) at substrate concentrations from 0 to 300 µM. NADPH absorbance was measured at 340 nM for ca. 100 seconds (dotted lines). The first 10 – 20 seconds (250 measurements/second) were used to perform a linear regression (solid lines) and determine the initial reaction velocity $v_0$. Each measurement and regression were performed three times (depicted in yellow, red and blue) from biologically independent reactions. Data was processed using GraphPad Prism 10 (GraphPad software).

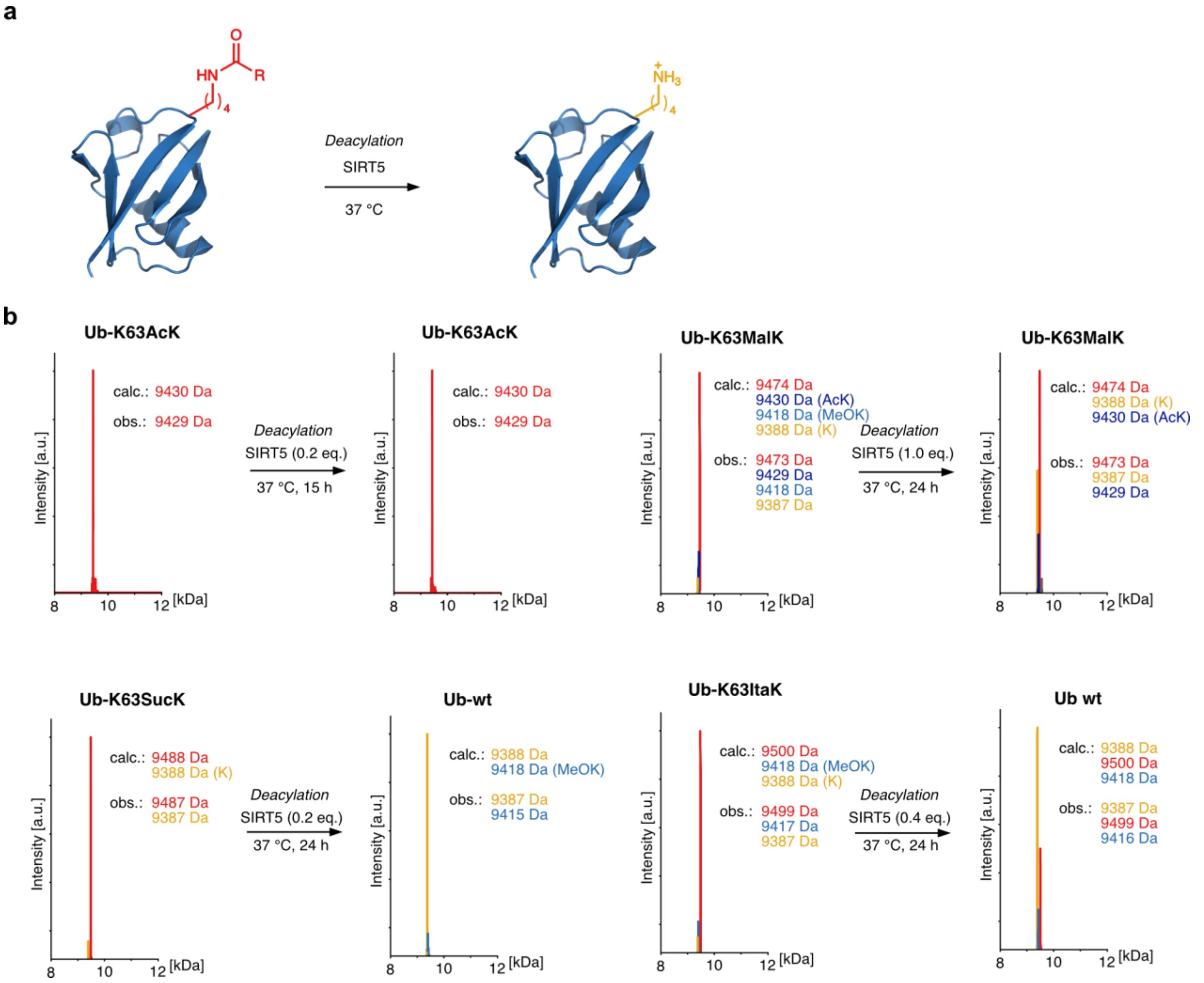

**Supplementary Figure 13 | Deacylation of Ub variants with SIRT5**

**a)** General reaction scheme displaying deacylation mediated by SIRT5. Protein Data Bank file 1UBQ was used[5]. **b)** Top left: Incubation of Ub-K63AcK with SIRT5. Top right: Incubation of Ub-K63MalK with SIRT5. Bottom left: Incubation of Ub-K63SucK with SIRT5. Bottom right: Incubation of Ub-K63ItaK with SIRT5. General reaction conditions were 10 µM acylated Ub, SIRT5 concentration and reaction time as indicated, at 37 °C in 50 mM Tris pH 7.4 at 37 °C, 100 mM NaCl, 5 mM $MgCl_2$, 1 mM DTT and 5 mM $NAD^+$. Ub-K63AcK was used at 20 µM with 4 µM SIRT5.

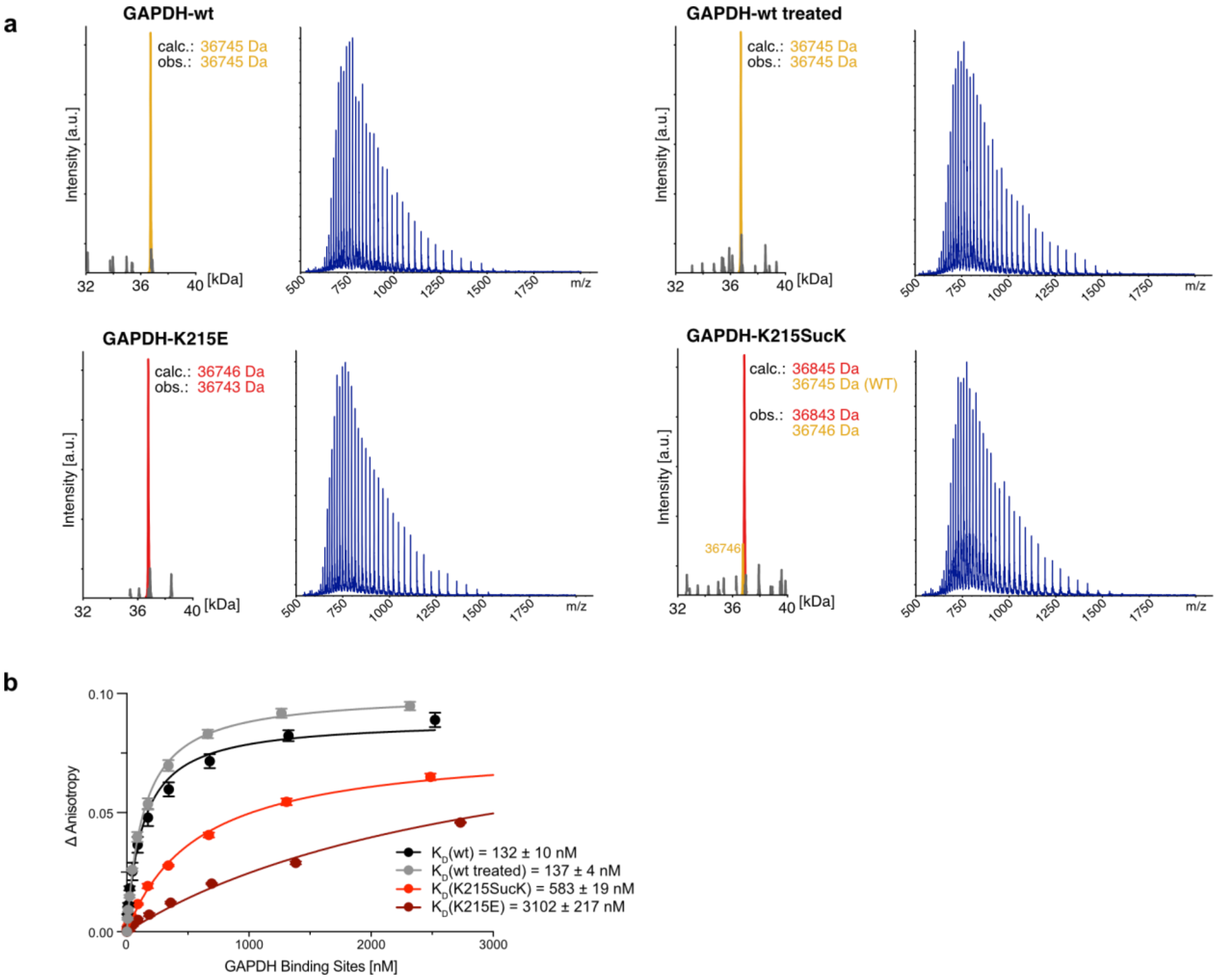

**Supplementary Figure 14 | GAPDH succinylation influences RNA binding 1**
**a)** Full-length mass spectra of GAPDH-wt, GAPDH-wt treated, GAPDH-K215E and GAPDH-K215SucK. Ionization patterns on which deconvolutions were performed are displayed to the right of each mass spectrum. GAPDH-wt treated underwent identical conditions as used for installation of SucK on GAPDH-K215MeOK. GAPDH-wt peaks are coloured yellow, acylated or glutamate mutant GAPDH peaks are displayed in red. **b)** Fluorescence anisotropy results of binding event between fluorophore-labelled 20A-RNA and GAPDH variants including GAPDH-wt treated. Change in anisotropy was plotted against the concentration of binding sites (two per tetramer) and fitted with a single-site binding model to determine $K_D$ values. The average values and standard deviations were calculated from three biologically independent experiments ($n$ = 3). $K_D$ values are given ± standard error. Data was processed using GraphPad Prism 10 (GraphPad software).

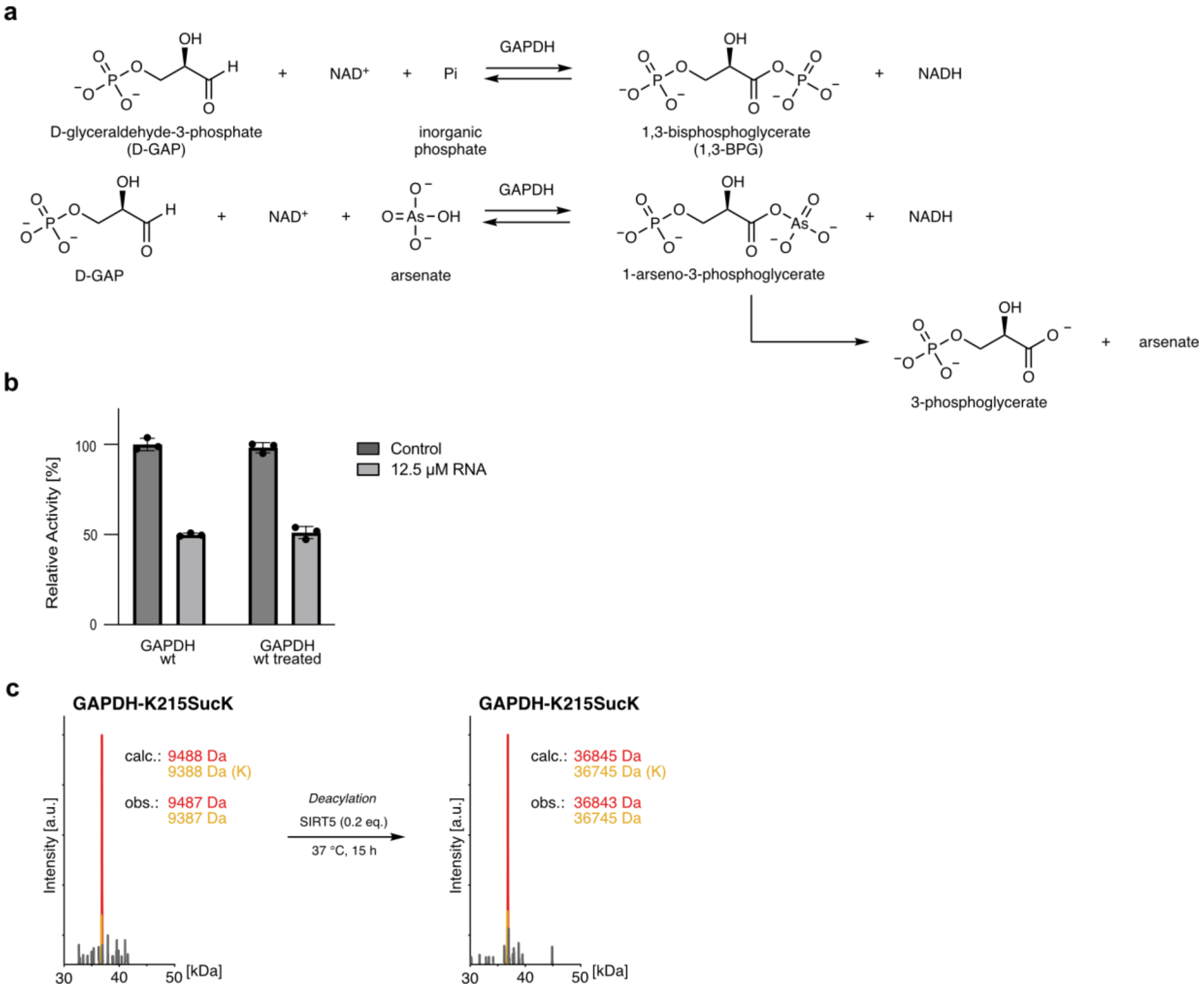

**Supplementary Figure 15 | GAPDH succinylation influences RNA binding 2**

**a)** Top: Depiction of the reaction catalysed by GAPDH, converting D-GAP to 1,3-BPG. Bottom: In the performed kinetic experiments, arsenate was used to render the reaction irreversible, as produced 1-arseno-3-phosphogylcerate rapidly hydrolyses in aqueous solution to 3-phosphoglycerate and hence allows investigation of the forward reaction (D-GAP to 1-arseno-3-phosphogylcerate). **b)** Comparison of relative activity of GAPDH-wt and GAPDH-wt treated in the activity assay displayed in Fig. 5e. Activity of GAPDH-wt was normalized to 100%. Results from three biologically independent experiments ($n$ = 3) are shown as black dots. **c)** Full-length mass spectra of GAPDH-K215SucK before and after incubation with deacylase SIRT5. No significant change was observed to the modified protein under the tested conditions. GAPDH-K215SucK peaks are displayed in red, GAPDH-wt peaks in yellow. Conditions: 20 µM GAPDH-K215SucK, 4 µM SIRT5, 50 mM Tris pH 7.4, 100 mM NaCl, 5 mM $MgCl_2$, 1 mM DTT and 5 mM $NAD^+$, 37 °C, 15 hours.

### Experimental procedures

#### 1 General methods: Plasmids and reagents

Codon optimized genes encoding Ubiquitin, SUMO2, Histone H2B (*mouse*), IDH (*E. coli*), NfsA (*E. coli*), NfsB (*E. coli*) and SIRT5 were purchased as DNA strings (Twist Bioscience). GAPDH (glyceraldehyde 3-phosphate dehydrogenase) vector was purchased from Addgene plasmid #83910. Genes were cloned into pBAD, pPylT, pETM11 or pET17 vectors via restriction cloning (see Supplementary Table S2). Point mutations were introduced via site-directed mutagenesis by PCR-driven overlap extension.[9] Oligonucleotide primers were designed using NEBBuilder and purchased from Microsynth. Extinction coefficients and molecular weight of proteins were calculated with ProtParam (https://web.expasy.org/protparam/). Amino acid sequences of all proteins are listed in section 1.3. RNA oligonucleotides were purchased from Integrated DNA Technologies (IDT).

All reaction solvents and chemical reagents were purchased from Sigma Aldrich, Acros Organics, TCI, Fisher Scientific, Fluorochem, BLDpharm, Apollo Scientific or Carl Roth and were used without further purification unless stated otherwise. Technical grade solvents (cyclohexane, hexane, ethyl acetate, pentane and diethyl ether) were obtained from the ETH solvent distribution facility and distilled before using for workup or chromatography. Bolt 4-12% Bis-Tris gradient gels (Invitrogen) or self-casted 15% SDS-PAGE gels were run (165 V for 43 min) on a Bolt™ Mini Gel Tank system (Invitrogen). Gels were stained with QuickBlue Protein Stain (LubioScience). PageRuler Prestained Plus Protein Ladder 10-250 kDa (ThermoFisher) was used as the protein marker. Protein and DNA concentrations were measured on a NanoPhotometer® NP60 (Implen). Ni Sepharose™ 6 Fast Flow was purchased from Cytiva.

Unless stated otherwise, antibiotics in bacterial cultures were used at the following concentrations: tetracycline (12.5 µg/mL), ampicillin (100 µg/mL), kanamycin (50 µg/mL) or chloramphenicol (50 µg/mL) for full strength (1x).

##### 1.1 RNA oligonucleotides for GAPDH fluorescence anisotropy and activity assay

Supplementary Table S1: RNA oligonucleotides for GAPDH fluorescence and activity assay

| Oligo | Modification | Sequence (5' → 3') |
|---|---|---|
| 1 | 5'-ATTO532 | AAAAA AAAAA AAAAA AAAAA |
| 2 | none | AAAAA AAAAA AAAAA AAAAA |

##### 1.2 Plasmids

Supplementary Table S2: Plasmids.

| Plasmid | Description |
|---|---|
| pBAD_sfGFP-V2P-H6 | sfGFP (superfolder green fluorescent protein) wild-type (wt) with V2P mutation for efficient methionine processing and C-terminal H6-tag under an arabinose promoter (ampicillin resistance). |
| pBAD_sfGFP-V2P-N150TAG-H6 | sfGFP with V2P mutation for efficient methionine processing, N150TAG and C-terminal H6-tag under an arabinose promoter (ampicillin resistance). |
| pPylT_Ub-K27TAG-H6 | Ubiquitin-K27TAG mutant with C-terminal H6-tag under an arabinose promoter and a *M. alvus* PylT copy under a constitutive promoter (tetracycline resistance). |
| pBAD_Ub-K27TAG-H6 | Ubiquitin-K27TAG mutant with C-terminal H6-tag under an arabinose promoter (ampicillin resistance). |
| pPylT_Ub-K48TAG-H6 | Ubiquitin-K48TAG mutant with C-terminal H6-tag under an arabinose promoter and a *M. alvus* PylT copy under a constitutive promoter (tetracycline resistance). |
| pPylT_Ub-K63TAG-H6 | Ubiquitin-K63TAG mutant with C-terminal H6-tag under an arabinose promoter and a *M. alvus* PylT copy under a constitutive promoter (tetracycline resistance). |

| | |
|---|---|
| pPylT_SUMO2-K11TAG-H6 | SUMO2-K11TAG mutant with C-terminal H6-tag under an arabinose promoter and a *M. alvus* PylT copy under a constitutive promoter (tetracycline resistance). |
| pPylT_SUMO2-K45TAG-H6 | SUMO2-K45TAG mutant with C-terminal H6-tag under an arabinose promoter and a *M. alvus* PylT copy under a constitutive promoter (tetracycline resistance). |
| pBAD_Histone_H2B-K58TAG-H6 | *Mouse* Histone H2B with K58TAG mutation, C-terminal H6-tag and GS linker under an arabinose promoter (ampicillin resistance). |
| pPylT_IDH-wt-H6 | *E. coli* IDH (isocitrate dehydrogenase) wt with C-terminal H6-tag under an arabinose promoter and a *M. alvus* PylT copy under a constitutive promoter (tetracycline resistance). |
| pBAD_IDH-K242TAG-H6 | *E. coli* IDH-K242TAG with C-terminal H6-tag under an arabinose promoter (ampicillin resistance). |
| pPylT_IDH-K242TAG-H6_Mb | *E. coli* IDH (isocitrate dehydrogenase) K242TAG mutant with C-terminal H6-tag under an arabinose promoter and a *M. barkeri* PylT copy under a constitutive promoter (tetracycline resistance). |
| pPylT_GAPDH-wt-H6 | *Human* GAPDH (glyceraldehyde 3-phosphate dehydrogenase) wt with C-terminal H6-tag under an arabinose promoter and a *M. alvus* PylT copy under a constitutive promoter (tetracycline resistance). |
| pPylT_GAPDH-K215E-H6 | *Human* GAPDH-K215E mutant with C-terminal H6-tag under an arabinose promoter and a *M. barkeri* PylT copy under a constitutive promoter (tetracycline resistance). |
| pBAD_GAPDH-K215TAG-H6 | *Human* GAPDH-K215TAG mutant with C-terminal H6-tag under an arabinose promoter (ampicillin resistance). |
| pETM11_H6-TEV-NfsA | *E. coli* NfsA wt with N-terminal H6-tag and TEV cleavage site under a T7 promoter (kanamycin resistance). |
| pETM11_H6-TEV-NfsB | *E. coli* NfsB wt with N-terminal H6-tag and TEV cleavage site under a T7 promoter (kanamycin resistance). |
| pET17_H6-TEV-SIRT5 | H6-TEV-SIRT5 (Sirtuin 5) without a mitochondrial localization signal (MLS) under a T7 promoter (ampicillin resistance). |
| pBK_pNZ-MeOKRS | *M. alvus* "pNZ-MeOKRS" PylRS under a constitutive GlnS promoter (ampicillin resistance). |
| pEVOL_oAZ-MeOKRS_PylT | *M. mazei* "oAZ-MeOKRS" PylRS under a constitutive GlnS and an arabinose inducible promoter with a pyrrolysyl-tRNA (PylT) copy under a constitutive promoter (chloramphenicol resistance). |
| pBK_AcKRS | *M. barkeri* PylRS mutant known to incorporate AcK under a constitutive GlnS promoter (ampicillin resistance). |
| pREP_CAT-D111TAG_PylT | Chloramphenicol acetyltransferase (CAT) with a D111TAG mutation under a constitutive promoter and *M. alvus* PylT (tetracycline resistance). |
| pYOBB2_Barnase-Q3TAG-D45TAG_PylT | Barnase with Q3TAG and D45TAG mutations under an arabinose inducible promoter and *M. alvus* PylT (chloramphenicol resistance). |
| pREP_CAT-D111TAG_sfGFP-N150TAG-His6_PylT | CAT with a D111TAG mutation under a constitutive promoter, sfGFP with a N150TAG mutation and C-terminal H6-tag and *M. alvus* PylT (tetracycline resistance). |

##### 1.3 Amino acid sequences of proteins

sfGFP-V2P-H6
MPSKGEELFTGVVPILVELDGDVNGHKFSVRGEGEGDATNGKLTLKFICTTGKLPVPWPTLVTTLTYGVQCFS
RYPDHMKRHDFFKSAMPEGYVQERTISFKDDGTYKTRAEVKFEGDTLVNRIELKGIDFKEDGNILGHKLEYNF
NSHNVYITADKQKNGIKANFKIRHNVEDGSVQLADHYQQNTPIGDGPVLLPDNHYLSTQSVLSKDPNEKRDH
MVLLEFVTAAGITHGMDELYKGSHHHHHH*

sfGFP-V2P-N150TAG-H6
MPSKGEELFTGVVPILVELDGDVNGHKFSVRGEGEGDATNGKLTLKFICTTGKLPVPWPTLVTTLTYGVQCFS
RYPDHMKRHDFFKSAMPEGYVQERTISFKDDGTYKTRAEVKFEGDTLVNRIELKGIDFKEDGNILGHKLEYNF
NSH*VYITADKQKNGIKANFKIRHNVEDGSVQLADHYQQNTPIGDGPVLLPDNHYLSTQSVLSKDPNEKRDH
MVLLEFVTAAGITHGMDELYKGSHHHHHH*

Ub-K27TAG-H6
MQIFVKTLTGKTITLEVEPSDTIENV*AKIQDKEGIPPDQQRLIFAGKQLEDGRTLSDYNIQKESTLHLVLRLRGG
HHHHHH*

Ub-K48TAG-H6
MQIFVKTLTGKTITLEVEPSDTIENVKAKIQDKEGIPPDQQRLIFAG*QLEDGRTLSDYNIQKESTLHLVLRLRGG
HHHHHH*

Ub-K63TAG-H6
MQIFVKTLTGKTITLEVEPSDTIENVKAKIQDKEGIPPDQQRLIFAGKQLEDGRTLSDYNIQ*ESTLHLVLRLRGG
HHHHHH*

SUMO2-K11TAG-H6
MADEKPKEGV*TENNDHINLKVAGQDGSVVQFKIKRHTPLSKLMKAYCERQGLSMRQIRFRFDGQPINETDTP
AQLEMEDEDTIDVFQQQTGGHHHHHH*

SUMO2-K45TAG-H6
MADEKPKEGVKTENNDHINLKVAGQDGSVVQFKIKRHTPLSKLM*AYCERQGLSMRQIRFRFDGQPINETDTP
AQLEMEDEDTIDVFQQQTGGHHHHHH*

Histone_H2B-K58TAG-H6
MPEPSKSAPAPKKGSKKAISKAQKKDGKKRKRSRKESYSVYVYKVLKQVHPDTGISS*AMGIMNSFVNDIFERI
ASEASRLAHYNKRSTITSREIQTAVRLLLPGELAKHAVSEGTKAVTKYTSSKGSHHHHHH*

IDH-wt-H6
MESKVVVPAQGKKITLQNGKLNVPENPIIPYIEGDGIGVDVTPAMLKVVDAAVEKAYKGERKISWMEIYTGEK
STQVYGQDVWLPAETLDLIREYRVAIKGPLTTPVGGGIRSLNVALRQELDLYICLRPVRYYQGTPSPVKHPELT
DMVIFRENSEDIYAGIEWKADSADAEKVIKFLREEMGVKKIRFPEHCGIGIKPCSEEGTKRLVRAAIEYAIANDR
DSVTLVHKGNIMKFTEGAFKDWGYQLAREEFGGELIDGGPWLKVKNPNTGKEIVIKDVIADAFLQQILLRPAE
YDVIACMNLNGDYISDALAAQVGGIGIAPGANIGDECALFEATHGTAPKYAGQDKVNPGSIILSAEMMLRHM
GWTEAADLIVKGMEGAINAKTVTYDFERLMDGAKLLKCSEFGDAIIENMHHHHHH*

IDH-K242TAG-H6
MESKVVVPAQGKKITLQNGKLNVPENPIIPYIEGDGIGVDVTPAMLKVVDAAVEKAYKGERKISWMEIYTGEK
STQVYGQDVWLPAETLDLIREYRVAIKGPLTTPVGGGIRSLNVALRQELDLYICLRPVRYYQGTPSPVKHPELT
DMVIFRENSEDIYAGIEWKADSADAEKVIKFLREEMGVKKIRFPEHCGIGIKPCSEEGTKRLVRAAIEYAIANDR
DSVTLVHKGNIMKFTEGAF*DWGYQLAREEFGGELIDGGPWLKVKNPNTGKEIVIKDVIADAFLQQILLRPAE
YDVIACMNLNGDYISDALAAQVGGIGIAPGANIGDECALFEATHGTAPKYAGQDKVNPGSIILSAEMMLRHM
GWTEAADLIVKGMEGAINAKTVTYDFERLMDGAKLLKCSEFGDAIIENMHHHHHH*

GAPDH-wt-H6
MGKVKVGVNGFGRIGRLVTRAAFNSGKVDIVAINDPFIDLNYMVYMFQYDSTHGKFHGTVKAENGKLVING
NPITIFQERDPSKIKWGDAGAEYVVESTGVFTTMEKAGAHLQGGAKRVIISAPSADAPMFVMGVNHEKYDNSL
KIISNASCTTNCLAPLAKVIHDNFGIVEGLMTTVHAITATQKTVDGPSGKLWRDGRGALQNIIPASTGAAKAVG
KVIPELNGKLTGMAFRVPTANVSVVDLTCRLEKPAKYDDIKKVVKQASEGPLKGILGYTEHQVVSSDFNSDTH
SSTFDAGAGIALNDHFVKLISWYDNEFGYSNRVVDLMAHMASKEHHHHHH*

GAPDH-K215E-H6
MGKVKVGVNGFGRIGRLVTRAAFNSGKVDIVAINDPFIDLNYMVYMFQYDSTHGKFHGTVKAENGKLVING
NPITIFQERDPSKIKWGDAGAEYVVESTGVFTTMEKAGAHLQGGAKRVIISAPSADAPMFVMGVNHEKYDNSL
KIISNASCTTNCLAPLAKVIHDNFGIVEGLMTTVHAITATQKTVDGPSGKLWRDGRGALQNIIPASTGAAEAVG
KVIPELNGKLTGMAFRVPTANVSVVDLTCRLEKPAKYDDIKKVVKQASEGPLKGILGYTEHQVVSSDFNSDTH
SSTFDAGAGIALNDHFVKLISWYDNEFGYSNRVVDLMAHMASKEHHHHHH*

GAPDH-K215TAG-H6
MGKVKVGVNGFGRIGRLVTRAAFNSGKVDIVAINDPFIDLNYMVYMFQYDSTHGKFHGTVKAENGKLVING
NPITIFQERDPSKIKWGDAGAEYVVESTGVFTTMEKAGAHLQGGAKRVIISAPSADAPMFVMGVNHEKYDNSL
KIISNASCTTNCLAPLAKVIHDNFGIVEGLMTTVHAITATQKTVDGPSGKLWRDGRGALQNIIPASTGAA*AVG
KVIPELNGKLTGMAFRVPTANVSVVDLTCRLEKPAKYDDIKKVVKQASEGPLKGILGYTEHQVVSSDFNSDTH
SSTFDAGAGIALNDHFVKLISWYDNEFGYSNRVVDLMAHMASKEHHHHHH*

H6-TEV-NfsA
MKHHHHHHPMSDYDIPTTENLYFQGAMAMTPTIELICGHRSIRHFTDEPISEAQREAIINSARATSSSSFLQCSSII
RITDKALREELVTLTGGQKHVAQAAEFWVFCADFNRHLQICPDAQLGLAEQLLLGVVDTAMMAQNALIAAES
LGLGGVYIGGLRNNIEAVTKLLKLPQHVLPLFGLCLGWPADNPDLKPRLPASILVHENSYQPLDKGALAQYDE
QLAEYYLTRGSNNRRDTWSDHIRRTIIKESRPFILDYLHKQGWATR*

H6-TEV-NfsB
MKHHHHHHPMSDYDIPTTENLYFQGAMDIISVALKRHSTKAFDASKKLTPEQAEQIKTLLQYSPSSTNSQPWH
FIVASTEEGKARVAKSAAGNYVFNERKMLDASHVVVFCAKTAMDDVWLKLVVDQEDADGRFATPEAKAAN
DKGRKFFADMHRKDLHDDAEWMAKQVYLNVGNFLLGVAALGLDAVPIEGFDAAILDAEFGLKEKGYTSLV
VVPVGHHSVEDFNATLPKSRLPQNITLTEV*

H6-TEV-SIRT5
MHHHHHHGGENLYFQGSSSMADFRKFFAKAKHIVIISGAGVSAESGVPTFRGAGGYWRKWQAQDLATPLAF
AHNPSRVWEFYHYRREVMGSKEPNAGHRAIAECETRLGKQGRRVVVITQNIDELHRKAGTKNLLEIHGSLFKT
RCTSCGVVAENYKSPICPALSGKGAPEPGTQDASIPVEKLPRCEEAGCGGLLRPHVVWFGENLDPAILEEVDRE
LAHCDLCLVVGTSSVVYPAAMFAPQVAARGVPVAEFNTETTPATNRFRFHFQGPCGTTLPEALACHENETVS*

pNZ-MeOKRS
MTVKYTDAQIQRLREYGNGTYEQKVFEDLASRDAAFSKEMSVASTDNEKKIKGMIANPSRHGLTQLMNDIAD
ALVAEGFIEVRTPIFISKDALARMTITEDKPLFKQVFWIDEKRALRPMLAPNLWSVCRDLRDHTDGPVKIFEMG
SCFRKESHSGMHLEEFTMLNLVDMGPRGDATEVLKNYISVVMKAAGLPDYDLVQEESDVYKETIDVEINGQE
VCSAAVGPSGLDAAHDVHEPWSGAGFGLERLLTIREKYSTVKKGGASISYLNGAKIN*

oAZ-MeOKRS
MDKKPLNTLISATGLWMSRTGTIHKIKHHEVSRSKIYIEMACGDHLVVNNSRSSRTARALRHHKYRKTCKRCR
VSDEDLNKFLTKANEDQTSVKVKVVSAPTRTKKAMPKSVARAPKPLENTEAAQAQPSGSKFSPAIPVSTQESV
SVPASVSTSISSISTGATASALVKGNTNPITSMSAPVQASAPALTKSQTDRLEVLLNPKDEISLNSGKPFRELESE
LLSRRKKDLQQIYAEERENYLGKLEREITRFFVDRGFLEIKSPILIPLEYIERMGIDNDTELSKQIFRVDKNFCLRP
MLAPNLANYLRKLDRALPDPIKIFEIGPCYRKESDGKEHLEEFTMLNFCQMGSGCTRENLESIITDFLNHLGIDF
KIVGDSCMVFGDTLDVMHGDLELSSAVVGPIPLDREWGIDKPWIGAGFGLERLLKVKHDFKNIKRAARSESYY
NGISTNL*

AcKRS3
MDKKPLDVLISATGLWMSRTGTLHKIKHHEVSRSKIYIEMACGDHLVVNNSRSCRTARAFRHHKYRKTCKRC
RVSDEDINNFLTRSTESKNSVKVRVVSAPKVKKAMPKSVSRAPKPLENSVSAKASTNTSRSVPSPAKSTPNSSV
PASAPAPSLTRSQLDRVEALLSPEDKISLNMAKPFRELEPELVTRRKNDFQRLYTNDREDYLGKLERDITKFFV
DRGFLEIKSPILIPAEYVERMGINNDTELSKQIFRVDKNLCLRPMMAPTIFNYARKLDRILPGPIKIFEVGPCYRK
ESDGKEHLEEFTMVNFFQMGSGCTRENLEALIKEFLDYLEIDFEIVGDSCMVYGDTLDIMHGDLELSSAVVGP
VSLDREWGIDKPWIGAGFGLERLLKVMHGFKNIKRASRSESYYNGISTNL*

### 2 Chemical Synthesis

#### 2.1 Synthesis of pNZ-MeOK 1

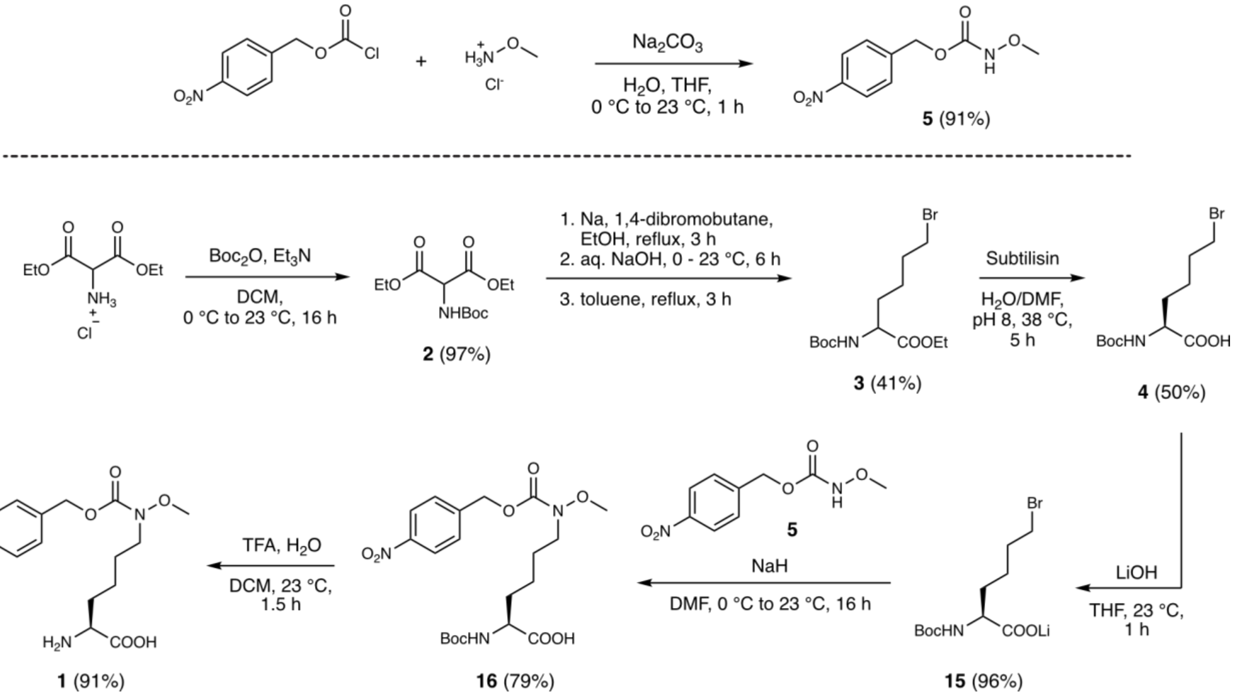

**Scheme S1:** Synthesis of pNZ-MeOK **1**

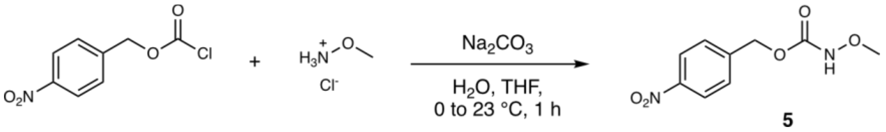

**4-Nitrobenzyl methoxycarbamate (5).** *O*-Methylhydroxylammonium chloride (812 mg, 9.8 mmol, 1.05 eq.) was dissolved in water (46.5 mL) and cooled in an ice bath. $Na_2CO_3$ (2.0 g, 18.6 mmol, 2.0 eq.) was added, the cooling bath removed and THF (46.5 mL) added. 4-Nitrobenzyl chloroformate (2.0 g, 9.3 mmol, 1.0 eq.) was added and the reaction stirred for 1 hour at room temperature. The reaction was then diluted with EtOAc (150 mL), the layers separated and the organic layer washed with water (50 mL) and brine (50 mL), then dried over anhydrous sodium sulfate and concentrated under reduced pressure. The crude solid was purified by recrystallization from a mixture of cyclohexane and EtOAc. 4-Nitrobenzyl methoxycarbamate (**5**, 1.92 g, 8.5 mmol, 91%) was obtained as white needles.

$^{1}$H NMR (500 MHz, $CDCl_3$) δ 8.24 – 8.19 (m, 2H), 7.59 (br s, 1H), 7.54 – 7.49 (m, 2H), 5.28 (s, 2H), 3.76 (s, 3H).

$^{13}$C NMR (125 MHz, $CDCl_3$) δ 157.0, 147.9, 143.0, 128.5, 124.0, 66.0, 65.0.

HRMS $[M+Na]^+$ calculated for $C_9H_{10}N_2NaO_5$ = 249.0482, found 249.0483.

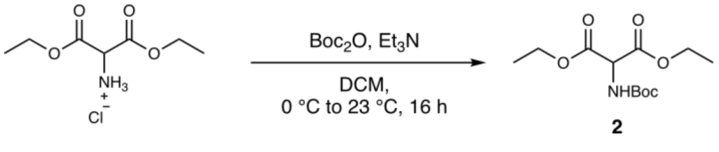

**Diethyl 2-((*tert*-butoxycarbonyl)amino)malonate (2).** Diethyl aminomalonate hydrochloride (29.1 g, 137.5 mmol, 1.2 eq.) was suspended in DCM (400 mL) and cooled in an ice bath. Triethylamine (24.0 mL, 171.8 mmol, 1.5 eq.) was added, followed by di-*tert*-butyl dicarbonate (25.0 g, 114.6 mmol, 1.0 eq.) as a solution in DCM (200 mL). The reaction was stirred for 5 minutes with cooling. The ice bath was removed and stirring continued overnight. Aqueous hydrogen chloride solution (1 M, 200 mL) was added. The layers were separated and the organic layer washed with aqueous hydrogen chloride solution (1 M, 200 mL), water (200 mL) and brine (200 mL). The organic layer was dried over anhydrous sodium sulfate and concentrated under reduced pressure to obtain diethyl 2-((*tert*-butoxycarbonyl)amino)malonate (**2**, 30.6 g, 111.2 mmol, 97%) as a yellow oil.

$^1$H NMR (500 MHz, $CDCl_3$) δ 5.61 – 5.49 (m, 1H), 5.01 – 4.90 (m, 1H), 4.36 – 4.18 (m, 4H), 1.45 (s, 9H), 1.29 (t, $J$ = 7.2 Hz, 6H).

$^{13}$C NMR (125 MHz, $CDCl_3$) δ 166.8, 80.8, 67.3, 62.6, 57.7, 28.4, 14.2.

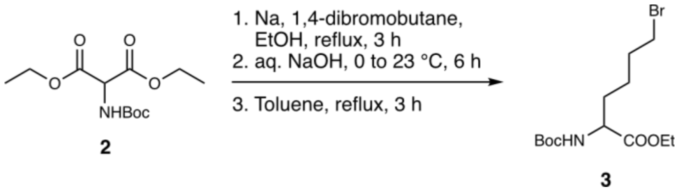

**Ethyl 6-bromo-2-((*tert*-butoxycarbonyl)amino)hexanoate (3).** Absolute ethanol (100 mL) was transferred to a 500 mL round-bottom flask under argon atmosphere. Metallic sodium (2.18 g, 95.0 mmol, 0.95 eq.) was added in smaller pieces and a reflux condenser attached. The flask was heated to 60 °C in an oil bath until the sodium was completely dissolved. The solution was cooled to room temperature and diethyl 2-((*tert*-butoxycarbonyl)amino)malonate (**2**, 27.5 g, 100.0 mmol, 1.0 eq.) added. The reaction was heated to reflux for 45 minutes, then cooled to room temperature. 1,4-Dibromobutane (59.7 mL, 500.0 mmol, 5.0 eq.) was added. The reaction was heated to reflux for 3 hours. Then cooled to 0 °C with an ice bath. Aqueous sodium hydroxide solution (2 M, 70 mL) was added in portions of 10 mL. The ice bath was removed and the reaction stirred at room temperature for 6 hours. Ethanol was removed under reduced pressure. Hexane (200 mL) and saturated aqueous $NaHCO_3$ (400 mL) added. Layers were separated and the aqueous layer washed with hexane (200 mL). The aqueous layer was acidified with solid citric acid to pH 3 – 4, then extracted with EtOAc (3 x 150 mL). The combined organic layers were washed with brine, dried over anhydrous sodium sulfate and concentrated under reduced pressure to obtain the monoester as a yellowish oil. The residue was dissolved in toluene (250 mL) and heated to reflux for 3 hours. The solution was allowed to cool to room temperature and concentrated under reduced pressure. The residue was purified by flash column chromatography on silica gel (gradient 0 – 30% EtOAc in hexane) to obtain ethyl 6-bromo-2-((*tert*-butoxycarbonyl)amino)hexanoate (**3**, 13.7 g, 40.5 mmol, 41%) as a colourless oil.

$^1$H NMR (500 MHz, $CDCl_3$) δ 5.13 – 4.94 (m, 1H), 4.34 – 4.25 (m, 1H), 4.25 – 4.15 (m, 2H), 3.40 (t, $J$ = 6.7 Hz, 2H), 1.95 – 1.77 (m, 3H), 1.71 – 1.60 (m, 1H), 1.44 (s, 9H), 1.28 (t, $J$ = 7.1 Hz, 3H).

$^{13}$C NMR (125 MHz, $CDCl_3$) δ 172.8, 155.5, 80.0, 61.5, 53.3, 33.4, 32.3, 32.1, 28.5, 24.0, 14.4.

HRMS $[M+Na]^+$ calculated for $C_{13}H_{24}BrNNaO_4$ = 360.0781, found 360.0775.

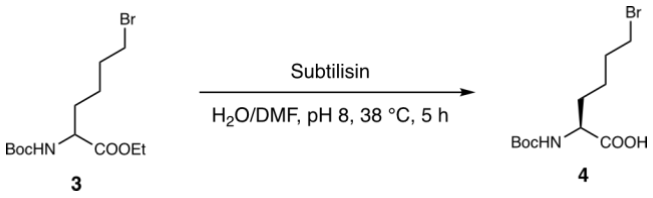

**(*S*)-6-Bromo-2-((*tert*-butoxycarbonyl)amino)hexanoic acid (4).** Ethyl 6-bromo-2-((*tert*-butoxycarbonyl)amino)hexanoate (**3**, 28.0 g, 82.8 mmol, 1.0 eq.) was dissolved in water (213 mL) and DMF (71 mL) and the pH of the solution adjusted to approximately 8 using aqueous ammonia (1 M). The solution was warmed to 38 °C and subtilisin (41.4 mg, 0.5 mg/mmol) added. The reaction was stirred at 38 °C and the pH regularly checked to maintain at or slightly above pH 8 by addition of aqueous ammonia (1 M). After 7 hours, the reaction was concentrated under reduced pressure. The residue was dissolved in hexane (300 mL) and aqueous half-saturated $NaHCO_3$ (500 mL), the layers separated and the aqueous layer washed with hexane (200 mL). The aqueous layer was acidified with solid citric acid to a pH of 3 – 4, then extracted with EtOAc (3 x 200 mL). The combined EtOAc layers were washed with brine (200 mL), dried over anhydrous sodium sulfate and concentrated under reduced pressure to obtain (*S*)-6-bromo-2-((*tert*-butoxycarbonyl)amino)hexanoic acid (**4**, 12.7 g, 41.0 mmol, 50%) as a colorless residue.

$^{1}H$ NMR (500 MHz, $CDCl_3$) δ 5.10 – 4.97 (m, 1H), 4.39 – 4.25 (m, 1H), 3.41 (t, *J* = 6.7 Hz, 2H), 2.00 – 1.81 (m, 3H, 1.81 – 1.64 (m, 1H), 1.63 – 1.49 (m, 2H), 1.45 (s, 9H).

$^{13}C$ NMR (125 MHz, $CDCl_3$) δ 177.1, 155.8, 80.5, 53.2, 33.3, 32.2, 31.7, 28.5, 24.1.

HRMS $[M+Na]^+$ calculated for $C_{11}H_{20}BrNNaO_4$ = 332.0468, found 332.0461.

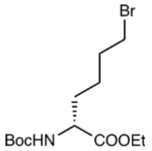

**14**

To obtain the *R*-enantiomer **14**, the hexane layers were washed with brine (150 mL), dried over anhydrous sodium sulfate and concentrated under reduced pressure. Ethyl (*R*)-6-bromo-2-((*tert*-butoxycarbonyl)amino)hexanoate (**14**, 14.0 g, 41.4 mmol, 50%) was obtained as a colourless oil.

$^{1}H$ NMR (500 MHz, $CDCl_3$) δ 5.11 – 4.97 (m, 1H), 4.35 – 4.24 (m, 1H), 4.24 – 4.14 (m, 2H), 3.39 (t, *J* = 6.8 Hz, 2H), 1.97 – 1.77 (m, 3H), 1.70 – 1.58 (m, 1H), 1.57 – 1.46 (m, 2H), 1.44 (s, 9H), 1.28 (t, *J* = 7.1 Hz, 3H).

$^{13}C$ NMR (125 MHz, $CDCl_3$) δ 172.8, 155.5, 80.0, 61.5, 53.3, 33.4, 32.2, 32.1, 28.5, 24.0, 14.3.

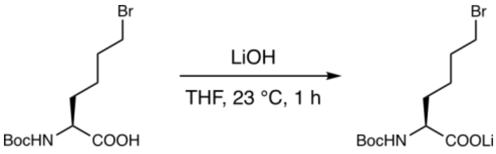

**4** **15**

**(*S*)-6-Bromo-2-((*tert*-butoxycarbonyl)amino)hexanoate lithium salt (15).** (*S*)-6-Bromo-2-((*tert*-butoxycarbonyl)amino)hexanoic acid (**4**, 12.7 g, 41.0 mmol, 1.0 eq.) was dissolved in THF (240 mL) and a solution of LiOH (933 mg, 39.0 mmol, 0.95 eq.) in water (15 mL) added. The reaction was stirred for 1 hour, then concentrated under reduced pressure and coevaporated with acetonitrile. The resulting white solid was freeze dried to obtain (*S*)-6-bromo-2-((*tert*-butoxycarbonyl)amino)hexanoate lithium salt (**15**, 12.5 g, 39.6 mmol, 96%) as a white powder.

$^{1}H$ NMR (500 MHz, MeOD) δ 4.03 – 3.95 (m, 1H), 3.43 (t, *J* = 6.7 Hz, 2H), 1.92 – 1.76 (m, 3H), 1.69 – 1.58 (m, 1H), 1.56 – 1.46 (m, 2H), 1.44 (s, 9H).

$^{13}C$ NMR (125 MHz, MeOD) δ 179.1, 157.7, 80.0, 56.7, 34.2, 33.8, 33.4, 28.8, 25.3.

HRMS $[M+H]^+$ calculated for $C_{11}H_{20}BrLiNO_4$ = 316.073, found 316.073.

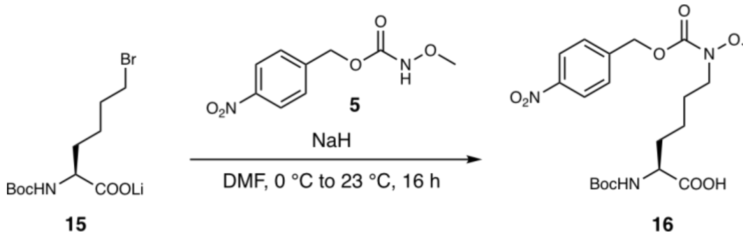

**$N^2$-(*tert*-butoxycarbonyl)-$N^6$-methoxy-$N^6$-(((4-nitrobenzyl)oxy)carbonyl)-*L*-lysine (16).** In a flask under inert atmosphere, 4-nitrobenzyl methoxycarbamate (**5**, 5.24 g, 23.2 mmol, 1.25 eq.) was dissolved in anhydrous DMF (28.5 mL) and cooled in an ice bath. To the solution was added NaH (60% dispersion in mineral oil, 890 mg, 22.2 mmol, 1.2 eq.). Cooling was removed and the reaction stirred for 30 minutes at room temperature, then again cooled in an ice bath**.** (*S*)-6-bromo-2-((*tert*-butoxycarbonyl)amino)hexanoate lithium salt (**15**, 5.9 g, 18.5 mmol, 1.0 eq.) was added to the reaction as a solution in anhydrous DMF (28.5 mL). Stirring was continued for 30 minutes at 0 °C, then overnight at room temperature. The reaction was again cooled in an ice bath and quenched by addition of 10% aq. $KH_2PO_4$ (pH 3.5-4, 150 mL). If needed, the pH of the aqueous solution was adjusted to 3-4 by addition of solid citric acid. The aqueous solution was extracted with EtOAc (1 x 300 mL). The organic layer was washed with 10% aq. $KH_2PO_4$ (pH 3.5-4, 2 x 100 mL) and brine (100 mL), then dried over anhydrous sodium sulfate and concentrated under reduced pressure. The resulting residue was purified by flash column chromatography (gradient from 0 to 4% MeOH in DCM with 0.5% acetic acid) to obtain $N^2$-(*tert*-butoxycarbonyl)-$N^6$-methoxy-$N^6$-(((4-nitrobenzyl)oxy)carbonyl)-*L*-lysine (**16**, 6.7 g, 14.6 mmol, 79%) as a viscous yellow oil.

$^1$H NMR (500 MHz, $CDCl_3$) δ 8.26 – 8.21 (m, 2H), 7.56 – 7.52 (m, 2H), 5.28 (s, 2H), 5.10 – 5.01 (m, 1H), 4.33 – 4.24 (m, 1H), 3.71 (s, 3H), 3.58 – 3.48 (m, 2H), 1.96 – 1.81 (m, 1H), 1.76 – 1.61 (m, 3H), 1.49 – 1.36 (m, 11H).

$^{13}$C NMR (125 MHz, $CDCl_3$) δ 176.4, 156.5, 155.9, 147.9, 143.5, 128.4, 124.0, 80.5, 66.3, 62.6, 53.3, 48.4, 31.9, 28.4, 26.7, 22.5.

HRMS $[M+Na]^+$ calculated for $C_{20}H_{29}N_3NaO_9$ = 478.1796, found 478.1788.

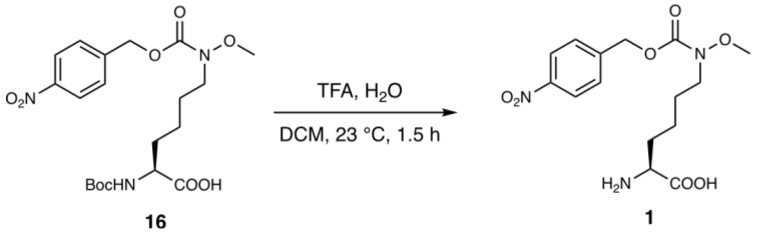

**$N^6$-Methoxy-$N^6$-(((4-nitrobenzyl)oxy)carbonyl)-*L*-lysine (pNZ-MeOK, 1).** $N^2$-(*tert*-Butoxycarbonyl)-$N^6$-methoxy-$N^6$-(((4-nitrobenzyl)oxy)carbonyl)-*L*-lysine (**16**, 6.7 g, 14.6 mmol, 1.0 eq.) was dissolved in DCM (33.6 mL) and water (1.1 mL, 58.5 mmol, 4.0 eq.) added. Then, trifluoroacetic acid (33.6 mL, 439 mmol, 30.0 eq.) was added and the reaction stirred for 1.5 hours. The reaction was concentrated under reduced pressure. The residue was purified by reversed phase HPLC and lyophilized to obtain $N^6$-methoxy-$N^6$-(((4-nitrobenzyl)oxy)carbonyl)-*L*-lysine (**1**, 4.8 g, 14.6 mmol, 91%) as a white lyophilized solid.

$^1$H NMR (500 MHz, DMSO-$d_6$) δ 8.28 – 8.23 (m, 2H), 7.67 – 7.62 (m, 2H), 5.28 (s, 2H), 3.64 (s, 3H), 3.46 (t, $J$ = 7.2 Hz, 2H), 3.16 – 3.10 (m, 1H), 1.78 – 1.67 (m, 1H), 1.64 – 1.49 (m, 3H), 1.43 – 1.28 (m, 2H).

$^{13}$C NMR (125 MHz, DMSO-$d_6$) δ 169.8, 155.6, 147.1, 144.2, 128.3, 132.7, 65.5, 61.9, 54.0, 48.0, 30.8, 26.5, 22.3.

HRMS $[M+H]^+$ calculated for $C_{15}H_{22}N_3O_7$ = 356.1452, found 356.1449.

#### 2.2 Synthesis of oAZ-MeOK 13

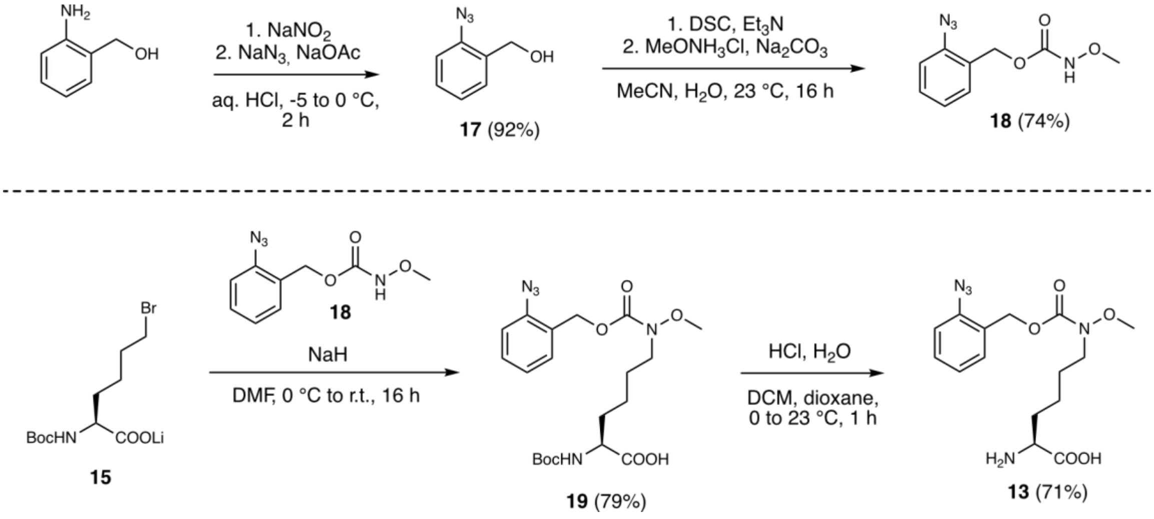

**Scheme S2:** Synthesis of oAZ-MeOK **13**

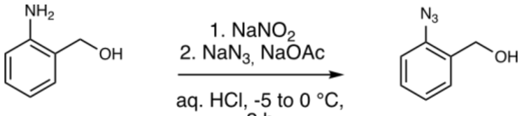

**17**

**(2-Azidophenyl)methanol (17).** In a 100 mL flask, (2-aminophenyl)methanol (2.0 g, 16.2 mmol, 1.0 eq.) was suspended in aq. HCl (2 M, 25 mL) and cooled to -5 °C in a salt/ice bath. Sodium nitrite (1.3 g, 19.5 mmol, 1.2 eq.) was dissolved in water (5 mL), cooled to 0 °C and added dropwise. The reaction was stirred for 15 minutes at -5 °C, then urea (195 mg, 3.25 mmol, 0.2 eq.) added. In a separate 250 mL flask, sodium acetate (4.0 g, 48.7 mmol, 3 eq.) was dissolved in water (25 mL) and sodium azide (2.1 g, 32.5 mmol, 2.0 eq.) added. The solution was cooled in an ice bath. Then, the solution containing the diazonium salt was added dropwise while the reaction was vigorously stirred. After complete addition, the reaction was stirred for 2 hours at 0 °C. The reaction was extracted with diethyl ether (150 mL). The organic layer was washed twice with aq. NaOH (1 M, 50 mL), water (50 mL) and brine (50 mL), then dried over anhydrous sodium sulfate, filtered and concentrated under reduced pressure to obtain (2-azidophenyl)methanol (**17**, 2.2 g, 14.9 mmol, 92%) as a dark orange liquid.

$^{1}$H NMR (400 MHz, $CDCl_3$) δ 7.40 – 7.32 (m, 2H), 7.20 – 7.11 (m, 2H), 4.64 (s, 2H).

$^{13}$C NMR (101 MHz, $CDCl_3$) δ 138.0, 132.0, 129.4, 129.3, 125.1, 118.2, 61.8.

HRMS $[M]^+$ calculated for $C_7H_7N_3O$ = 149.0584, found 149.0583.

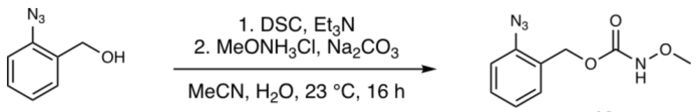

**18**

**2-Azidobenzyl methoxycarbamate (18).** In a 250 mL flask, (2-Azidophenyl)methanol (2.2 g, 14.9 mmol, 1.0 eq.) was dissolved in acetonitrile (80 mL) and triethylamine (4.2 mL, 29.8 mmol, 2.0 eq.) added, followed by *N,N*-disuccinimidyl carbonate (5.7 g, 22.3 mmol, 1.5 eq.). The reaction was stirred at room temperature for one hour. In a separate flask, *O*-methylhydroxylamine hydrochloride (1.9 g, 22.3 mmol, 1.5 eq.) was combined with acetonitrile (40 mL) and sodium carbonate (3.2 g, 29.8 mmol, 2.0 eq.). Water was added to the suspension until all solids completely dissolved. Then, the solution was added to the activated carbonate and stirred for 16 hours. The reaction was concentrated under reduced pressure and the residue dissolved in EtOAc (100 mL) and water (50 mL). The layers were separated and the organic layer washed with aq. HCl (1 M, 50 mL), water (50 mL) and brine (50 mL). The organic layer was dried over anhydrous sodium sulfate, filtered and concentrated under reduced pressure. The crude product was purified by flash column chromatography (gradient from 5 to 35% EtOAc in hexane) to obtain 2-azidobenzyl methoxycarbamate (**18**, 2.5 g, 11.0 mmol, 74%) as a yellow oil.

$^{1}$H NMR (400 MHz, $CDCl_3$) δ 7.48 – 7.35 (m, 3H), 7.22 – 7.11 (m, 2H), 5.17 (s, 2H), 3.75 (s, 3H).

$^{13}$C NMR (101 MHz, $CDCl_3$) δ 157.4, 138.8, 130.6, 130.1, 126.9, 125.0, 118.4, 64.9, 63.2.

HRMS $[M]^+$ calculated for $C_9H_{10}N_4NaO_3$ = 245.0645, found 245.0644.

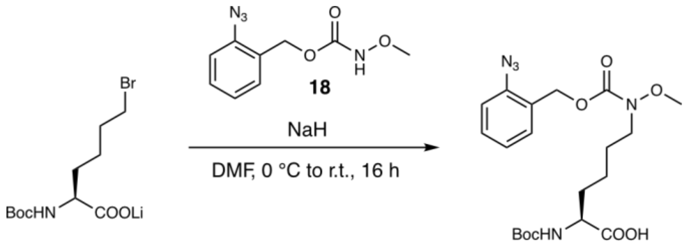

15 19

***N*⁶-(((2-azidobenzyl)oxy)carbonyl)-*N*²-(*tert*-butoxycarbonyl)-*N*⁶-methoxy-*L*-lysine (19).** In a flask under inert atmosphere, 2-azidobenzyl methoxycarbamate (**18**, 4.8 g, 21.8 mmol, 1.25 eq.) was dissolved in anhydrous DMF (22 mL). To the solution was added NaH (60% dispersion in mineral oil, 835 mg, 20.9 mmol, 1.2 eq.). The reaction was stirred for 30 minutes at room temperature, then cooled in an ice bath**.** (*S*)-6-bromo-2-((*tert*-butoxycarbonyl)amino)hexanoate lithium salt (**15**, 5.5 g, 17.4 mmol, 1.0 eq.) was added to the reaction as a solution in anhydrous DMF (22 mL). Stirring was continued for 30 minutes at 0 °C, then overnight at room temperature. The reaction was again cooled in an ice bath and quenched by addition of 10% aq. $KH_2PO_4$ (pH 3.5-4, 150 mL). If needed, the pH of the aqueous solution was adjusted to 3-4 by addition of solid citric acid. The aqueous solution was extracted with EtOAc (1 x 300 mL). The organic layer was washed with 10% aq. $KH_2PO_4$ (pH 3.5-4, 2 x 100 mL) and brine (100 mL), then dried over anhydrous sodium sulfate and concentrated under reduced pressure. The resulting residue was purified by flash column chromatography (gradient from 0 to 3% MeOH in DCM with 0.5% acetic acid) to obtain *N*⁶-(((2-azidobenzyl)oxy)carbonyl)-*N*²-(*tert*-butoxycarbonyl)-*N*⁶-methoxy-*L*-lysine (**19**, 6.22 g, 13.8 mmol, 79%) as a pale orange residue.

$^{1}$H NMR (500 MHz, $CDCl_3$) δ 7.42 – 7.34 (m, 2H), 7.21 – 7.11 (m, 2H), 5.17 (s, 2H), 5.13 – 5.01 (m, 1H), 4.35 – 4.21 (m, 1H), 3.69 (s, 3H), 3.61 – 3.43 (m, 2H), 1.96 – 1.78 (m, 1H), 1.77 – 1.58 (m, 3H), 1.52 – 1.34 (m, 2H), 1.44 (s, 9H)

$^{13}$C NMR (125 MHz, $CDCl_3$) δ 176.4, 157.0, 156.0, 138.6, 130.3, 129.8, 127.4, 125.0, 118.4, 80.5, 63.4, 62.5, 53.4, 48.4, 31.8, 28.4, 26.8, 22.5

HRMS $[M+Na]^+$ calculated for $C_{20}H_{29}N_5NaO_7$ = 474.1959, found 474.1956.

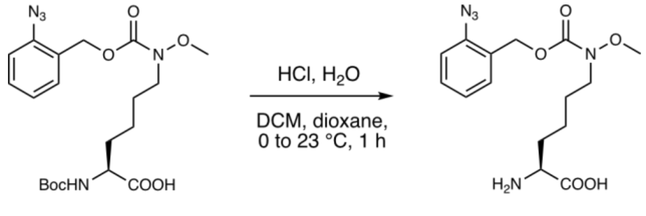

19 13

***N*⁶-(((2-azidobenzyl)oxy)carbonyl)-*N*⁶-methoxy-*L*-lysine (oAZ-MeOK, 13)**. *N*⁶-(((2-Azidobenzyl)oxy)carbonyl)-*N*²-(*tert*-butoxycarbonyl)-*N*⁶-methoxy-*L*-lysine (**19**, 6.2 g, 13.8 mmol, 1.0 eq.) was dissolved in DCM (100 mL) and water (496 µL, 27.6 mmol, 2.0 eq.) added. The solution was cooled in an ice bath and 4 M HCl in dioxane (41.3 mL, 165.3 mmol, 12 eq.) added. After 5 minutes, the ice bath was removed and the reaction stirred at room temperature for 1 hour. The reaction was concentrated under reduced pressure to remove all volatiles. The resulting crude residue was purified by reversed phase HPLC and lyophilized to obtain *N*⁶-(((2-azidobenzyl)oxy)carbonyl)-*N*⁶-methoxy-*L*-lysine (**13**, 3.4 g, 9.7 mmol, 71%) as a white, lyophilized solid.

$^1H$ NMR (500 MHz, DMSO-$d_6$) δ 7.48 – 7.38 (m, 2H), 7.37 – 7.32 (m, 1H), 7.27 – 7.19 (m, 1H), 5.07 (s, 2H), 3.61 (s, 3H), 3.42 (t, $J$ = 7.2 Hz, 2H), 3.14 – 3.08 (m, 1H), 1.76 – 1.67 (m, 1H), 1.63 – 1.55 (m, 1H), 1.54 – 1.46 (m, 2H), 1.40 – 1.25 (m, 2H)

$^{13}C$ NMR (125 MHz, DMSO-$d_6$) δ 169.8, 155.8, 137.7, 129.9, 129.7, 127.0, 125.1, 118.8, 62.5, 61.8, 54.0, 48.1, 30.8, 26.5, 22.4

HRMS $[M+H]^+$ calculated for $C_{15}H_{22}NNaO_4$ = 360.0781, found 360.0775.

#### 2.3 Synthesis of acylboronates

##### 2.3.1 General remarks and procedures

**General remarks to synthesis**

When purifying MIDA acylboronates by flash column chromatography, it is recommended to concentrate the crude product on celite (not silica) for dry loading or direct application as a solution. MIDA acylboronates were generally purified fast over short/small columns to limit the time the compound is in contact with silica.

**General procedure A:**

General procedure A was adapted from a literature procedure.[10] A heat-dried flask under inert atmosphere was charged with 2,2,6,6-tetramethylpiperidine (1.1 eq.) and anhydrous THF (0.67 mL/mmol) added. The solution was cooled in an ice bath. *n*-BuLi (1.6 M solution in hexane, 1.1 eq.) was added dropwise. The mixture was stirred at 0 °C for 10 minutes. In a separate heat-dried flask under inert atmosphere, bis(4,4,5,5-tetramethyl-1,3,2-dioxaborolan-2-yl)methane (1.0 eq.) was dissolved in anhydrous THF (2 mL/mmol) and cooled in an ice bath. The LiTMP solution was added and the reaction stirred for 5 minutes. Then, the alkyl halide or other compatible electrophile (1.1 eq.) was added. After 5 minutes, the cooling was removed and the reaction allowed to warm to room temperature. After 1 – 16 hours, the reaction was quenched with saturated aqueous $NH_4Cl$ (approx. 5 mL/mmol). $Et_2O$ and little water (to dissolve remaining solid salts) were added, layers separated and the aqueous layer extracted with $Et_2O$. Combined organic layers were dried over anhydrous sodium sulfate, filtered and concentrated under reduced pressure. The crude product was purified by flash column chromatography on silica gel to obtain the pure product.

**General procedure B:**

General procedure B was performed as described in the literature.[11] Methyliminodiacetic acid (6.0 eq.) was added to a heat-dried high-pressure reaction tube under a counterflow of argon. The starting material (1.0 eq.) was either added as a solid or as a solution together with the solvent. Anhydrous DMSO (1.5 mL/mmol) was added, followed by triethyl orthoformate (4.0 eq.). The tube was capped and placed in a preheated heat block at 130 °C. The reaction was stirred vigorously for 16 – 24 hours. Then, the reaction was allowed to cool down to room temperature and water (7.5 mL/mmol) added. EtOAc was added, layers separated and the aqueous layer further extracted twice with EtOAc. The combined organic layers were washed with brine, dried over anhydrous sodium sulfate, filtered and concentrated under reduced pressure. The crude product was purified by flash column chromatography on silica gel to obtain the pure product. The unreacted starting material can be recovered.

**General procedure C:**

General procedure C was performed as described in the literature.[11] In a round-bottom flask, the unsymmetrical geminal diboryl substrate (1.0 eq.) was dissolved or suspended in a mixture of THF (1.88 mL/mmol) and buffer ($KH_2PO_4$/NaOH, pH = 7, 1M, 1.88 mL/mmol). The flask was cooled in an ice bath and $NaBO_3 \cdot H_2O$ (1.3 eq.) added. The reaction was allowed to warm to room temperature over 3 hours. Then, the reaction was diluted with EtOAc and anhydrous sodium sulfate was added. The solution was filtered and the remaining sodium sulfate washed at least five times with EtOAc. The organic layer was concentrated under reduced pressure. The crude product was purified by flash column chromatography on silica gel to obtain the pure product.

**General procedure D:**

General procedure D was performed as described in the literature[12] with minor modifications (use of PBS instead of phosphate buffer). In a round-bottom flask, the vinyl boronate (1.0 eq.) was dissolved in acetone (2.66 mL/mmol) and PBS pH 7 (667 µL/mmol) added, followed by NMO (3.0 eq.) and $OsO_4$ (4% in $H_2O$, 0.05 eq.). After complete consumption of the starting material was observed by TLC (approx. 1 hour), sodium sulfite (3.0 eq.) was added to quench the reaction and further stirred for 30 minutes. The reaction was then transferred to a separatory funnel with EtOAc and brine (approx. 25 mL/mmol) added. Layers were separated and the aqueous layer extracted with EtOAc. The combined organic layers were dried over anhydrous sodium sulfate, filtered and concentrated under reduced pressure. The crude product was purified by flash column chromatography on silica gel to obtain the pure product.

**General procedure E:**
General procedure E was performed as described in the literature.[12] The vicinal diol substrate (1.0 eq.) was dissolved or suspended in DCM (20 mL/mmol) and finely powdered sodium periodate (1.05 eq.) added. The reaction was stirred vigorously and water (1 mL/mmol) added. The progress of the reaction was monitored by TLC. After completion, the reaction was transferred to a separatory funnel with DCM and water added. The layers were separated and the aq. Layer extracted with DCM. The combined organic layers were dried over anhydrous sodium sulfate, filtered and concentrated under reduced pressure. Usually, the obtained product did not need any further purification.

##### 2.3.2 Synthesis of MIDA acetylboronate 9a

**2,2'-(Ethane-1,1-diyl)bis(4,4,5,5-tetramethyl-1,3,2-dioxaborolane) (20)**

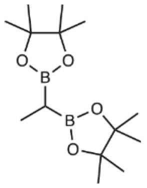

Compound **20** was prepared according to general procedure A, starting from bis[(pinacolato)boryl]methane (400 mg, 1.49 mmol, 1.0 eq.) and methyl iodide (102 µL, 1.64 mmol, 1.1 eq.). The crude product was purified by flash column chromatography (gradient of 0 to 40% EtOAc in hexane) to obtain the desired product (**20**, 344 mg, 1.22 mmol, 82%) as a colourless oil.

The recorded NMR data is in accordance with the literature.[13]

$^{1}$H NMR (500 MHz, $CDCl_3$) δ 1.23 (s, 12H), 1.22 (s, 12H), 1.04 (d, *J* = 7.3 Hz, 3H), 0.72 (q, *J* = 7.3 Hz, 1H).

$^{13}$C NMR (125 MHz, $CDCl_3$) δ 83.1, 25.0, 24.7, 9.2.

$^{11}$B NMR (128 MHz, $CDCl_3$) δ 34.2.

HRMS $[M+Na]^+$ calculated for $C_{14}H_{28}B_2NaO_4$ = 305.2066, found 305.2062.

**1-(4,4,5,5-Tetramethyl-1,3,2-dioxaborolan-2-yl)ethylboronic acid MIDA ester (21)**

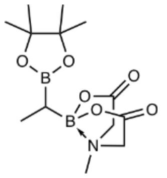

Compound **21** was prepared according to general procedure B, starting from **20** (325 mg, 1.15 mmol, 1.0 eq.). The crude product was purified by flash column chromatography (gradient of 80 – 100% EtOAc in cyclohexane) to obtain **21** (216 mg, 0.7 mmol, 60%) as a white solid.

$^{1}$H NMR (400 MHz, $CDCl_3$) δ 4.09 (d, *J* = 15.9 Hz, 1H), 3.81 – 3.77 (m, 2H), 3.74 (d, *J* = 15.9 Hz, 1H), 3.02 (s, 3H), 1.22 (s, 6H), 1.22 (s, 6, 1.12 (d, *J* = 7.6 Hz, 3H), 0.43 (q, *J* = 7.6 Hz, 1H).

$^{13}$C NMR (101 MHz, $CDCl_3$) δ 167.4, 167.2, 83.4, 63.6, 63.0, 46.2, 25.2, 24.8, 10.3.
$^{11}$B NMR (128 MHz, $CD_3CN$) δ 34.3, 14.0.

HRMS $[M+Na]^+$ calculated for $C_{13}H_{23}B_2NNaO_6$ = 334.1604, found 334.1605.

**(1-Hydroxyethyl)boronic acid MIDA ester (22)**

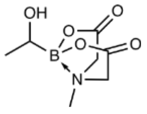

Compound **22** was prepared according to general procedure C from **21** (190 mg, 611 µmol, 1.0 eq.). The crude product was purified by flash column chromatography (gradient of 80 to 100% EtOAc in cyclohexane) to obtain **22** (120 mg, 597 µmol, 98%) as a white solid.

The recorded NMR data is in accordance with the literature.[14]

$^{1}$H NMR (500 MHz, $CD_3CN$) δ 3.93 (d, *J* = 17.1 Hz, 1H), 3.90 (d, *J* = 16.2 Hz, 1H), 3.80 (d, *J* = 16.2 Hz, 1H), 3.79 (d, *J* = 17.1 Hz, 1H), 3.43 (dq, *J* = 7.2, 4.1 Hz, 1H), 3.02 (s, 3H), 2.33 (d, *J* = 4.1 Hz, 1H), 1.18 (d, *J* = 7.2 Hz, 3H).

$^{13}$C NMR (125 MHz, $CD_3CN$) δ 169.9, 169.0, 63.3, 63.1, 46.2, 19.7.

$^{11}$B NMR (128 MHz, $CD_3CN$) δ 11.18.

HRMS $[M+Na]^+$ calculated for $C_7H_{12}BNNaO_5$ = 224.0701, found 224.0699.

**MIDA acetylboronate (9a)**

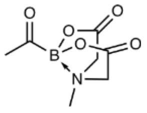

Substrate **22** (42.3 mg, 211 µmol, 1.0 eq.) was dissolved in acetonitrile (12.4 mL) and DMP (134 mg, 316 µmol, 1.5 eq.) added. The reaction was stirred open to atmosphere for 30 minutes, then filtered over a pad of celite and the filtrate concentrated. The crude product was purified by flash column chromatography (gradient of 80 to 100% EtOAc in hexane, dry load on celite) to obtain the desired product **9a** (38.4 mg, 193 µmol, 92%) as a white solid.

The recorded NMR data is in accordance with the literature.[14]

$^{1}$H NMR (500 MHz, $CD_3CN$) δ 4.03 (d, *J* = 17.0 Hz, 2H), 3.88 (d, *J* = 17.0 Hz, 2H), 2.83 (s, 3H), 2.23 (s, 3H).

$^{13}$C NMR (125 MHz, $CD_3CN$) δ 169.0, 63.0, 47.4.

$^{11}$B NMR (161 MHz, $CD_3CN$) δ 4.2.

HRMS $[M+Na]^+$ calculated for $C_7H_{10}BNNaO_5$ = 222.0544, found 222.0544.

**2.3.3 Synthesis of MIDA propionylboronate 9b**

**2,2'-(propane-1,1-diyl)bis(4,4,5,5-tetramethyl-1,3,2-dioxaborolane) (23)**

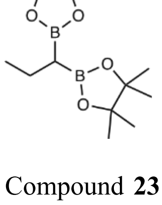

Compound **23** was prepared according to general procedure A, starting from bis[(pinacolato)boryl]methane (800 mg, 2.99 mmol, 1.0 eq.) and ethyl bromide (243 µL, 3.28 mmol, 1.1 eq.). The reaction was stirred for 2 hours after the addition of ethyl bromide. The crude product was purified by flash column chromatography (gradient from 0 to 15% EtOAc in hexane) to obtain the desired product **23** (680 mg, 2.3 mmol, 77%) as a colourless oil.

$^{1}$H NMR (500 MHz, $CDCl_3$) δ 1.62 – 1.53 (m, 2H), 1.23 (s, 12H), 1.22 (s, 12H), 0.92 (t, *J* = 7.4 Hz, 3H), 0.66 (t, *J* = 7.8 Hz, 1H).

$^{13}$C NMR (125 MHz, $CDCl_3$) δ 83.0, 25.0, 24.7, 19.2, 17.2.

$^{11}$B NMR (161 MHz, $CDCl_3$) δ 34.3.

HRMS $[M+Na]^+$ calculated for $C_{15}H_{30}B_2NaO_4$ = 319.2222, found 319.2227.

**1-(4,4,5,5-Tetramethyl-1,3,2-dioxaborolan-2-yl)propylboronic acid MIDA ester (24)**

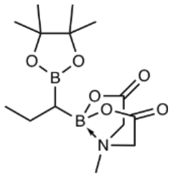

Compound **24** was prepared according to general procedure B from **23** (661 mg, 2.23 mmol, 1.0 eq.). The crude product was purified by flash column chromatography (gradient from 70 to 100% EtOAc in hexane) to obtain the desired product **24** (360 mg, 1.11 mmol, 50%) as a white solid.

$^{1}$H NMR (500 MHz, $CD_3CN$) δ 3.90 (dd, *J* = 17.1, 15.2 Hz, 2H), 3.81 (dd, *J* = 17.8, 17.0 Hz, 2H), 2.93 (s, 3H), 1.53 – 1.42 (m, 2H), 1.20 (s, 12H), 0.95 (t, *J* = 7.4 Hz, 3H), 0.40 – 0.30 (m, 1H).

$^{13}$C NMR (125 MHz, $CD_3CN$) δ 169.2, 169.0, 83.7, 63.5, 63.4, 46.9, 25.1, 20.5, 17.0.

$^{11}$B NMR (161 MHz, $CD_3CN$) δ 34.7, 13.4.

HRMS $[M+NH_4]^+$ calculated for $C_{14}H_{29}B_2N_2O_6$ = 343.2206, found 343.2206.

**(1-Hydroxypropyl)boronic acid MIDA ester (25)**

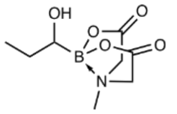

Compound **25** was prepared according to general procedure C from **24** (275 mg, 846 μmol, 1.0 eq.). The crude product was purified by flash column chromatography (gradient from 0 to 70% MeCN in DCM) to obtain **25** (171 mg, 795 μmol, 94%) as a white solid.
$^{1}$H NMR (500 MHz, $CD_3CN$) δ 3.90 (dd, *J* = 17.2, 8.1 Hz, 2H), 3.80 (dd, *J* = 17.2, 15.7 Hz, 2H), 3.19 – 3.12 (m, 1H), 3.02 (s, 3H), 2.35 (d, *J* = 4.9 Hz, 1H), 1.64 – 1.54 (m, 1H), 1.53 – 1.41 (m, 1H), 0.96 (t, *J* = 7.4 Hz, 3H).

$^{13}$C NMR (125 MHz, $CD_3CN$) δ 169.9, 169.0, 63.2, 62.8, 46.1, 27.1, 11.4.

$^{11}$B NMR (161 MHz, $CD_3CN$) δ 11.1.

HRMS $[M+Na]^+$ calculated for $C_8H_{14}BNNaO_5$ = 238.0857, found 238.0858.

**MIDA propionylboronate (9b)**

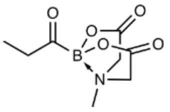

Substrate **25** (118 mg, 549 µmol, 1.0 eq.) was dissolved in acetonitrile (30 mL) and dichloromethane (6 mL). DMP (349 mg, 823 µmol, 1.5 eq.) was added. The reaction was stirred open to atmosphere for 2 hours, then filtered over a pad of celite and the filtrate concentrated. The crude product was purified by flash column chromatography (gradient of 10 to 60% MeCN in DCM, dry load on celite) to obtain the desired product **9b** (79.0 mg, 371 µmol, 68%) as a white solid.

$^{1}$H NMR (500 MHz, $CD_3CN$) δ 4.02 (d, *J* = 16.9 Hz, 2H), 3.89 (d, *J* = 16.9 Hz, 2H), 2.80 (s, 3H), 2.65 (q, *J* = 7.2 Hz, 0.93 (t, *J* = 7.2 Hz, 3H).

$^{13}$C NMR (125 MHz, $CD_3CN$) δ 169.0, 62.9, 47.3, 6.3.

$^{11}$B NMR (161 MHz, $CD_3CN$) δ 4.4.

HRMS $[M+Na]^+$ calculated for $C_8H_{12}BNNaO_5$ = 236.0701, found 236.0703.

##### 2.3.4 Synthesis of MIDA butyrylboronate 9c

**2,2'-(butane-1,1-diyl)bis(4,4,5,5-tetramethyl-1,3,2-dioxaborolane) (26)**

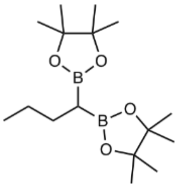

Compound **26** was prepared according to general procedure A from bis[(pinacolato)boryl]methane (400 mg, 1.49 mmol, 1.0 eq.) and propyl bromide (149 µL, 1.64 mmol, 1.1 eq.). The reaction was stirred for 4 hours after addition of the alkyl bromide. The crude product was purified by flash column chromatography (gradient of 0 to 15% EtOAc in cyclohexane) to obtain the desired product **26** (328 mg, 1.06 mmol, 71%) as a colourless oil.
The recorded NMR data is in accordance with the literature.[13]

$^{1}$H NMR (500 MHz, $CDCl_3$) δ 1.56 – 1.49 (m, 2H), 1.36 – 1.25 (m, 2H), 1.23 (s, 12H), 1.22 (s, 12H), 0.87 (t, *J* = 7.2 Hz, 3H), 0.73 (t, *J* = 7.8 Hz, 1H).

$^{13}$C NMR (125 MHz, $CDCl_3$) δ 83.0, 28.1, 25.8, 25.0, 24.7, 14.3.

$^{11}$B NMR (161 MHz, $CDCl_3$) δ 34.0.

HRMS $[M+Na]^+$ calculated for $C_{16}H_{32}B_2NaO_4$ = 333.2379, found 333.238.

**1-(4,4,5,5-tetramethyl-1,3,2-dioxaborolan-2-yl)butylboronic acid MIDA ester (27)**

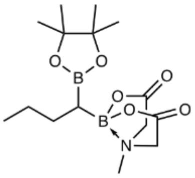

Compound **27** was prepared according to general procedure B from **26** (307 mg, 990 µmol, 1.0 eq.). The crude product was purified by flash column chromatography (gradient of 80 – 100% EtOAc in cyclohexane) to obtain the desired product **27** (159 mg, 469 µmol, 47%) as an off-white solid.

$^{1}H$ NMR (400 MHz, $CDCl_3$) δ 3.94 (d, *J* = 16.0 Hz, 1H), 3.77 (s, 2H), 3.73 (d, *J* = 16.0 Hz, 1H), 3.01 (s, 3H), 1.65 – 1.41 (m, 3H), 1.36 – 1.25 (m, 1H), 1.22 (s, 12H), 0.89 (t, *J* = 7.1 Hz, 3H), 0.49 – 0.39 (m, 1H).

$^{13}C$ NMR (101 MHz, $CDCl_3$) δ 167.2, 167.0, 83.3, 63.3, 62.9, 46.3, 28.8, 25.3, 25.1, 25.0, 14.6.

$^{11}B$ NMR (128 MHz, $CDCl_3$) δ 34.9, 13.8.

HRMS $[M+Na]^+$ calculated for $C_{15}H_{27}B_2NNaO_6$ = 362.1917, found 362.1918.

**(1-Hydroxybutyl)boronic acid MIDA ester (28)**

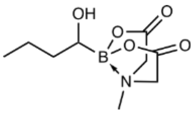

Compound **28** was prepared according to general procedure C from **27** (149 mg, 440 µmol, 1.0 eq.). The crude product was purified by flash column chromatography (gradient of 80 – 100% EtOAc in cyclohexane) to obtain the desired product **28** (33 mg, 144 µmol, 33%) as a colourless residue.

$^{1}H$ NMR (500 MHz, $CD_3CN$) δ 3.92 (d, *J* = 17.2 Hz, 1H), 3.90 (d, *J* = 16.2 Hz, 1H), 3.81 (d, *J* = 16.2 Hz, 1H), 3.79 (d, *J* = 17.2 Hz, 1H), 3.31 – 3.22 (m, 1H), 3.02 (s, 3H), 2.32 (d, *J* = 5.0 Hz, 1H), 1.59 – 1.42 (m, 3H), 1.41 – 1.26 (m, 1H), 0.93 (t, *J* = 7.2 Hz, 3H).

$^{13}C$ NMR (125 MHz, $CD_3CN$) δ 169.9, 169.0, 63.2, 62.9, 46.1, 36.4, 20.1, 14.3.

$^{11}B$ NMR (161 MHz, $CD_3CN$) δ 11.1.

HRMS $[M+Na]^+$ calculated for $C_9H_{16}BNNaO_5$ = 252.1014, found 252.1017.

**MIDA butyrylboronate (9c)**

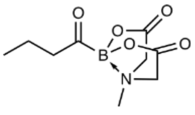

Compound **28** (32 mg, 140 µmol, 1.0 eq.) was dissolved in acetonitrile (8.2 mL) and DMP (89 mg, 210 µmol, 1.5 eq.) added. The reaction was stirred at room temperature for 30 minutes, then filtered over a pad of celite and the filtrate concentrated under reduced pressure. The crude material was purified by flash column chromatography (gradient of 60 to 100% EtOAc in cyclohexane, dry-load on celite) to obtain the desired product **9c** (21 mg, 93 µmol, 66%) as a white solid.

$^{1}H$ NMR (500 MHz, $CD_3CN$) δ 4.02 (d, *J* = 16.9 Hz, 2H), 3.88 (d, *J* = 16.9 Hz), 2.80 (s, 3H), 2.61 (t, *J* = 7.2 Hz, 2H), 1.52 (qt, *J* = 7.4, 7.2 Hz, 2H), 0.87 (t, *J* = 7.4 Hz, 3H).

$^{13}C$ NMR (125 MHz, $CD_3CN$) δ 168.1, 62.0, 46.4, 15.2, 13.1.

$^{11}B$ NMR (161 MHz, $CD_3CN$) δ 4.2.

HRMS $[M+Na]^+$ calculated for $C_9H_{14}BNNaO_5$ = 250.0857, found 250.0856.

**2.3.5 Synthesis of MIDA crotonylboronate 9d**

**2,2'-(But-3-ene-1,1-diyl)bis(4,4,5,5-tetramethyl-1,3,2-dioxaborolane) (29)**

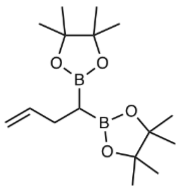

Compound **29** was prepared according to general procedure A, starting from bis[(pinacolato)boryl]methane (400 mg, 1.49 mmol, 1.0 eq.) and allyl bromide (142 µL, 1.64 mmol, 1.1 eq.). The reaction was stirred for 4 hours after the addition of allyl bromide. The crude product was purified by flash column chromatography (gradient of 0 to 30% EtOAc in hexane) to obtain the desired product **29** (354 mg, 1.15 mmol, 77%) as a colourless oil.

The recorded NMR data is in accordance with the literature[11].

$^1H$ NMR (500 MHz, $CDCl_3$) δ 5.92 – 5.83 (m, 1H), 5.02 – 4.96 (m, 1H), 4.89 – 4.84 (m, 1H), 2.33 – 2.26 (m, 2H), 1.23 (s, 12H), 1.21 (s, 12H), 0.84 (t, $J$ = 8.0 Hz, 1H).

$^{13}C$ NMR (125 MHz, $CDCl_3$) δ 140.9, 113.3, 83.2, 29.8, 25.0, 24.7.

$^{11}B$ NMR (161 MHz, $CDCl_3$) δ 34.1.

HRMS $[M+Na]^+$ calculated for $C_{16}H_{30}B_2NaO_4$ = 331.2222, found 331.2224.

**(1-(4,4,5,5-Tetramethyl-1,3,2-dioxaborolan-2-yl)but-3-en-1-yl)boronic acid MIDA ester (30)**

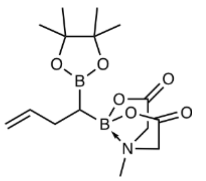

Compound **30** was prepared according to general procedure B from **29** (344 mg, 1.12 mmol, 1.0 eq.). The crude product was purified by flash column chromatography (gradient of 80 – 100% EtOAc in cyclohexane) to obtain the desired product **30** (242 mg, 718 µmol, 64%) as a white solid.

The recorded NMR data is in accordance with the literature[11].

$^1H$ NMR (500 MHz, $CDCl_3$) δ 6.02 – 5.89 (m, 1H), 5.06 – 4.97 (m, 1H), 4.93 – 4.86 (m, 1H), 3.94 (d, $J$ = 16.0 Hz, 1H), 3.83 (d, $J$ = 16.6 Hz, 1H), 3.79 (d, $J$ = 16.0 Hz, 1H), 3.77 (d, $J$ = 16.6 Hz, 1H), 3.03 (s, 3H), 2.41 – 2.27 (m, 2H), 1.21 (s, 12H), 0.56 – 0.48 (m, 1H).

$^{13}C$ NMR (125 MHz, $CDCl_3$) δ 167.4, 167.1, 140.9, 113.8, 83.5, 63.2, 62.8, 46.4, 30.6, 25.2, 25.0.

$^{11}B$ NMR (161 MHz, $CDCl_3$) δ 33.8, 13.5.

HRMS $[M+Na]^+$ calculated for $C_{15}H_{25}B_2NNaO_6$ = 360.176, found 360.1764.

**(*E*)-(1-(4,4,5,5-tetramethyl-1,3,2-dioxaborolan-2-yl)but-2-en-1-yl)boronic acid MIDA ester (31)**

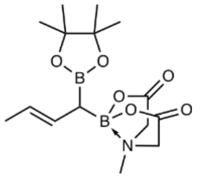

Compound **31** was prepared according to a literature procedure.[11] The substrate **30** (224 mg, 665 µmol, 1.0 eq.) and the catalyst acetonitrile(cyclopentadienyl)[2-(di-isopropylposphino)-4-(*t*-butyl)-1-Me-1-H-imidazole]Ru(II) (8.1 mg, 13.3 µmol, 0.02 eq.) were combined in a flask and set under inert atmosphere. DCM (anhydrous, 2.3 mL) was added and the reaction stirred at room temperature for 17 hours. Then, the reaction mixture was directly charged on a small silica column and purified (isocratic, 100% EtOAc) to obtain the desired product **31** (224 mg, 665 µmol, quant.) as a white foam.

The recorded NMR data is in accordance with the literature[11].

$^{1}$H NMR (500 MHz, $CD_3CN$) δ 5.55 (ddq, *J* = 15.4, 9.5, 1.6 Hz, 1H), 5.37 (dqd, *J* = 15.4, 6.4, 0.7 Hz, 1H), 3.79 – 3.68 (m, 4H), 2.99 (s, 3H), 1.64 (dd, *J* = 6.4, 1.6 Hz, 3H), 1.57 (d, *J* = 9.5 Hz, 1H), 1.25 (s, 12H).

$^{13}$C NMR (125 MHz, $CD_3CN$) δ 167.1, 166.7, 128.1, 125.3, 83.5, 62.9, 45.8, 24.9, 18.5.

$^{11}$B NMR (161 MHz, $CD_3CN$) δ 32.8, 12.8.

HRMS $[M+Na]^+$ calculated for $C_{15}H_{25}B_2NNaO_6$ = 360.176, found 360.1761.

**(*E*)-(1-hydroxybut-2-en-yl)boronic acid MIDA ester (32)**

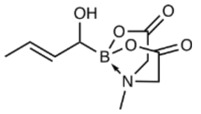

Compound **32** was prepared according to general procedure C from **31** (220 mg, 653 µmol, 1.0 eq.). The crude product was purified by flash column chromatography (gradient of 80 – 100% EtOAc in cyclohexane) to obtain the desired product **32** (124 mg, 546 µmol, 84%) as a colourless residue.

The recorded NMR data is in accordance with the literature[11].

$^{1}$H NMR (500 MHz, $CD_3CN$) δ 5.68 – 5.50 (m, 2H), 3.95 – 3.78 (m, 5H), 3.03 (s, 3H), 2.53 (s, 1H), 1.72 – 1.66 (m, 3H).

$^{13}$C NMR (125 MHz, $CD_3CN$) δ 169.7, 169.0, 133.6, 124.3, 63.2, 63.1, 46.3, 18.0.

$^{11}$B NMR (161 MHz, $CD_3CN$) δ 10.5.

HRMS $[M+Na]^+$ calculated for $C_9H_{14}BNNaO_5$ = 250.0857, found 250.0859.

**MIDA crotonylboronate (9d)**

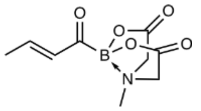

Substrate **32** (100 mg, 440 µmol, 1.0 eq.) was dissolved MeCN (17.5 mL). DMP (280 mg, 661 µmol, 1.5 eq.) was added. The reaction was stirred at room temperature for 1 hour. The reaction was filtered and the filtrate concentrated. The crude product was purified by flash column chromatography (gradient of 0 to 45% MeCN in DCM, dry-load on celite) to obtain the desired product **9d** (60.3 mg, 268 µmol, 61%) as a pale yellow foamy solid.

The recorded NMR data is in accordance with the literature[11].

$^{1}$H NMR (500 MHz, $CD_3CN$) δ 7.14 (dq, *J* = 16.0, 6.8 Hz, 1H), 6.33 (dq, *J* = 16.0, 1.6 Hz, 1H), 4.02 (d, *J* = 16.8 Hz, 2H), 3.90 (d, *J* = 16.8 Hz, 2H), 2.82 (s, 3H), 1.91 (dd, *J* = 6.8, 1.6 Hz, 3H).

$^{13}$C NMR (125 MHz, $CD_3CN$) δ 169.0, 146.4, 137.5, 62.8, 47.3, 19.0.

$^{11}$B NMR (161 MHz, $CD_3CN$) δ 4.9.

HRMS $[M+Na]^+$ calculated for $C_9H_{12}BNNaO_5$ = 248.0701, found 248.0701.

##### 2.3.6 Synthesis of MIDA benzoylboronate 9e

**(1-Phenylvinyl)boronic acid MIDA ester (33)**

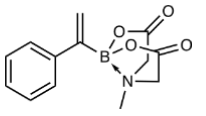

Compound **33** was prepared according to general procedure B from 1-phenylvinylboronic acid (250 mg, 1.69 mmol, 1.0 eq.). The crude product was purified by flash column chromatography (gradient of 60 – 100% EtOAc in hexane) to obtain the desired product **33** (242 mg, 934 µmol, 55%) as an orange solid.
$^{1}$H NMR (500 MHz, $CD_3CN$) δ 7.38 – 7.30 (m, 4H), 7.29 – 7.23 (m, 1H), 5.71 (s, 2H), 3.94 (d, *J* = 17.0 Hz, 2H), 3.62 (d, *J* = 17.0 Hz, 2H), 2.62 (s, 3H).

$^{13}$C NMR (125 MHz, $CD_3CN$) δ 169.2, 145.1, 129.3, 128.6, 128.5, 127.6, 62.7, 47.8.

$^{11}$B NMR (161 MHz, $CD_3CN$) δ 10.6.

HRMS $[M+Na]^+$ calculated for $C_{13}H_{14}BNNaO_4$ = 282.0908, found 282.0915.

**(1,2-Dihydroxy-1-phenylethyl)boronic acid MIDA ester (34)**

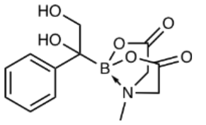

Compound **34** was prepared according to general procedure D from compound **33** (255 mg, 984 µmol, 1.0 eq.). The crude product was purified by flash column chromatography (gradient from 0 to 50% MeCN in DCM) to obtain the desired product **34** (270 mg, 921µmol, 94%) as an off-white solid.

The recorded NMR data is in accordance with the literature[12].

$^{1}$H NMR (500 MHz, $CD_3CN$) δ 7.52 – 7.46 (m, 2H), 7.36 – 7.30 (m, 2H), 7.22 – 7.17 (m, 1H), 3.94 – 3.81 (m, 5H), 3.44 (d, *J* = 17.0 Hz, 1H), 3.29 (s, 1H), 2.67 (s, 3H), 2.53 (dd, *J* = 6.9, 4.9 Hz, 1H), 1.96 (s, 1H).

$^{13}$C NMR (125 MHz, $CD_3CN$) δ 169.2, 169.0, 145.9, 128.7, 126.8, 126.6, 69.3, 63.7, 63.3, 47.6.

$^{11}$B NMR (161 MHz, $CD_3CN$) δ 10.2.

HRMS $[M+Na]^+$ calculated for $C_{13}H_{16}BNNaO_6$ = 316.0963, found 316.0957.

**MIDA benzoylboronate (9e)**

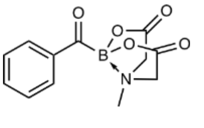

Compound **9e** was prepared according to general procedure E from compound **34** (192 mg, 655 µmol, 1.0 eq.). The reaction was stirred for 10 minutes. The desired product **9e** (132 mg, 506 µmol, 77%) was obtained as a purple solid without the need for further purification.

The recorded NMR data is in accordance with the literature[12].

$^{1}$H NMR (500 MHz, $CD_3CN$) δ 8.09 – 8.03 (m, 2H), 7.66 – 7.58 (m, 1H), 7.57 – 7.49 (m, 2H), 4.10 (d, $J$ = 16.8 Hz, 2H), 3.99 (d, $J$ = 16.8 Hz, 2H), 2.94 (s, 3H).

$^{13}$C NMR (125 MHz, $CD_3CN$) δ 168.9, 134.1, 129.6, 129.1, 62.9, 47.4.

$^{11}$B NMR (161 MHz, $CD_3CN$) δ 5.37.

HRMS $[M+Na]^+$ calculated for $C_{12}H_{12}BNNaO_5$ = 284.0701, found 284.07.

**2.3.7 Synthesis of MIDA 4-oxononanoylboronate 9f**

**1,1-bis(4,4,5,5-tetramethyl-1,3,2-dioxaborolan-2-yl)nonan-4-one (35)**

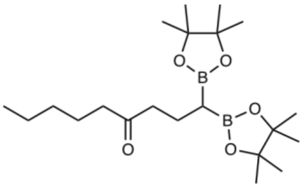

Compound **35** was prepared according to general procedure A from bis[(pinacolato)boryl]methane (400 mg, 1.49 mmol, 1.0 eq.) and 1-octen-3-one (249 µL, 1.64 mmol, 1.1 eq.). The reaction was stirred for 5.5 hours after addition of the electrophile. The crude product was purified by flash column chromatography (gradient of 0 to 20% EtOAc in cyclohexane) to obtain the desired product **35** (248 mg, 629 µmol, 42%) as a colourless oil.

$^{1}$H NMR (500 MHz, $CDCl_3$) δ 2.42 – 2.34 (m, 4H), 1.85 – 1.77 (m, 2H), 1.62 – 1.50 (m, 2H), 1.37 – 1.16 (m, 28H), 0.88 (t, $J$ = 7.1 Hz, 3H), 0.69 (t, $J$ = 7.8 Hz, 1H).

$^{13}$C NMR (125 MHz, $CDCl_3$) δ 211.9, 83.2, 45.3, 42.8, 31.6, 25.0, 24.7, 23.8, 22.6, 20.5, 14.1.

$^{11}$B NMR (161 MHz, $CDCl_3$) δ 34.0.

HRMS $[M+H]^+$ calculated for $C_{21}H_{41}B_2O_5$ = 395.3135, found 395.3137.

**(4-Oxo-1-(4,4,5,5-tetramethyl-1,3,2-dioxaborolan-2-yl)nonyl)boronic acid MIDA ester (36)**

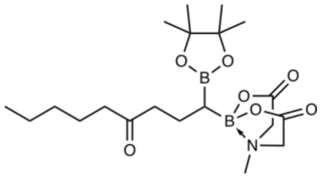

Compound **36** was prepared according to general procedure B from **35** (228 mg, 578 µmol, 1.0 eq.). The crude product was purified by flash column chromatography (gradient of 60 – 100% EtOAc in cyclohexane) to obtain the desired product **36** (64 mg, 151 µmol, 26%) as a pale-yellow solid.

$^{1}$H NMR (500 MHz, $CDCl_3$) δ 3.97 (d, $J$ = 16.0 Hz, 1H), 3.83 (d, $J$ = 16.4 Hz, 1H), 3.76 (d, $J$ = 16.4 Hz, 1H), 3.72 (d, $J$ = 16.0 Hz, 1H), 3.07 (s, 3H), 2.65 – 2.42 (m, 2H), 2.41 – 2.35 (m, 2H), 1.88 – 1.75 (m, 2H), 1.60 – 1.51 (m, 2H), 1.38 – 1.19 (m, 16H), 0.88 (t, $J$ = 7.0 Hz, 3H), 0.50 – 0.43 (m, 1H).

$^{13}$C NMR (125 MHz, $CDCl_3$) δ 212.5, 167.0, 83.5, 63.4, 63.0, 46.4, 44.3, 43.0, 31.6, 25.1, 24.9, 23.8, 22.6, 20.4, 14.1.

$^{11}$B NMR (161 MHz, $CDCl_3$) δ 35.7, 14.1.

HRMS $[M+Na]^+$ calculated for $C_{20}H_{35}B_2NNaO_7$ = 446.2492, found 446.2494.

**(1-Hydroxy-4-oxononyl)boronic acid MIDA ester (37)**

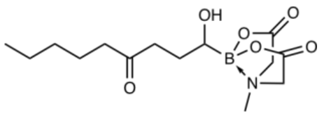

Compound **37** was prepared according to general procedure C from **36** (60.4 mg, 143 µmol, 1.0 eq.). The crude product was purified by flash column chromatography (gradient of 80 – 100% EtOAc in cyclohexane) to obtain the desired product **37** (31.5 mg, 101 µmol, 70%) as a colourless residue.

$^{1}$H NMR (500 MHz, $CD_3CN$) δ 3.98 – 3.71 (m, 4H), 3.25 – 3.17 (m, 1H), 3.07 – 2.96 (m, 3H), 2.63 (d, $J$ = 5.0 Hz, 1H), 2.59 – 2.52 (m, 1H), 2.47 – 2.40 (m, 1H), 2.11 – 1.21 (m, 10H), 0.93 – 0.84 (m, 3H).

$^{13}$C NMR (125 MHz, $CD_3CN$) δ 212.9, 170.0, 169.8, 168.9, 168.7, 63.21, 63.0, 62.9, 62.7, 46.3, 46.2, 43.2, 41.0, 40.9, 40.5, 38.6, 38.0, 32.9, 32.1, 28.3, 27.1, 27.0, 25.1, 25.0, 24.2, 23.3, 23.2, 14.3, 14.2.

$^{11}$B NMR (161 MHz, $CD_3CN$) δ 11.0.

HRMS $[M+Na]^+$ calculated for $C_{14}H_{24}BNNaO_6$ = 336.1589, found 336.1588.

**MIDA 4-oxononanoylboronate (9f)**

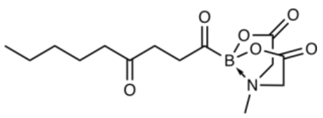

The substrate **37** (28 mg, 89 µmol, 1.0 eq.) was dissolved in acetonitrile (5.3 mL) and DMP (57 mg, 134 µmol, 1.5 eq.) added. The reaction was stirred at room temperature for 30 minutes, then filtered over a pad of celite. The filtrate was concentrated and the obtained residue purified by flash column chromatography (gradient from 60 to 100% EtOAc in cyclohexane, dry-load on celite) to obtain the desired product **9f** (25 mg, 80 µmol, 90%) as a colourless residue.

$^{1}$H NMR (500 MHz, $CD_3CN$) δ 4.03 (d, *J* = 16.9 Hz, 2H), 3.89 (d, *J* = 16.9 Hz, 2H), 2.85 – 2.82 (m, 2H), 2.81 (s, 3H), 2.65 – 2.60 (m, 2H), 2.46 – 2.40 (m, 2H), 1.55 – 1.46 (m, 2H), 1.37 – 1.20 (m, 4H), 0.88 (t, *J* = 7.2 Hz, 3H).

$^{13}$C NMR (125 MHz, $CD_3CN$) δ 210.7, 169.0, 63.0, 47.5, 42.9, 41.0, 35.6, 32.0, 24.3, 23.1, 14.2.

$^{11}$B NMR (161 MHz, $CD_3CN$) δ 4.3.

HRMS $[M+Na]^+$ calculated for $C_{14}H_{22}BNNaO_6$ = 334.1432, found 334.1433.

##### 2.3.8 Synthesis of MIDA β-hydroxybutyrylboronate 9g

**(*R*)-4-Pentyn-2-ol (38)**

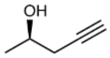

A 100 mL round-bottom flask was charged with lithium acetylide-ethylenediamine complex (2.22 g, 24.1 mmol, 1.4 eq.) and placed under argon atmosphere. DMSO (anhydr., 17.3 mL) was added and the flask cooled in an ice bath. (*R*)-(+)-propylene oxide (1.21 mL, 17.2 mmol, 1.0 eq.) was added dropwise. After the addition, the cooling bath was removed and the reaction stirred overnight at room temperature. Cold water (20 mL) was added to quench the reaction. The mixture was extracted with DCM (3 x 70 mL). The combined organic layers were washed with water (2 x 50 mL) and brine (1 x 50 mL), then dried over anhydrous sodium sulfate, filtered and concentrated under reduced pressure. (*R*)-4-pentyn-2-ol (**38**, 700 mg, 8.3 mmol, 48%) was obtained as a pale-yellow liquid and used in the next step without further purification.

The recorded NMR data is in accordance with the literature[15].

$^{1}$H NMR (500 MHz, $CDCl_3$) δ 4.03 – 3.93 (m, 1H), 2.41 (ddd, *J* = 16.6, 5.0, 2.7 Hz, 1H), 2.32 (ddd, *J* = 16.6, 6.6, 2.7 Hz, 1H), 2.07 (t, *J* = 2.6 Hz, 1H), 1.92 (d, *J* = 4.8 Hz, 1H), 1.27 (d, *J* = 6.2 Hz, 3H).

$^{13}$C NMR (125 MHz, $CDCl_3$) δ 81.0, 71.0, 66.4, 29.1, 22.4.

**(*R*)-*tert*-butyldimethyl(pent-4-yn-2-yloxy)silane (39)**

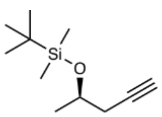

(*R*)-4-pentyn-2-ol (**38**, 987 mg, 11.7 mmol, 1.0 eq.) was dissolved in anhydrous DCM (26 mL) and imidazole (2.4 g, 35.2 mmol, 3.0 eq.) added, followed by *tert*-butylchlorodimethylsilane (2.65 g, 17.6 mmol, 1.5 eq.). The reaction was stirred at room temperature for 3 hours. Then, the reaction was quenched by addition of water (40 mL). The mixture was extracted with DCM (3 x 50 mL). The combined organic layers were washed with brine (40 mL), dried over anhydrous sodium sulfate, filtered and concentrated under reduced pressure. The crude product was purified by flash column chromatography (pentane) to obtain the desired product **39** (1.43 g, 7.2 mmol, 61%) as a colourless liquid.

The recorded NMR data is in accordance with the literature.[15]

$^{1}$H NMR (500 MHz, $CDCl_3$) δ 4.02 – 3.92 (m, 1H), 2.35 (ddd, *J* = 16.5, 5.5, 2.7 Hz, 1H), 2.24 (ddd, *J* = 16.5, 7.1, 2.7 Hz, 1H), 1.97 (t, *J* = 2.7 Hz, 1H), 1.24 (d, *J* = 6.0 Hz, 3H), 0.89 (s, 9H), 0.08 (s, 3H), 0.07 (s, 3H).

$^{13}$C NMR (125 MHz, $CDCl_3$) δ 82.1, 69.8, 67.7, 29.5, 26.0, 23.4, 18.3, -4.5, -4.6.

**(*R*)-*tert*-butyldimethyl((4-(4,4,5,5-tetramethyl-1,3,2-dioxaborolan-2-yl)pent-4-en-2-yl)oxy)silane (40)**

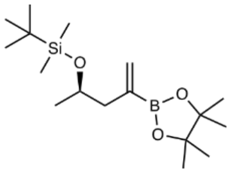

Compound **40** was prepared according to a literature procedure.[16] A heat-dried reaction vial under inert atmosphere was charged with bis(pinacolato)diboron (1.98 g, 7.81 mmol, 1.1eq.), tricyclohexylphosphine (199 mg, 710 µmol, 0.1 eq.) and $Pd(OAc)_2$ (80 mg, 355 µmol, 0.05 eq.) under a light counterflow of argon. The vial was set under inert atmosphere by evacuating and refilling with argon (three times). To the reaction vial was added anhydrous and degassed toluene (16.6 mL). *Anhydrous toluene was degassed in a heat-dried reaction vial under inert atmosphere by purging with argon while ultrasonicating for 10 minutes.* Then, the substrate **39** (1.41 g, 7.1 mmol, 1.0 eq.), bromobenzene (827 µL, 7.81 mmol, 1.1 eq.) and degassed TFE (1.04 mL, 14.2 mmol, 2.0 eq.) were added in that order. *TFE was degassed as described above for toluene.* The vial was capped and placed in a preheated heat-block at 60 °C and stirred for 3 hours. Then, the reaction was cooled down to room temperature, filtered over a pad of celite and the filter cake washed with $Et_2O$. The filtrate was concentrated under reduced pressure. The crude product was purified by flash column chromatography on silica gel (gradient from 0 to 5% EtOAc in hexane) to obtain the desired product **40** (1.31 g, 4.03 mmol, 67%) as a brown liquid.

$^1$H NMR (500 MHz, $CDCl_3$) δ 5.85 – 5.81 (m, 1H), 5.66 – 5.60 (m, 1H), 3.99 – 3.88 (m, 1H), 2.41 – 2.33 (m, 1H), 2.23 – 2.15 (m, 1H), 1.26 (s, 6H), 1.25 (s, 6H), 1.09 (d, *J* = 6.1 Hz, 3H), 0.88 (s, 9H), 0.05 (s, 3H), 0.04 (s, 3H).

$^{13}$C NMR (125 MHz, $CDCl_3$) δ 132.2, 83.5, 68.4, 46.5, 26.1, 25.0, 24.9, 23.4, 18.4, -4.4, -4.4.

$^{11}$B NMR (161 MHz, $CDCl_3$) δ 29.9.

HRMS $[M+Na]^+$ calculated for $C_{17}H_{35}BNaO_3Si$ = 349.2341, found 349.2343.

**(*R*)-(4-((*tert*-butyldimethylsilyl)oxy)pent-1-en-2-yl)boronic acid MIDA ester (41)**

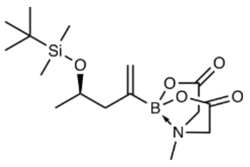

Compound **41** was prepared according to a procedure from the literature.[12] Substrate **40** (1.3 g, 3.98 mmol, 1.0 eq.) was dissolved in acetone (26.5 mL) and water (13.3 mL). Ammonium acetate (951 mg, 12.3 mmol, 3.1 eq.) and sodium periodate (2.64 g, 12.3 mmol, 3.1 eq.) were added to the solution. The reaction was heated to reflux for two hours, then allowed to cool down to room temperature. Aqueous sodium thiosulfate solution (0.5 M, 50 mL) was added and the mixture extracted with EtOAc (3 x 70 mL). The combined organic layers were washed with brine, dried over anhydrous sodium sulfate, filtered and concentrated under reduced pressure. The obtained residue was set under argon atmosphere by purging the flask with argon for 5 minutes. Anhydrous toluene (30 mL) and anhydrous DMSO (10 mL) was added, followed by methyliminodiacetic acid (1.76 g, 11.9 mmol, 3.0 eq.). A reflux condenser was attached and the reaction was heated to 105 °C for 2 hours. After cooling to room temperature, 30% brine (50 mL) was added and the mixture extracted with EtOAc (3 x 80 mL). The combined organic layers were dried over anhydrous sodium sulfate, filtered and concentrated under reduced pressure. The crude product was purified by flash column chromatography (gradient from 50 to 100% EtOAc in hexane) to obtain the desired product **41** (658 mg, 1.85 mmol, 47%) as a light brown sticky solid.

$^1$H NMR (500 MHz, $CD_3CN$) δ 5.56 – 5.47 (m, 1H), 5.45 – 5.35 (m, 1H), 4.09 – 4.01 (m, 1H), 3.94 (d, *J* = 17.0 Hz, 2H), 3.79 (d, *J* = 17.0, 1.0 Hz, 2H), 2.75 (s, 3H), 2.26 – 2.17 (m, 1H), 2.11 – 2.03 (m, 1H), 1.10 (d, *J* = 6.0 Hz, 3H), 0.87 (s, 9H), 0.06 (s, 3H), 0.04 (s, 3H).

$^{13}$C NMR (125 MHz, $CD_3CN$) δ 169.3, 127.0, 68.6, 62.5, 47.5, 46.6, 26.3, 24.1, -4.3.

$^{11}B$ NMR (161 MHz, $CD_3CN$) δ 10.5.

HRMS $[M+Na]^+$ calculated for $C_{16}H_{30}BNNaO_5Si$ = 387.1879, found 378.1875.

**((4*R*)-4-((*tert*-butyldimethylsilyl)oxy)-1,2-dihydroxypentan-2-yl)boronic acid MIDA ester (42)**

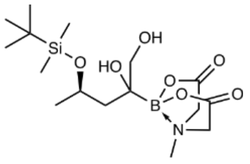

Compound **42** was prepared according to general procedure D from compound **41** (647 mg, 1.82 mmol, 1.0 eq.). The reaction was stirred for 4.5 hours before TLC analysis showed complete consumption of the starting material. The crude product was purified by flash column chromatography (gradient from 0 to 40% MeCN in DCM) to obtain the desired product **42** (583 mg, 1.5 mmol, 82%) as an orange solid.

$^1H$ NMR (500 MHz, $CD_3CN$) δ 4.46 – 4.30 (m, 1H), 4.04 – 3.79 (m, 4H), 3.77 – 3.41 (m, 2H), 3.15 – 3.09 (m, 3H), 3.02 – 2.96 (m, 1H), 2.98 – 2.84 (m, 1H), 1.88 – 1.58 (m, 2H), 1.22 – 1.15 (m, 3H), 0.93 – 0.84 (m, 9H), 0.16 – 0.06 (m, 6H).

$^{13}C$ NMR (125 MHz, $CD_3CN$) δ 169.7, 169.2, 69.1, 68.7, 68.1, 67.3, 64.0, 63.9, 63.3, 63.3, 47.6, 45.0, 43.9, 26.2, 26.1, 25.9, 18.5, -3.2, -3.4, -4.5, -4.5.

$^{11}B$ NMR (161 MHz, $CD_3CN$) δ 10.7.

HRMS $[M+Na]^+$ calculated for $C_{16}H_{32}BNNaO_7Si$ = 412.1933, found 412.1928.

**(*R*)-(3-((*tert*-butyldimethylsilyl)oxy)butanoyl)boronic acid MIDA ester (43)**

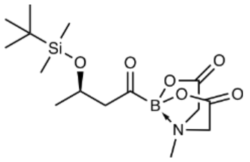

Compound **43** was prepared according to general procedure E from compound **42** (205 mg, 527 µmol, 1.0 eq.). The reaction was stirred for 30 minutes. The desired product **43** (47.5 mg, 133 µmol, 25%) was obtained as a white solid after purification by flash column chromatography (gradient from 0 to 40% MeCN in DCM).

$^1H$ NMR (500 MHz, $CD_3CN$) δ 4.43 – 4.35 (m, 1H), 4.02 (d, $J$ = 17.0 Hz, 2H), 3.87 (dd, $J$ = 17.0, 3.0 Hz, 2H), 2.89 (dd, $J$ = 16.9, 6.2 Hz, 1H), 2.80 (s, 3H), 2.63 (dd, $J$ = 16.9, 6.9 Hz, 1H), 1.12 (d, $J$ = 6.2 Hz, 3H), 0.85 (s, 9H), 0.07 (s, 3H), 0.04 (s, 3H).

$^{13}C$ NMR (125 MHz, $CD_3CN$) δ 168.9, 64.7, 63.0, 57.2, 47.4, 26.2, 24.5, 18.5, -4.4, -4.6.

$^{11}B$ NMR (161 MHz, $CD_3CN$) δ 4.0.

HRMS $[M+Na]^+$ calculated for $C_{15}H_{28}BNNaO_6Si$ = 380.1671, found 380.1673.

**MIDA β-hydroxybutyrylboronate (9g)**

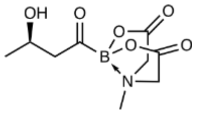

The substrate **43** (18 mg, 50 µmol, 1.0 eq.) was placed under inert atmosphere by purging the flask with argon for 5 minutes. 1 M TBAF in THF (504 µL, 504 µmol, 10.0 eq.) was added and the reaction stirred at room temperature for 6 hours. The reaction was diluted with THF (10 mL) and directly concentrated on celite. The crude product was purified by flash column chromatography (gradient from 0 to 70% MeCN in DCM, dry-load on celite) to obtain **9g** (5.8 mg, 24 µmol, 47%) as a white residue.

$^{1}$H NMR (500 MHz, $CD_3CN$) δ 4.27 – 4.16 (m, 1H), 4.02 (dd, *J* = 17.0, 0.8 Hz, 2H), 3.89 (dd, *J* = 17.0, 2.5 Hz, 2H), 2.82 (s, 3H), 2.76 (dd, *J* = 16.9, 8.0 Hz, 1H), 2.68 (dd, *J* = 16.9, 4.4 Hz, 1H), 1.11 (d, *J* = 6.2 Hz, 3H).

$^{13}$C NMR (125 MHz, $CD_3CN$) δ 169.0, 64.0, 63.1, 47.5, 23.6.

$^{11}$B NMR (161 MHz, $CD_3CN$) δ 4.1.

HRMS $[M+Na]^+$ calculated for $C_9H_{14}BNNaO_6$ = 266.0806, found 266.0811.

**2.3.9 Synthesis of MIDA malonylboronate 9h**

***Tert*-butyl 3,3-bis(4,4,5,5-tetramethyl-1,3,2-dioxaborolan-2-yl)propanoate (44)**

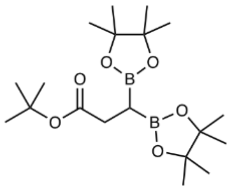

Compound **44** was prepared according to general procedure A, starting from bis[(pinacolato)boryl]methane (800 mg, 2.99 mmol, 1.0 eq.) and *tert*-butyl bromoacetate (485 µL, 3.28 mmol, 1.1 eq.). After the addition of the electrophile, the reaction was stirred for 2 hours. The crude product was purified by flash column chromatography (gradient of 0 to 20% EtOAc in hexane) to obtain the desired product **44** (960 mg, 2.51 mmol, 84%) as a colourless oil.

$^{1}$H NMR (500 MHz, $CDCl_3$) δ 2.49 (d, *J* = 8.4 Hz, 2H), 1.41 (s, 9H), 1.23 (s, 12H), 1.20 (s, 12H), 1.04 (t, *J* = 8.4 Hz, 1H).

$^{13}$C NMR (125 MHz, $CDCl_3$) δ 174.34, 83.24, 79.76, 31.87, 28.25, 25.01, 24.87, 24.60.

$^{11}$B NMR (161 MHz, $CDCl_3$) δ 33.6.

HRMS $[M+Na]^+$ calculated for $C_{19}H_{36}B_2NaO_6$ = 405.259, found 405.259.

**(3-(*Tert*-butoxy)-3-oxo-1-(4,4,5,5-tetramethyl-1,3,2-dioxaborolan-2-yl)propyl)boronic acid MIDA ester (45)**

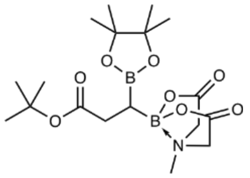

Compound **45** was prepared according to general procedure B from **44** (950 mg, 2.49 mmol, 1.0 eq.). The crude product was purified by flash column chromatography (gradient of 50 – 100% EtOAc in hexane) to obtain the desired product **45** (128 mg, 311 µmol, 13%) as a white solid.

$^{1}$H NMR (500 MHz, $CD_3CN$) δ 3.94 (d, *J* = 17.2 Hz, 1H), 3.91 (d, *J* = 16.8 Hz, 1H), 3.84 (d, *J* = 16.8 Hz, 1H), 3.80 (d, *J* = 17.2 Hz, 1H), 2.99 (s, 3H), 2.38 – 2.24 (m, 2H), 1.41 (s, 9H), 1.19 (s, 6H), 1.17 (s, 6H), 0.84 – 0.74 (m, 1H).

$^{13}$C NMR (125 MHz, $CD_3CN$) δ 175.1, 168.9, 168.7, 84.0, 80.3, 63.5, 63.2, 47.1, 33.2, 28.3, 25.2, 25.1.

$^{11}$B NMR (161 MHz, $CD_3CN$) δ 34.2, 12.8.

HRMS $[M+Na]^+$ calculated for $C_{18}H_{31}B_2NNaO_8$ = 434.2128, found 434.2139.

**(3-(*Tert*-butoxy)-1-hydroxy-3-oxopropyl)boronic acid MIDA ester (46)**

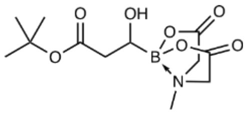

Compound **46** was prepared according to general procedure C from **45** (126 mg, 307 µmol, 1.0 eq.). The crude product was purified by flash column chromatography (gradient of 70 – 100% EtOAc in hexane) to obtain the desired product **46** (94 mg, 307 µmol, quant.) as a white solid.

$^{1}$H NMR (500 MHz, $CD_3CN$) δ 3.95 (d, *J* = 17.2 Hz, 1H), 3.94 (d, *J* = 16.4 Hz, 1H), 3.82 (d, *J* = 17.2 Hz, 1H), 3.81 (d, *J* = 16.4 Hz, 1H), 3.74 – 3.67 (m, 1H), 3.06 (s, 3H), 3.02 – 2.93 (m, 1H), 2.38 (d, *J* = 1.1 Hz, 1H), 2.36 (s, 1H), 1.45 (s, 9H).

$^{13}$C NMR (125 MHz, $CD_3CN$) δ 173.9, 169.6, 168.8, 81.3, 63.3, 63.1, 46.3, 39.5, 28.3.

$^{11}$B NMR (161 MHz, $CD_3CN$) δ 10.7.

HRMS $[M+Na]^+$ calculated for $C_{12}H_{20}BNNaO_7$ = 324.1225, found 324.1227.

**(3-(*Tert*-butoxy)-1-hydroxy-3-oxoprop-1-en-1-yl)boronic acid MIDA ester (47)**

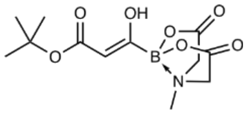

Substrate **46** (87 mg, 289 µmol, 1.0 eq.) was dissolved in acetonitrile (17 mL) and DMP (184 mg, 433 µmol, 1.5 eq.) added. The reaction was stirred at room temperature for 1.5 hours. Then, the reaction was filtered over a pad of celite and concentrated under reduced pressure. The crude material was purified by flash column chromatography (gradient from 50 to 100% EtOAc in hexane, dry-load on celite) to obtain the desired product **47** (63 mg, 211 µmol, 73%) as a white solid.

$^{1}H$ NMR (500 MHz, $CD_3CN$) δ 11.77 (s, 1H), 5.32 (s, 1H), 4.04 (d, *J* = 16.9 Hz, 2H), 3.88 (d, *J* = 16.9 Hz, 2H), 2.90 (s, 3H), 1.48 (s, 9H).

$^{13}C$ NMR (125 MHz, $CD_3CN$) δ 172.8, 168.9, 101.0, 82.0, 62.8, 47.6, 28.4.

$^{11}B$ NMR (161 MHz, $CD_3CN$) δ 8.0.

HRMS $[M+Na]^+$ calculated for $C_{12}H_{18}BNNaO_7$ = 322.1069, found 322.1069.

**MIDA malonylboronate (9h)**

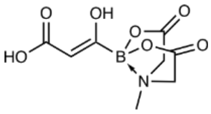

The substrate **47** (61 mg, 204 µmol, 1.0 eq.) was dissolved in DCM (1 mL) and TFA (468 µl, 6.1 mmol, 30 eq.) added. The reaction was stirred at room temperature for 30 minutes. Then, the reaction was concentrated under reduced pressure at 30 °C bath temperature. The crude residue was redissolved MeCN and DCM and concentrated again to obtain the desired product **9h** (39 mg, 160 µmol, 79%) as a white solid that did not require further purification.

$^{1}H$ NMR (500 MHz, $CD_3CN$) δ 12.00 (s, 1H), 5.44 (s, 1H), 4.05 (d, *J* = 16.9 Hz, 2H), 3.89 (d, *J* = 16.9 Hz, 2H), 2.90 (s, 3H).

$^{13}C$ NMR (125 MHz, $CD_3CN$) δ 173.7, 168.9, 98.5, 62.8, 47.6.

$^{11}B$ NMR (161 MHz, $CD_3CN$) δ 7.8.

HRMS $[M+Na]^+$ calculated for $C_8H_{10}BNNaO_7$ = 266.0443, found 266.0444.

**2.3.10 Synthesis of MIDA succinylboronate 9i**

***Tert*-butyl-4-pentynoate (48)**

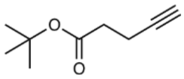

In a 50 mL round-bottom flask, 4-pentynoic acid (1.57 g, 16.0 mmol, 1.0 eq.) was dissolved in DCM (5.3 mL) and *tert*-butanol (3.04 mL, 32.1 mmol, 2.0 eq.) added, followed by DMAP (78 mg, 641 µmol, 0.04 eq.). DCC (3.64 g, 17.6 mmol, 1.1 eq.) was added to the mixture as a solution in DCM (5.3 mL). The reaction was stirred overnight at room temperature, then filtered and the filter cake washed with DCM. The filtrate was washed with aq. 0.5 M HCl (2 x 50 mL) and aq. sat. $NaHCO_3$ (2 x 50 mL), then dried over anhydrous sodium sulfate, filtered and concentrated under reduced pressure. The crude product was purified by flash column chromatography (gradient from 0 to 20% ether in pentane) to obtain *tert*-butyl-4-pentynoate (**48**, 1.86 g, 12.04 mmol, 75%) as a colourless oil.

The recorded NMR data is in accordance with the literature.[17]

$^{1}H$ NMR (500 MHz, $CDCl_3$) δ 2.47 – 2.43 (m, 4H), 1.97 – 1.95 (m, 1H), 1.45 (s, 9H).

$^{13}C$ NMR (125 MHz, $CDCl_3$) δ 171.2, 82.9, 81.0, 68.9, 34.6, 28.2, 14.6.

HRMS $[M+Na]^+$ calculated for $C_9H_{14}NaO_2$ = 177.0886, found 177.089.

**_Tert_-butyl 4-(4,4,5,5-tetramethyl-1,3,2-dioxaborolan-2-yl)pent-4-enoate (49)**

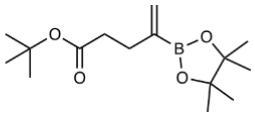

Compound **49** was prepared according to a literature procedure.[16] A heat-dried reaction vial under inert atmosphere was charged with bis(pinacolato)diboron (1.79 g, 7.06 mmol, 1.1eq.), tricyclohexylphosphine (180 mg, 642 µmol, 0.1 eq.) and $Pd(OAc)_2$ (72 mg, 321 µmol, 0.05 eq.) under a light counterflow of argon. The vial was set under inert atmosphere by evacuating and refilling with argon (three times). To the reaction vial was added anhydrous and degassed toluene (15 mL). *Anhydrous toluene was degassed in a heat-dried reaction vial under inert atmosphere by purging with argon while ultrasonicating for 10 minutes.* Then, *tert*-butyl-4-pentynoate (**48**, 1.08 mL, 6.42 mmol, 1.0 eq.), bromobenzene (744 µL, 7.06 mmol, 1.1 eq.) and degassed TFE (936 µL, 128 mmol, 2.0 eq.) were added in that order. *TFE was degassed as described above for toluene.* The vial was capped and placed in a preheated heat-block at 60 °C and stirred for 3 hours. Then, the reaction was cooled down to room temperature, filtered over a pad of celite and the filter cake washed with $Et_2O$. The filtrate was concentrated under reduced pressure. The crude product was purified by flash column chromatography on silica gel (gradient from 0 to 20% $Et_2O$ in pentane) to obtain the desired product **49** (401 mg, 1.42 mmol, 22%) as a yellow liquid.

$^1$H NMR (500 MHz, $CDCl_3$) δ 5.84 – 5.73 (m, 1H), 5.70 – 5.56 (m, 1H), 2.49 – 2.33 (m, 4H), 1.43 (s, 9H), 1.26 (s, 12H).

$^{13}$C NMR (125 MHz, $CDCl_3$) δ 173.0, 129.9, 83.6, 80.1, 35.2, 31.0, 28.3, 24.9.

$^{11}$B NMR (161 MHz, $CDCl_3$) δ 29.9.

HRMS $[M+Na]^+$ calculated for $C_{15}H_{27}BNaO_4$ = 305.1895, found 305.1892.

**(5-(_Tert_-butoxy)-5-oxopent-1-en-2-yl)boronic acid MIDA ester (50)**

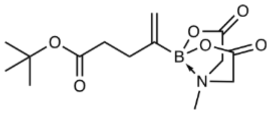

Compound **50** was prepared according to general procedure B from **49** (380 mg, 1.35 mmol, 1.0 eq.). The crude product was purified by flash column chromatography (gradient from 70 to 100% EtOAc in cyclohexane) to obtain the desired product **50** (369 mg, 1.19 mmol, 88%) as an off-white solid.

$^1$H NMR (500 MHz, $CDCl_3$) δ 5.54 – 5.49 (m, 1H), 5.49 – 5.44 (m, 1H), 3.89 (d, $J$ = 16.4 Hz, 2H), 3.78 (d, $J$ = 16.4 Hz, 2H), 2.87 (s, 3H), 2.50 – 2.44 (m, 2H), 2.33 – 2.26 (m, 2H), 1.42 (s, 9H).

$^{13}$C NMR (125 MHz, $CDCl_3$) δ 173.2, 167.6, 123.8, 80.5, 62.0, 46.7, 33.6, 29.2, 28.2.

$^{11}$B NMR (161 MHz, $CDCl_3$) δ 11.0.

HRMS $[M+Na]^+$ calculated for $C_{14}H_{22}BNNaO_6$ = 334.1432, found 334.1436.

**(5-(_Tert_-butoxy)-1,2-dihydroxy-5-oxopentan-2-yl)boronic acid MIDA ester (51)**

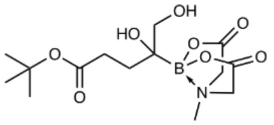

Compound **51** was prepared according to general procedure D from compound **50** (150 mg, 482 µmol, 1.0 eq.). The reaction was stirred for 1 hour before TLC analysis showed complete consumption of the starting material. The crude product was purified by flash column chromatography (gradient from 20 to 70% MeCN in DCM) to obtain the desired product **51** (134 mg, 388 µmol, 81%) as an off-white foamy solid.

$^{1}$H NMR (500 MHz, $CD_3CN$) δ 3.94 (dd, *J* = 16.6, 5.4 Hz, 2H), 3.88 (dd, *J* = 16.6, 2.5 Hz, 2H), 3.60 (d, *J* = 11.7 Hz, 1H), 3.47 (d, *J* = 11.7 Hz, 1H), 3.15 (s, 3H), 2.33 – 2.26 (m, 2H), 1.92 – 1.81 (m, 1H), 1.80 – 1.70 (m, 1H), 1.45 (s, 9H).

$^{13}$C NMR (125 MHz, $CD_3CN$) δ 174.6, 169.3, 80.6, 65.9, 63.6, 63.2, 47.4, 31.5, 31.1, 28.2.

$^{11}$B NMR (161 MHz, $CD_3CN$) δ 10.5.

HRMS $[M+Na]^+$ calculated for $C_{14}H_{24}BNNaO_8$ = 368.1487, found 368.1487.

**(4-(*Tert*-butoxy)-4-oxobutanoyl)boronic acid MIDA ester (52)**

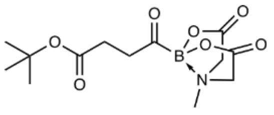

Compound **52** was prepared according to general procedure E from compound **51** (131 mg, 380 µmol, 1.0 eq.). The reaction was stirred for 10 minutes. The desired product **52** (107 mg, 342 µmol, 90%) was obtained as a pale brownish foamy solid an did not require any purification.
$^{1}$H NMR (500 MHz, $CD_3CN$) δ 4.03 (d, *J* = 16.9 Hz, 2H), 3.89 (d, *J* = 16.9 Hz, 2H), 2.89 – 2.84 (m, 2H), 2.81 (s, 3H), 2.41 – 2.37 (m, 2H), 1.39 (s, 9H).

$^{13}$C NMR (125 MHz, $CD_3CN$) δ 173.2, 169.0, 80.7, 63.0, 47.5, 42.1, 28.6, 28.2.

$^{11}$B NMR (161 MHz, $CD_3CN$) δ 4.3.

HRMS $[M+Na]^+$ calculated for $C_{13}H_{20}BNNaO_7$ = 336.1225, found 336.1229.

**MIDA succinylboronate (9i)**

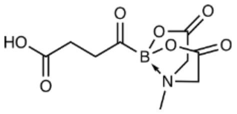

The substrate **52** (50 mg, 160 µmol, 1.0 eq.) was dissolved in DCM (1 mL) and TFA (367 µL, 4.79 mmol, 30 eq.) added. The reaction was stirred for 2 hours at room temperature. TLC (50% MeCN in DCM) indicated complete consumption of the starting material. The reaction was concentrated under reduced pressure at room temperature. The crude product was purified by flash column chromatography (gradient from 30 to 100% MeCN in DCM, dry-load on celite) to obtain the desired product **9i** (37 mg, 144 µmol, 90%) as a white solid.

Based on the NMR spectra, we assume that the obtained compound exists in two forms. In the structure that is displayed and one in which the succinyl carboxylic acid coordinates to boron, replacing one of the previous substituents.

$^{1}$H NMR (500 MHz, $CD_3CN$) δ 4.06 – 3.99 (m, 2H), 3.94 – 3.86 (m, 2H), 3.10 and 2.81 (s, 3H), 2.92 – 2.87, 2.69 – 2.60 and 2.54 – 2.41 (m, 4H).

$^{13}$C NMR (125 MHz, $CD_3CN$) δ 174.5, 169.0, 63.6, 63.0, 47.8, 47.5, 33.1, 28.0, 26.6.

$^{11}$B NMR (161 MHz, $CD_3CN$) δ 8.9, 4.3.

HRMS $[M+Na]^+$ calculated for $C_9H_{12}BNNaO_7$ = 280.0599, found 280.0606.

**2.3.11 Synthesis of MIDA glutarylboronate 9j**

***Tert*-butyl 4-iodobutanoate (53)**

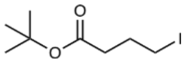

*Tert*-butyl 4-bromobutanoate (2.23 g, 10.0 mmol, 1.0 eq.) was dissolved in acetone (40 mL) and sodium iodide (3.0 g, 20.0 mmol, 2.0 eq.) added. The reaction was wrapped in aluminium foil and stirred at room temperature for 72 hours. Then, the reaction was filtered and the filtrate concentrated under reduced pressure. The residue was dissolved in EtOAc (150 mL) and washed with water (50 mL) and brine (50 mL). The organic layer was dried over anhydrous sodium sulfate, filtered and concentrated under reduced pressure to obtain *tert*-butyl 4-iodobutanoate (**53**, 2.37 g, 8.77 mmol, 88%) as a brown liquid which was used in the next step without further purification.

The recorded NMR data is in accordance with the literature.[18]

$^1$H NMR (500 MHz, $CDCl_3$) δ 3.23 (t, *J* = 6.8 Hz, 2H), 2.35 (t, *J* = 7.2 Hz, 2H), 2.13 – 2.05 (m, 2H), 1.45 (s, 9H).

***Tert*-butyl 5,5-bis(4,4,5,5-tetramethyl-1,3,2-dioxaborolan-2-yl)pentanoate (54)**

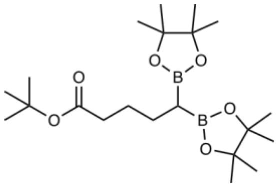

Compound **54** was prepared according to general procedure A, starting from bis[(pinacolato)boryl]methane (800 mg, 2.99 mmol, 1.0 eq.) and *tert*-butyl 4-iodobutanoate (**53**, 887 mg, 3.28 mmol, 1.1 eq.). The reaction was stirred for 2 hours after the addition of *tert*-butyl 4-iodobutanoate. The crude product was purified by flash column chromatography (gradient from 0 to 15% EtOAc in hexane). The desired product **54** (851 mg, 1.78 mmol, 60%, 86% purity) was obtained as a colourless oil containing some bis[(pinacolato)boryl]methane. The product was entered into the next reaction without further purification as products are easily separated after the next step.

$^1$H NMR (500 MHz, $CDCl_3$) δ 2.21 – 2.14 (m, 2H), 1.59 – 1.54 (m, 4H), 1.42 (s, 9H), 1.22 (s, 12H), 1.22 (s, 12H), 0.78 – 0.67 (m, 1H).

$^{13}$C NMR (125 MHz, $CDCl_3$) δ 173.4, 83.1, 79.8, 36.0, 28.3, 28.0, 25.4, 25.0, 24.7.

$^{11}$B NMR (161 MHz, $CDCl_3$) δ 33.6.

HRMS $[M+Na]^+$ calculated for $C_{21}H_{40}B_2NaO_6$ = 433.2903, 433.2907 found.

**(5-(*Tert*-butoxy)-5-oxo-1-(4,4,5,5-tetramethyl-1,3,2-dioxaborolan-2-yl)pentyl)boronic acid MIDA ester (55)**

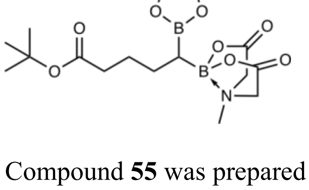

Compound **55** was prepared according to general procedure B from **54** (831 mg, 2.03 mmol, 1.0 eq.). The crude product was purified by flash column chromatography (gradient from 70 to 100% EtOAc in hexane) to obtain the desired product **55** (376 mg, 856 µmol, 52%) as a white solid.

$^{1}H$ NMR (500 MHz, $CD_3CN$) δ 3.90 (dd, *J* = 17.1, 14.4 Hz, 2H), 3.80 (dd, *J* = 17.8, 17.1 Hz, 2H), 2.93 (s, 3H), 2.21 – 2.09 (m, 2H), 1.71 – 1.60 (m, 1H), 1.55 – 1.33 (m, 3H), 1.41 (s, 9H), 1.20 (s, 12H), 0.46 – 0.36 (m, 1H).

$^{13}C$ NMR (125 MHz, $CD_3CN$) δ 173.8, 169.1, 168.9, 83.8, 80.3, 63.5, 63.4, 46.9, 36.5, 28.4, 28.3, 26.8, 25.2.

$^{11}B$ NMR (161 MHz, $CD_3CN$) δ 34.5, 13.2.

HRMS $[M+Na]^+$ calculated for $C_{20}H_{35}B_2NNaO_8$ = 462.2441, 462.2444 found.

**(5-(*Tert*-butoxy)-1-hydroxy-5-oxopentyl)boronic acid MIDA ester (56)**

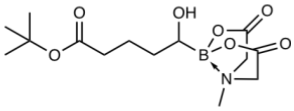

Compound **56** was prepared according to general procedure C from **55** (372 mg, 847 µmol, 1.0 eq.). The crude product was purified by flash column chromatography (gradient from 0 to 70% MeCN in DCM) to obtain **56** (240 mg, 729 µmol, 86%) as a white solid.

$^{1}H$ NMR (400 MHz, $CD_3CN$) δ 3.91 (d, *J* = 17.2 Hz, 1H), 3.90 (d, *J* = 16.2 Hz, 1H), 3.81 (d, *J* = 16.2 Hz, 1H), 3.78 (d, *J* = 17.2 Hz, 1H), 3.28 – 3.20 (m, 1H), 3.01 (s, 3H), 2.45 (d, *J* = 5.1 Hz, 1H), 2.22 (t, *J* = 7.3 Hz, 2H), 1.80 – 1.66 (m, 1H), 1.65 – 1.44 (m, 3H), 1.43 (s, 9H).

$^{13}C$ NMR (100 MHz, $CD_3CN$) δ 174.1, 169.9, 169.0, 80.5, 63.2, 62.9, 46.5, 35.9, 33.6, 28.3, 22.5.

$^{11}B$ NMR (128 MHz, $CD_3CN$) δ 11.0.

HRMS $[M+Na]^+$ calculated for $C_{14}H_{24}BNNaO_7$ = 352.1538, 352.1538 found.

**(5-(*tert*-butoxy)-5-oxopentanoyl)boronic acid MIDA ester (57)**

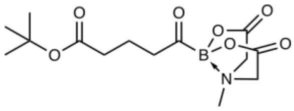

The substrate **56** (227 mg, 690 µmol, 1.0 eq.) was dissolved in acetonitrile (27 mL) and DMP (439 mg, 1.03 mmol, 1.5 eq.) added. After one hour, additional DMP (293 mg, 690 µmol, 1.0 eq.) added due to remaining starting material. After further 30 minutes, the reaction was filtered through a pad of celite and the filtrate concentrated under reduced pressure. The crude product was purified by flash column chromatography on silica (gradient from 0 to 60% MeCN in DCM, dry-load on celite) to obtain **57** (185 mg, 566 µmol, 82%) as a white solid.

$^{1}H$ NMR (500 MHz, $CD_3CN$) δ 4.02 (d, *J* = 16.9 Hz, 2H), 3.89 (d, *J* = 16.9 Hz, 2H), 2.81 (s, 3H), 2.67 (t, *J* = 7.1 Hz, 2H), 2.16 (t, *J* = 7.5 Hz, 2H), 1.76 – 1.68 (m, 2H), 1.42 (s, 9H).

$^{13}C$ NMR (125 MHz, $CD_3CN$) δ 173.5, 169.0, 80.6, 63.0, 47.4, 46.3, 35.3, 28.2, 18.4.

$^{11}B$ NMR (161 MHz, $CD_3CN$) δ 4.2.

HRMS $[M+NH_4]^+$ calculated for $C_{14}H_{26}BN_2O_7$ = 345.1828, 345.1828 found.

**MIDA glutarylboronate (9j)**

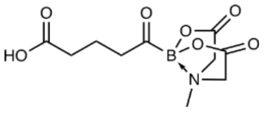

The substrate **57** (56 mg, 171 µmol, 1.0 eq.) was dissolved in DCM (1.2 mL) and TFA (393 µL, 5.14 mmol, 30.0 eq.) added. The reaction was stirred for 1.5 hours, then concentrated under reduced pressure. The crude product was purified by flash column chromatography (gradient from 20 to 70% MeCN in DCM + 0.5% acetic acid, dry load on celite) to obtain the desired product **9j** (38 mg, 140 µmol, 82%) as a white solid.

$^{1}$H NMR (500 MHz, $CD_3CN$) δ 4.02 (d, $J$ = 16.9 Hz, 2H), 3.89 (d, $J$ = 16.9 Hz, 2H), 2.81 (s, 3H), 2.69 (t, $J$ = 7.1 Hz, 2H), 2.25 (t, $J$ = 7.5 Hz, 2H), 1.78 – 1.71 (m, 2H).

$^{13}$C NMR (125 MHz, $CD_3CN$) δ 174.8, 169.0, 63.0, 47.4, 46.3, 33.3, 18.1.

$^{11}$B NMR (161 MHz, $CD_3CN$) δ 4.24.

HRMS $[M+NH_4]^+$ calculated for $C_{10}H_{14}BNNaO_7$ = 294.0756, 294.0757 found.

**2.3.12 Synthesis of MIDA itaconylboronate 9k**

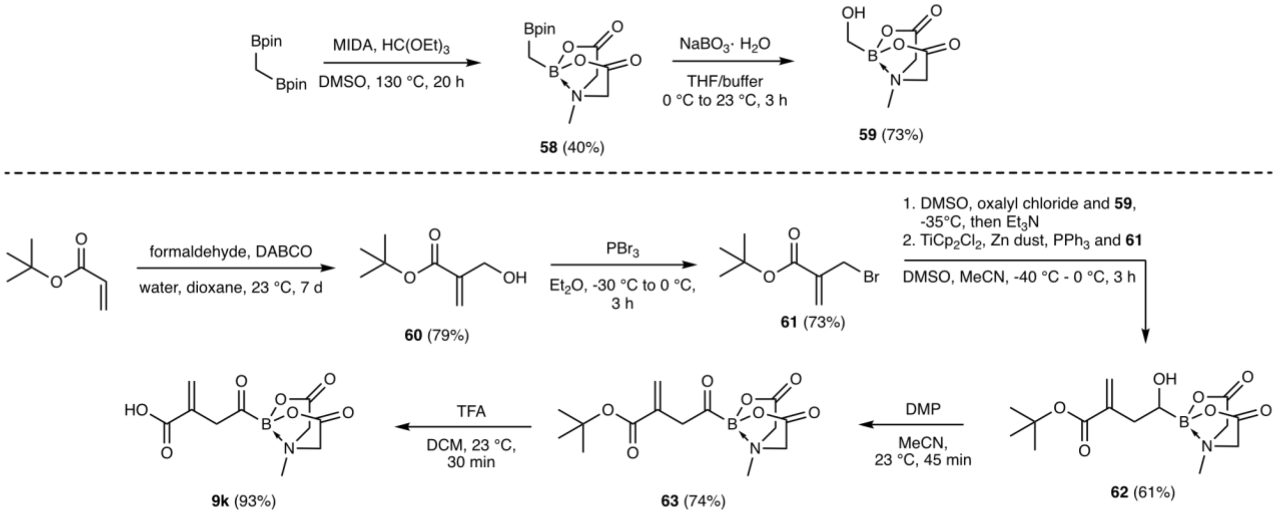

**Scheme S3:** Synthesis of MIDA itaconylboronate **9k**.

**((4,4,5,5-Tetramethyl-1,3,2-dioxaborolan-2-yl)methyl)boronic acid MIDA ester (58)**

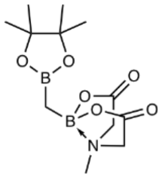

Compound **58** was prepared according to general procedure B from bis(4,4,5,5-tetramethyl-1,3,2-dioxaborolan-2-yl)methane (800 mg, 2.99 mmol, 1.0 eq.). The crude product was purified by flash column chromatography (gradient from 80 to 100% EtOAc in hexane) to obtain the desired product **58** (354 mg, 1.19 mmol, 40%) as a white solid.

The recorded NMR data is in accordance with the literature.[11]

$^{1}H$ NMR (500 MHz, $CD_3CN$) δ 3.91 (d, *J* = 16.8 Hz, 2H), 3.82 (d, *J* = 16.8 Hz, 2H), 2.90 (s, 3H), 1.20 (s, 12H), 0.08 (s, 1H).

$^{13}C$ NMR (125 MHz, $CD_3CN$) δ 169.1, 83.8, 63.0, 47.1, 25.1.

$^{11}B$ NMR (161 MHz, $CD_3CN$) δ 33.9, 13.5.

HRMS $[M+Na]^+$ calculated for $C_{12}H_{21}B_2NNaO_6$ = 320.1447, found 320.1454.

**(Hydroxymethyl)boronic acid MIDA ester (59)**

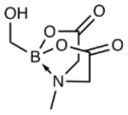

Compound **59** was prepared according to general procedure C from **58** (348 mg, 1.17 mmol, 1.0 eq.). The crude product was purified by flash column chromatography (gradient from 20 to 70% MeCN in DCM) to obtain **59** (159 mg, 850 µmol, 73%) as a white solid.

The recorded NMR data is in accordance with the literature[11].

$^{1}H$ NMR (500 MHz, $CD_3CN$) δ 3.93 (d, *J* = 16.7 Hz, 2H), 3.80 (d, *J* = 16.7 Hz, 2H), 3.23 (s, 2H), 3.02 (s, 3H), 2.34 (s, 1H).

$^{13}C$ NMR (125 MHz, $CD_3CN$) δ 169.4, 63.0, 46.4.

$^{11}B$ NMR (161 MHz, $CD_3CN$) δ 11.3.

HRMS $[M+Na]^+$ calculated for $C_6H_{10}BNNaO_5$ = 210.0544, found 210.0546.

***Tert*-butyl 2-(hydroxymethyl)acrylate (60)**

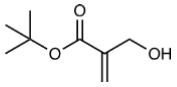

The desired compound **60** was prepared according to a literature procedure.[19] In a 250 mL round-bottom flask, *tert*-butyl acrylate (1.00 g, 7.8 mmol, 1.0 eq.) was dissolved in water (25 mL) and 1,4-dioxane (25 mL). DABCO (2.62 g, 23.4 mmol, 3.0 eq.) was added, followed by formaldehyde (37% aq. solution, 1.74 mL, 23.4 mmol, 3.0 eq.). The reaction was stirred at room temperature for 7 days. $Et_2O$ (100 mL) was added, the layers separated and the organic layer washed with brine (2 x 50 mL). The combined aqueous layers were extracted with $Et_2O$ (3 x 80 mL). The combined organic layers were dried over anhydrous sodium sulfate, filtered and concentrated under reduced pressure. The obtained crude liquid was purified by flash column chromatography (gradient from 0 to 40% EtOAc in hexane) to obtain *tert*-butyl 2-(hydroxymethyl)acrylate (**60**, 0.97 g, 6.1 mmol, 79% yield) as a colourless liquid.

The recorded NMR data is in accordance with the literature[19].

$^{1}H$ NMR (400 MHz, $CDCl_3$) δ 6.16 – 6.14 (m, 1H), 5.74 (dt, *J* = 1.4, 1.4 Hz, 1H), 4.29 (s, 2H), 1.51 (s, 9H).

HRMS $[M+Na]^+$ calculated for $C_8H_{14}NaO_3$ = 181.0835, found 181.0831.

***Tert*-butyl 2-(bromomethyl)acrylate (61)**

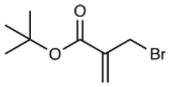

The desired compound **61** was prepared according to a literature procedure.[20] *Tert*-butyl 2-(hydroxymethyl)acrylate (**60**, 865 mg, 5.47 mmol, 1.0 eq.) were dissolved in anhydrous diethyl ether (18 mL) under argon atmosphere. The solution was cooled to -30 °C. $PBr_3$ (208 µL, 2.19 mmol, 0.4 eq.) was added dropwise. The reaction was stirred for 3 hours while slowly warming to 0 °C. Then, cooled to -10 °C and water (11 mL) added. The reaction was allowed to warm to room temperature, additional water (40 mL) and diethyl ether (100 mL) added. Layers were separated and the organic layer washed with brine (50 mL), dried over anhydrous sodium sulfate, filtered and concentrated under reduced pressure. The crude product was purified by flash column chromatography (gradient of 5 to 20% $Et_2O$ in pentane) to obtain *tert*-butyl 2-(bromomethyl)acrylate (**61**, 880 mg, 3.98 mmol, 73%) as a colourless liquid.

The recorded NMR data is in accordance with the literature[20].

$^1$H NMR (400 MHz, $CDCl_3$) δ 6.23 (d, *J* = 0.9 Hz, 1H), 5.85 (dt, *J* = 0.9, 0.9 Hz, 1H), 4.15 (d, *J* = 0.9 Hz, 2H), 1.52 (s, 9H).

$^{13}$C NMR (125 MHz, $CDCl_3$) δ 164.2, 139.1, 128.1, 81.9, 30.0, 28.2.

**(3-(*Tert*-butoxycarbonyl)-1-hydroxybut-3-en-1-yl)boronic acid MIDA ester (62)**

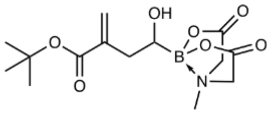

Compound **62** was prepared according to a literature procedure.[21] In a heat-dried two-necked flask equipped with a thermometer and under argon atmosphere, DMSO (68 µL, 963 µmol, 1.2 eq.) was dissolved in anhydrous acetonitrile (6 mL) and the solution cooled to -40 °C. Oxalyl chloride (79 µL, 923 µmol, 1.15 eq.) was added and the reaction stirred for 15 minutes while keeping the temperature between -40 and -30 °C. Then, a solution of hydroxymethyl MIDA boronate (**59**, 150 mg, 802 µmol, 1.0 eq.) in a 1:1 mixture of DMSO/MeCN (anhydrous, 1.5 mL) was added dropwise. The reaction was stirred for further 30 minutes while keeping the temperature between -40 and -30 °C. Then, triethylamine (257 µL, 1.85 mmol, 2.3 eq.) was added and the reaction allowed to warm to 0 °C over 40 minutes. Cooled again to -40 °C and titanocene dichloride (10.0 mg, 40 µmol, 0.05 eq.), triphenyl phosphine (10.5 mg, 40µmol, 0.05 eq.) and zinc powder (262 mg, 4.0 mmol, 5.0 eq.) added, followed by *tert*-butyl 2-(bromomethyl)acrylate (**61**, 355 mg, 1.6 mmol, 2.0 eq.). The reaction was allowed to warm to 0 °C over 30 minutes, then further stirred at 0 °C for 1 hour. Addition of acetic acid (0.9 mL), followed by water (3 mL). After 10 minutes, the reaction was filtered and the filtrate concentrated under reduced pressure. The obtained residue was taken up in EtOAc (20 mL) and half-saturated brine (20 mL). The layers were separated and the aqueous layer further extracted with EtOAc (2 x 20 mL). The combined organic layers were washed with brine, dried over anhydrous sodium sulfate, filtered and concentrated under reduced pressure. The crude product was purified by flash column chromatography on silica gel (gradient from 0 to 15% MeCN in EtOAc) to obtain **62** (159 mg, 486 µmol, 61%) as a white solid.

$^1$H NMR (500 MHz, $CD_3CN$) δ 6.08 (d, *J* = 1.9 Hz, 1H), 5.58 – 5.55 (m, 1H), 3.94 (d, *J* = 17.2 Hz, 1H), 3.91 (d, *J* = 16.2 Hz, 1H), 3.82 (d, *J* = 16.2 Hz, 1H), 3.81 (d, *J* = 17.2 Hz, 1H), 3.40 – 3.33 (m, 1H), 3.02 (s, 3H), 2.64 – 2.57 (m, 2H), 2.29 (ddd, *J* = 14.0, 10.6, 0.9 Hz, 1H), 1.48 (s, 9H).

$^{13}$C NMR (125 MHz, $CD_3CN$) δ 169.8, 168.9, 167.8, 141.2, 126.4, 81.3, 63.2, 63.0, 46.2, 37.1, 28.2.

$^{11}$B NMR (161 MHz, $CD_3CN$) δ 11.0.

HRMS $[M+Na]^+$ calculated for $C_{14}H_{22}BNNaO_7$ = 350.1382, found 350.1385.

**(3-(*Tert*-butoxycarbonyl)but-3-enoyl)boronic acid MIDA ester (63)**

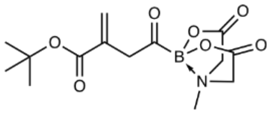

Substrate **62** (156 mg, 477 µmol, 1.0 eq.) was dissolved in acetonitrile (28 mL) and DMP (303 mg, 715 µmol, 1.5 eq.) added. The reaction was stirred open to atmosphere for 45 minutes, then filtered over a pad of celite and the filtrate concentrated. The crude product is purified by flash column chromatography (gradient of 60 to 100% EtOAc in hexane, dry load on celite) to obtain the desired product **63** (114 mg, 351 µmol, 74%) as a pale-yellow solid.

$^{1}$H NMR (500 MHz, $CD_3CN$) δ 6.14 (d, $J$ = 1.7 Hz, 1H), 5.50 – 5.47 (m, 1H), 4.04 (d, $J$ = 16.9 Hz, 2H), 3.90 (d, $J$ = 16.9 Hz, 2H), 3.62 (d, $J$ = 1.0 Hz, 2H), 2.85 (s, 3H), 1.42 (s, 9H).

$^{13}$C NMR (125 MHz, $CD_3CN$) δ 168.9, 137.8, 127.5, 81.3, 63.0, 47.6, 28.1.

$^{11}$B NMR (161 MHz, $CD_3CN$) δ 4.3.

HRMS $[M+Na]^+$ calculated for $C_{14}H_{20}BNNaO_7$ = 348.1225, found 348.1225.

**MIDA itaconylboronate 9k**

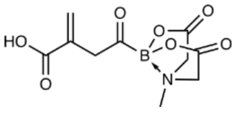

Substrate **63** was dissolved in DCM (0.4 mL) and TFA (374 µL, 4.9 mmol, 30 eq.) added. The reaction was stirred for 30 minutes at room temperature. Then, the reaction was concentrated under reduced pressure at 30 °C. The residue was dissolved in DCM and concentrated again to obtain the desired product **9k** (41 mg, 152 µmol, 93%) as a yellow solid.

Based on the NMR spectra, we assume that the obtained compound exists in two forms. In the structure that is displayed and one in which the itaconyl carboxylic acid coordinates to boron, replacing one of the previous substituents.

$^{1}$H NMR (500 MHz, $CD_3CN$) δ 6.24 and 5.61 (br s, 1H), 6.12 and 5.67 (br s, 1H), 4.11 – 4.01 (m, 2H), 3.97 – 3.86 (m, 2H), 3.65 (br s, 1H), 3.32 – 3.21 and 2.80 – 2.69 (m, 1H), 3.17 – 3.07 and 2.91 – 2.81 (m, 3H).

$^{13}$C NMR (125 MHz, $CD_3CN$) δ 168.9, 136.0, 129.1, 122.3, 63.7, 63.6, 63.1, 63.00 50.3, 47.8, 47.6, 39.5.

$^{11}$B NMR (161 MHz, $CD_3CN$) δ 8.7, 4.4.

HRMS $[M+Na]^+$ calculated for $C_{10}H_{12}BNNaO_7$ = 292.0599, found 292.06.

### 3 96-well based PylRS Screen

A screen of approx. 200 PylRS variants available in the lab was performed using a 96-well setup with a fluorescence readout. Therefore, chemically competent *E. coli* K12 cells were co-transformed with pPylT_sfGFP-N150TAG-H6 (encoding either *Mb* or *Ma* $tRNA_{CUA}$ and the C-terminally H6-tagged sfGFP-N150TAG) and pBK_PylRS (encoding either the *Mb* or *Ma* PylRS mutant) plasmids. After recovery with 0.2 mL of SOC medium (Super Optimal Broth medium containing 20 mM glucose) for 1 h at 37 °C, the cells were cultured overnight in 0.9 mL of 2xYT medium supplemented with tetracycline (17.5 µg/mL) and ampicillin (100 µg/mL) or kanamycin (25 µg/mL) at 37 °C, 200 rpm in a 96-well deep-well plate. After overnight incubation, the cultures were diluted 1:250 into autoinduction medium supplemented with 2 mM **1** or 4 mM **13** and the respective antibiotics (0.5x) in a 96-well deep-well plate. After incubation at 37 °C, 200 rpm for 24 h the cultures were diluted 1:1 into PBS (phosphate-buffered saline) and fluorescence (excitation: 485 nm, emission: 510 nm) was measured using a TECAN SPARK multimode microplate reader.

### 4 PylRS Evolution

Aminoacyl tRNA synthetase evolution was performed on a *M. alvus* PylRS library with five randomized positions (L125X, Y126X, M129X, H227X and Y228X). For the first positive selection[22,23], the library was transformed into electrocompetent K12 cells containing the positive selection plasmid. This plasmid carries the essential gene for chloramphenicol acetyltransferase with a TAG codon (CAT-D111TAG) under a constitutive promoter (cat), *M. alvus* PylT and tetracycline resistance. Cells were rescued with 5 mL of prewarmed SOC medium and incubated at 37 °C for 1 hour. Then, the rescued cells were transferred to 100 mL 2xYT medium containing tetracycline (1x) and ampicillin (1x). The culture was incubated at 37 °C until optical density at 600 nm (OD) reached 0.6. Then, 200 µL of the culture were transferred onto a 15 cm culture dish containing 30 mL AI-agar with the respective antibiotics, 1.2x chloramphenicol and 2 mM of **1**. The plate was incubated at 37 °C for 19 hours. Then, 30 mL 2xYT medium (containing 1x Amp and 1x Tet) were used to scratch the surviving colonies, which were further incubated at 37 °C for 5 hours. Then, the selected library plasmids were isolated using MiniPrep spin columns and purified with a 1% agarose gel.

The obtained library was transformed into electrocompetent DH10β cells containing the negative selection plasmid. This plasmid carries the toxic gene for barnase with two TAG codons (barnase-Q3TAG-D45TAG) under an arabinose promoter, *M. alvus* PylT and chloramphenicol resistance. Cells were rescued with 5 mL of prewarmed SOB medium and incubated at 37 °C for 1 hour. Then, the rescued cells were transferred to 100 mL 2xYT medium containing chloramphenicol (1x) and ampicillin (1x). The culture was incubated at 37 °C until OD reached 0.6. Then, 100 µL of the culture were transferred onto a 15 cm culture dish containing 30 mL LB-agar with the respective antibiotics and 0.2% arabinose. The plate was incubated at 37 °C for 24 hours. Then, 30 mL 2xYT medium (containing 1x Amp and 1x Cam) were used to scratch the surviving colonies, which were further incubated at 37 °C for 3 hours. Then, the selected library plasmids were isolated using MiniPrep spin columns and purified with a 1% agarose gel.

The obtained library was transformed into electrocompetent DH10β cells containing the positive selection double reporter plasmid. This plasmid carries the essential CAT-D111TAG under constitutive promoter (cat), sfGFP-N150TAG under an arabinose promoter, *M. alvus* PylT and tetracycline resistance. Cells were rescued with 5 mL of prewarmed SOC medium and incubated at 37 °C for 1 hour. Then, the rescued cells were transferred to 100 mL 2xYT medium containing tetracycline (1x) and ampicillin (1x). The culture was incubated at 37 °C OD reached 0.6. Then, 4 µL of the culture (diluted to 100 µL) were transferred onto a 15 cm culture dish containing 30 mL AI-agar with the respective antibiotics, 1.6x chloramphenicol and 2 mM of ncAA **1**. The plate was incubated at 37 °C for 22 hours. Colonies were picked and cultured in a 96-well deep-well plate (containing 2xYT medium with 1x Amp and 1x Tet). Large, small, fluorescent and non-fluorescent colonies were picked. The cultures were incubated for 16 hours at 37 °C. Then, cultures were used to inoculate (1:250) a new 96-well deep-well plate containing 0.5 mL AI medium, 0.5x Amp, 0.5x Tet and 2 mM ncAA **1**. A plate without ncAA was prepared as a control. The plates were incubated at 37 °C for 24 hours. Then, cultures were diluted 1:1 into PBS and GFP fluorescence (excitation: 485 nm, emission: 510 nm) was measured using a TECAN SPARK multimode microplate reader. Measurement was repeated after further 24 hours. Plasmids from cultures displaying large fluorescence over the corresponding cultures from the control plate were isolated and sent for sequencing to obtain PylRS mutants able to incorporate the ncAA **1**.

In parallel, the same procedure was performed while omitting ncAA **1** addition during positive selections as a negative control.

### 5 Protein expression and purification

##### 5.1 Expression and purification of Ub- and SUMO2-MeOK mutants

Chemically competent *E. coli* K12 cells were transformed with pBK_pNZ-MeOKRS and a pPylT plasmid containing the H6-tagged protein of interest (POI) variant (see Supplementary Table S2). After recovery with 1 mL SOC medium for 1 hour at 37 °C, the cells were cultured overnight in 5 mL 2xYT medium containing the corresponding antibiotics (1x) at 37 °C, 200 rpm. The overnight culture was diluted to an $OD_{600}$ of 0.05 in 100 mL of fresh 2xYT medium containing the corresponding antibiotics (0.5x) and supplemented with ncAA **1** (4 mM). The culture was incubated at 37 °C, 200 rpm, until $OD_{600}$ reached 0.6, at which point protein expression was induced by addition of 0.05% arabinose (w/v). The culture was further incubated at 37 °C, 200 rpm, for 16 hours. The cells were harvested by centrifugation (4000xg, 20 min, 4 °C) and resuspended in 20 mL lysis buffer (20 mM Tris pH 8.0 at 4 °C, 300 mM NaCl, 30 mM imidazole, 1 mM PMSF, 0.5 mM TCEP). The cells were lysed by sonication with cooling in an ice-water bath. The lysed cells were centrifuged (14000xg, 20 min, 4 °C) and the cleared lysate incubated with Ni beads (Ni Sepharose™ 6 Fast Flow, Cytiva) for 2 hours at 4 °C with agitation. After incubation, the Ni beads were collected on a plastic column and washed with 7 column volumes (CV) of wash buffer (20 mM Tris pH 8.0 at 4 °C, 300 mM NaCl, 30 mM imidazole, 0.5 mM TCEP). The protein was eluted with 1.2 mL wash buffer supplemented with 300 mM imidazole. The elution was further purified via size exclusion chromatography (SEC) using a Superdex™ 75 Increase 10/300 (GE Healthcare) with the storage buffer (20 mM Tris pH 7.5 at 4 °C, 100 mM NaCl). Fractions containing the pure protein were pooled and concentrated using Amicon® centrifugal filter units with a 3 kDa MWCO (Millipore). Protein concentrations were determined using the Pierce™ Rapid Gold BCA Protein Assay Kit (Thermo Fisher Scientific). The purified protein was flash frozen in aliquots using liquid nitrogen and stored at -80 °C until further use.

**5.2 Expression and purification of Histone H2B-K58MeOK**
Chemically competent *E. coli* K12 cells were transformed with pEVOL_oAZ-MeOKRS _PylT and pBAD_Histone_H2B-K58TAG-H6 (see Supplementary Table S2). After recovery with 1 mL SOC medium for 1 hour at 37 °C, the cells were cultured overnight in 5 mL 2xYT medium containing the corresponding antibiotics (1x) at 37 °C, 200 rpm. The overnight culture was diluted 1:500 into fresh AI medium (100 mL) containing the corresponding antibiotics (0.5x) and supplemented with the **13** (4 mM). The culture was incubated for 22 hours at 37 °C, 200 rpm. The cells were harvested by centrifugation (4000xg, 20 min, 4 °C) and resuspended in 25 mL lysis buffer (50 mM Tris pH 7.5 at 4 °C, 500 mM NaCl). The cells were lysed by sonication with cooling in an ice-water bath. The lysed cells were centrifuged (14000xg, 20 min, 4 °C), the supernatant removed and the pellet resuspended in 2% triton in water. The suspension was incubated on ice for 10 minutes, then centrifuged (14000 xg, 20 min, 4 °C). This washing step was repeated twice with 2% triton in water, then once with the lysis buffer (50 mM Tris pH 7.5 at 4 °C, 500 mM NaCl). Then, the pellet was resuspended in wash buffer (50 mM Tris pH 8.0 at 4 °C, 500 mM NaCl, 6 M urea, 30 mM imidazole, 10 mM lysine, 1 mM TCEP) and nearly completely dissolved using ultrasonication. The suspension was centrifuged (14000 xg, 20 min, 4 °C) and the cleared solution incubated with Ni beads (Ni Sepharose™ 6 Fast Flow, Cytiva) for 2 hours at 4 °C with agitation. After incubation, the Ni beads were collected on a plastic column and washed with 7 column volumes (CV) of wash buffer (50 mM Tris pH 8.0 at 4 °C, 500 mM NaCl, 6 M urea, 30 mM imidazole, 10 mM lysine). The protein was eluted with 1.2 mL wash buffer supplemented with 300 mM imidazole. The histone solution was then further purified via cation exchange chromatography using a Resource™ S 1 mL column (Cytiva) with a gradient from 50 mM Tris pH 8.0 at 4 °C, 6 M urea to 50 mM Tris pH 8.0 at °C, 6 M urea, 1 M NaCl. Fractions containing the pure protein were pooled and concentrated using a Amicon® centrifugal filter unit with a 10 kDa MWCO (Millipore). The obtained solution was rebuffered to 20 mM HEPES pH 7.4, 100 mM NaCl using a Pur-A-Lyzer™ Midi 3500 dialysis unit. Dialysis was performed in 6 steps, slowly lowering the urea concentration (3M -> 1.5 M -> 0.75 M -> 0.38 M -> 0.19 M -> 0 M). The solution of folded Histone H2B was then concentrated using a 10kDa Amicon® centrifugal filter unit. Protein concentration was determined using a NanoPhotometer® N60 (IMPLEN). Extinction coefficients were calculated with ProtParam (https://web.expasy.org/protparam/). The purified protein was flash frozen in aliquots using liquid nitrogen and stored at -80 °C until further use.

**5.3 Expression and purification of isocitrate dehydrogenase (IDH) MeOK mutants**
Chemically competent *E. coli* K12 cells were transformed with pEVOL_oAZ-MeOKRS_PylT and pBAD_IDH-K242TAG-H6 (see Supplementary Table S2). After recovery with 1 mL SOC medium for 1 hour at 37 °C, the cells were cultured overnight in 5 mL 2xYT medium containing the corresponding antibiotics (1x) at 37 °C, 200 rpm. The overnight culture was diluted 1:500 into fresh AI medium (50 mL) containing the corresponding antibiotics (0.5x) and supplemented with **13** (4 mM). The culture was incubated for 20 hours at 37 °C, 200 rpm. The cells were harvested by centrifugation (4000xg, 20 min, 4 °C) and resuspended in 20 mL lysis buffer (20 mM Tris pH 8.0 at 4 °C, 300 mM NaCl, 30 mM imidazole, 1 mM PMSF, 1 mM TCEP). The cells were lysed by sonication with cooling in an ice-water bath. The lysed cells were centrifuged (14000xg, 20 min, 4 °C) and the cleared lysate incubated with Ni beads (Ni Sepharose™ 6 Fast Flow, Cytiva) for 2 hours at 4 °C with agitation. After incubation, the Ni beads were collected on a plastic column and washed with 7 column volumes (CV) of wash buffer (20 mM Tris pH 8.0 at 4 °C, 300 mM NaCl, 30 mM imidazole, 0.5 mM TCEP). The protein was eluted with 1.2 mL wash buffer supplemented with 300 mM imidazole. The elution was concentrated and rebuffered in an Amicon® centrifugal filter unit with a 30 kDa MWCO (Millipore) to 20 mM Tris pH 7.5 at 4 °C, 150 mM NaCl, 2 mM TCEP, then kept at 4 °C for 16 hours. The solution was then further purified via size exclusion chromatography (SEC) using a Superdex™ 75 Increase 10/300 (GE Healthcare) with the storage buffer (20 mM Tris pH 7.5 at 4 °C, 150 mM NaCl). Fractions containing the pure protein were pooled and concentrated using a 30 kDa Amicon® centrifugal filter unit. Protein concentrations were determined using a NanoPhotometer® N60 (IMPLEN). Extinction coefficients were calculated with ProtParam (https://web.expasy.org/protparam/). The purified protein was flash frozen in aliquots using liquid nitrogen and stored at -80 °C until further use.

**5.4 Expression and purification of isocitrate dehydrogenase IDH-wt and IDH-K242AcK**
To express wild-type IDH and AcK mutants, the procedure described in section 5.3 was followed. For the wild-type, pPylT_IDH-wt-H6 was used. For K242AcK, cells were transformed with pBK_AcKRS[8] and pPylT_IDH-K242TAG-H6_Mb (see Supplementary Table S2). For AcK mutants, 10 mM AcK was added to the expression culture. The expression culture and the lysis buffer were supplemented with 20 mM nicotinamide to prevent deacylation.

##### 5.5 Expression and purification of GAPDH-K215MeOK

Chemically competent *E. coli* K12 cells were transformed with pEVOL_oAZ-MeOKRS_PylT and pBAD_GAPDH-K215TAG-H6 (see Supplementary Table S2). After recovery with 1 mL SOC medium for 1 hour at 37 °C, the cells were cultured overnight in 5 mL 2xYT medium containing the corresponding antibiotics (1x) at 37 °C, 200 rpm. The overnight culture was diluted 1:500 into fresh AI medium (100 mL) containing the corresponding antibiotics (0.5x) and supplemented with **13** (6 mM). The culture was incubated for 20 hours at 37 °C, 200 rpm. The cells were harvested by centrifugation (4000xg, 20 min, 4 °C) and resuspended in 20 mL lysis buffer (20 mM Tris pH 8.0 at 4 °C, 300 mM NaCl, 30 mM imidazole, one cOmplete™ protease inhibitor tablet (Roche) per 50 mL, 1 mM TCEP). The cells were lysed by sonication with cooling in an ice-water bath. The lysed cells were centrifuged (14000xg, 20 min, 4 °C) and the cleared lysate incubated with Ni beads (Ni Sepharose™ 6 Fast Flow, Cytiva) for 2 hours at 4 °C with agitation. After incubation, the Ni beads were collected on a plastic column and washed with 7 column volumes (CV) of wash buffer (20 mM Tris pH 8.0 at 4 °C, 300 mM NaCl, 30 mM imidazole, 0.5 mM TCEP). The protein was eluted with 1.2 mL wash buffer supplemented with 300 mM imidazole. The elution was concentrated and rebuffered in an Amicon® centrifugal filter unit with a 30 kDa MWCO (Millipore) to 20 mM Tris pH 7.5 at 4 °C, 150 mM NaCl, 2 mM TCEP, then kept at 4 °C for 4 hours. The solution was then further purified via size exclusion chromatography (SEC) using a Superdex™ 200 Increase 10/300 (GE Healthcare) with the storage buffer (20 mM Tris pH 7.5 at 4 °C, 150 mM NaCl). Fractions containing the pure protein were pooled and concentrated using a 30 kDa Amicon®. Protein concentrations were determined using a NanoPhotometer® N60 (IMPLEN). Extinction coefficients were calculated with ProtParam (https://web.expasy.org/protparam/). The purified protein was flash frozen in aliquots using liquid nitrogen and stored at -80 °C until further use.

*Note: We have noticed that it is possible to co-purify small quantities of endogenous E. coli GAPDH which we hypothesize form a heterotetramer with the POI and hence end up in the elution. For the expression of GAPDH-K215MeOK we therefore used 6 instead of 4 mM **13**, pushing expression and thus lowering the corresponding E. coli GAPDH impurity to a negligible fraction.*

##### 5.6 Expression and purification of GAPDH-wt and K215E mutant

To express wild-type GAPDH and the K215E mutant, the procedure described in section 5.5 was followed using the plasmids pPylT_GAPDH-wt-H6 and pPylT_GAPDH-K215E-H6, respectively (see Supplementary Table S2).

##### 5.7 Expression and purification of *E. coli* NfsA and NfsB

Chemically competent *E. coli* BL21 (DE3) cells were transformed with pETM11_H6-TEV-NfsA or pETM11_H6-TEV-NfsB (see Supplementary Table S2). After recovery with 1 mL SOC medium for 1 hour at 37 °C, the cells were cultured overnight in 5 mL 2xYT medium containing the corresponding antibiotics (1x) at 37 °C, 200 rpm. The overnight culture was diluted to an $OD_{600}$ = 0.05 into fresh 2xYT medium (100 mL) containing kanamycin (1x). The culture was incubated at 37 °C, 200 rpm, until $OD_{600}$ reached a value of 0.6. Then, protein expression was induced by addition of 0.4 mM isopropyl β-d-1-thiogalactopyranoside (IPTG). The expression culture was further incubated for 7 hours at 37 °C, 200 rpm. The cells were harvested by centrifugation (4000xg, 20 min, 4 °C) and resuspended in 20 mL lysis buffer (20 mM Tris pH 8.0 at 4 °C, 300 mM NaCl, 30 mM imidazole, 1 mM PMSF). The cells were lysed by sonication with cooling in an ice-water bath. The lysed cells were centrifuged (14000xg, 20 min, 4 °C) and the cleared lysate incubated with Ni beads (Ni Sepharose™ 6 Fast Flow, Cytiva) for 1 hour at 4 °C with agitation. After incubation, the Ni beads were collected on a plastic column and washed with 7 column volumes (CV) of wash buffer (20 mM Tris pH 8.0 at 4 °C, 300 mM NaCl, 30 mM imidazole). The protein was eluted with 1.2 mL wash buffer supplemented with 300 mM imidazole. The elution was concentrated and rebuffered in an Amicon® centrifugal filter unit with a 10 kDa MWCO (Millipore) to 20 mM Tris pH 8.0 at 4 °C, 100 mM NaCl. Protein concentrations were determined using a NanoPhotometer® N60 (IMPLEN). Extinction coefficients were calculated with ProtParam (https://web.expasy.org/protparam/). The purified protein was flash frozen in aliquots using liquid nitrogen and stored at -80 °C until further use.

##### 5.8 Expression and purification of SIRT5

Expresssion and purification of SIRT5 was performed as described previously.[24]

##### 5.9 Protein purification yields

Supplementary Table S3: Protein purification yields (per litre of culture for purified proteins).

| Protein | Yield |
|---|---|
| Ub-K48MeOK | 16 mg/L |
| Ub-K63MeOK | 5 mg/L |
| SUMO2-K11MeOK | 5 mg/L |
| SUMO2-K45MeOK | 6 mg/L |
| H2B-K58MeOK | 5 mg/L |
| IDH-wt | 85 mg/L |
| IDH-K242MeOK | 75 mg/L |
| IDH-K242AcK | 54 mg/L |
| GAPDH-wt | 52 mg/L |
| GAPDH-K215E | 42 mg/L |
| GAPDH-K215MeOK | 23 mg/L |
| NfsA-wt | 51 mg/L |
| NfsB-wt | 81 mg/L |

### 6 On-protein installation of lysine acylations

**General information**

To install lysine acylations on MeOK-modified POIs, synthesized acylboronates **9a-k** were freshly dissolved in DMSO to obtain an 80 or 120 mM stock solution. The stock solution was used for reactions for up to two days, then discarded. For malonylation, the acylboronate stock has to be used immediately as the compound decarboxylates in DMSO solution. If the POI tolerates up to 2.5% of MeCN, one can use MeCN for a more stable stock solution. Reactions were performed in 50 mM citrate buffer with a pH between 5 and 7, most often pH 6.5. After initial optimization on ubiquitin, reaction buffers generally contained 150 mM NaCl due to better compatibility with folded proteins. No difference in reaction performance was observed with or without NaCl. Reactions were performed at 23 °C in 1.5 mL Eppendorf tubes or 0.2 mL PCR tubes. To set up reactions, the reaction tube was filled with the buffer, followed by the acylboronates stock (yielding a reaction solvent that contains 2.5% DMSO). The solution was mixed gently by pipetting up and down. Then, MeOK-modified POI was added from a stock solution and mixed again. The reaction was then left at 23 °C without agitation. Once the installation of the acylation was complete, the modified protein was isolated from the reaction mixture by either rebuffering in Amicon® centrifugal filter units, purification over a size-exclusion column or by dialysis into a storage buffer. Concentrations were assessed using a NanoPhotometer® NP60 (Implen). Acylated proteins were analysed by SDS-PAGE and LC-MS.

**General procedure**

In a 1.5 mL Eppendorf tube, the reaction buffer (50 mM citrate buffer, 150 mM NaCl) was added, followed by the acylboronate stock (80 or 120 mM in DMSO). The solution was mixed gently by pipetting. Then, the MeOK-modified POI was added and mixed again. The reaction was incubated at room temperature until the acylation was completely installed.

**Ub-K63AcK, Ub-K48AcK, SUMO2-K11AcK and SUMO2-K45AcK**

The MeOK-modified POI was diluted into the reaction buffer to a concentration of 20 µM according to the general procedure. The citrate buffer at pH 7 contained 2 mM of the acylboronate for introduction of acetylation. The total reaction volume was 35 µL. After 3 hours, the reaction was analysed by LC-MS.

**Ub-K63 with various acylations**

The reactions were performed according to the general procedure. The exact conditions used are listed in Supplementary Fig. 5 and 6. The total reaction volume was 35 µL.

**Histone H2B-K58ItaK**

The reaction was performed according to the general procedure. 20 µM of H2B-K58MeOK was added to citrate buffer at pH 5.0 containing 1 mM acylboronate (70 µL reaction volume). The reaction was incubated at 23 °C for 7 hours, then added to an equilibrated Pur-A-Lyzer$^{TM}$ Mini 6000 dialysis unit and rebuffered to 20 mM Tris pH 7.2 @ 4 °C, 100 mM NaCl at 4 °C overnight. The dialysis buffer was exchanged and further dialyzed for 2 hours. The obtained acylated histone was analysed by LCMS and then further used for the deacylation assay (Section 7.1).

**IDH-K242AcK**

The reaction was performed according to the general procedure. 10 µM of IDH-K242MeOK was added to citrate buffer at pH 6.5 containing 3 mM acylboronate (500 µL reaction volume). The reaction was incubated at 23 °C for 4.5 hours, then overnight at 4 °C. IDH-K242AcK was isolated from the reaction mixture by rebuffering in a 10K Amicon® centrifugal filter unit to 20 mM Tris pH 7.5 at 4 °C, 100 mM NaCl. The concentration was assessed using a NanoPhotometer® NP60 (Implen) and the protein further analysed by SDS-PAGE and LC-MS.

**IDH-wt treated**

The treated wt control for IDH was subjected to the exact same conditions as described above for IDH-K242AcK.

**GAPDH-K215SucK**

The reaction was performed according to the general procedure. 10 µM of GAPDH-K215MeOK was added to citrate buffer at pH 6.5 containing 3 mM acylboronate (1 mL reaction volume). The reaction was incubated at 23 °C for 6 hours, then another 2 mM acylboronate was added. After further 3 hours, the reaction was kept at 4 °C overnight. The solution was then further purified via size exclusion chromatography (SEC) using a Superdex™ 200 Increase 10/300 (GE Healthcare) with the storage buffer (20 mM Tris pH 7.5 at 4 °C, 150 mM NaCl). The concentration was assessed using a NanoPhotometer® NP60 (Implen) and the protein further analysed by SDS-PAGE and LC-MS.

**GAPDH-wt treated**

The treated wt control for GAPDH was subjected to the exact same conditions as described above for GAPDH-K215SucK.

### 7 Protein Assays

#### 7.1 Deacylation assays

Acylated POIs were incubated in the presence of SIRT5 to assess deacylation activity. More specifically, SIRT5 (0.2 to 1.0 eq.) was added to the acylated POI (generally 10 µM) in the assay buffer (50 mM Tris pH 7.4 at 37 °C, 100 mM NaCl, 5 mM $MgCl_2$, 1 mM DTT and 5 mM $NAD^+$) and incubated at 37 °C. The respective reaction time and SIRT5 quantity is displayed in Supplementary Fig. 13 and 15c. Ub-K63AcK and H2B-K58ItaK were tested at 20 µM concentration. The reaction was assessed by LC-MS.

#### 7.2 IDH enzymatic activity assays

IDH variants were diluted to 10 nM in IDH assay buffer (50 mM Tris pH 7.5, 150 mM NaCl, 10 mM $MgCl_2$ and 0.5 mM $NADP^+$) containing increasing concentrations of D,L-isocitrate (0 – 300 µM). The reaction was started by addition of the enzyme and IDH activity was monitored by measuring the increasing absorbance at 340 nm caused by NADPH formation. Measurements were performed on a Cary 3500 Multicell UV-Vis spectrophotometer (Agilent) in a volume of 2 mL with stirring. Each substrate concentration was measured in triplicates from distinct experiments.

To determine the enzyme kinetics, the absorbance was plotted against time and the initial slope ($v_0$) was obtained by performing linear regression of the first 10 - 20 seconds (250 measurements/second). $v_0$ in units µM/s was plotted against the concentration of D,L-isocitrate. $V_{max}$ and $K_m$ were calculated with a Michaelis-Menten fit (GraphPad Prism 10, GraphPad Software).

#### 7.3 GAPDH RNA binding assay

5'-ATTO532-modified 20A-RNA was dissolved to a concentration of 10 nM in the assay buffer (20 mM Tris pH 7.5 at 23 °C, 50 mM NaCl). GAPDH variants were titrated to the RNA solution and fluorescence anisotropy measured with a JASCO FP-8350 spectrofluorometer. Measurements were performed at 25 °C with excitation wavelength at 530 nm and emission wavelength at 550 nm. To obtain the binding affinities ($K_D$), the change in anisotropy was plotted against the concentration of binding sites (two per GAPDH tetramer) and non-linear regression performed using a one-site specific binding model (GraphPad Prism). For each GAPDH variant, data was collected from three biologically independent experiments ($n$ = 3). Data was analysed using GraphPad Prism 10 (GraphPad Software).

#### 7.4 GAPDH enzymatic activity assays

GAPDH variants were diluted to 2 nM in GAPDH assay buffer (20 mM Tris pH 8.0, 50 mM NaCl, 2.5 mM EDTA, 20 µM $NAD^+$ and 1 mM D-glyceraldehyde-3-phosphate). For measurements in the presence of RNA, 12.5 µM 20A-RNA was added. The reaction was started by addition of 15 mM sodium arsenate and GAPDH activity was monitored by measuring the increasing absorbance at 340 nM caused by NADH formation. Measurements were performed on a Cary 3500 Multicell UV-Vis spectrophotometer (Agilent) in a volume of 100 µL.

To determine the enzyme kinetics, the absorbance was plotted against time and the initial slope ($v_0$) was obtained by performing linear regression of the first 20 seconds (250 measurements/second). Data was collected from three biologically independent experiments ($n$ = 3) and analysed using GraphPad Prism 10 (GraphPad Software).

### 9 NMR, LC-MS and HRMS

**NMR**
All NMR spectra were measured in deuterated solvents at room temperature with a Bruker Avance 400, Bruker Ascend 400, Bruker Ultrashield 400 or a Bruker Avance 500 spectrometer. Chemical shifts in $CDCl_3$ and $CD_3CN$ are referenced to the solvent residual signal. For $CDCl_3$ $^1$H: δ = 7.26 ppm, $^{13}$C: δ = 77.16 ppm and for $CD_3CN$ $^1$H: δ = 1.94 ppm, $^{13}$C: δ = 118.26 ppm (nitrile carbon). All chemical shifts are reported in parts per million (ppm). The spectra were analysed using MestReNova (Mestrelab Research S.L.). The coupling constants are given in Hertz (Hz) and signal multiplicity is characterized as follows: s (singlet), d (doublet), t (triplet), q (quartet), qint (quintet), m (multiplet), br (broad), and combinations thereof.

**HR-MS**
For HR-MS analysis samples were submitted to the Molecular and Biomolecular Analysis Service MoBiAS (ETH Zurich) and analysed on a Bruker Daltonics maXis ESI-QTOF.
**LC-MS**
LC-MS analysis of small molecules was performed on an Agilent 1260 Infinity Series LC system with an Agilent 6130 ESI Single Quadrupole mass spectrometer using a Luna® Omega 3 µM PS C18 (2.1 mm, 100 mm, 100 Å, 3 µm) capillary column (Phenomenex, Torrence, USA). The analysis was performed at RT with a flow rate of 550 µL/min and a gradient of 15-95 % solvent B in 3 min, followed by 1.3 min at 95% solvent B (solvent A: 0.1 % formic acid (FA) in water, solvent B 0.1 % FA in acetonitrile (ACN)).

LC-MS analysis of full-length proteins was performed on an Agilent 1260 Infinity Series LC system with an Agilent 6210 ESI Single Quadrupole mass spectrometer using a Jupiter C4 column (2 mm, 150 mm, 300 Å, 5 µm) capillary column (Phenomenex, Torrence, USA). The analysis was performed at RT with a flow rate of 900 µL/min and a gradient of 10-90% solvent B in 3 min, followed by 0.7 min at 90% solvent B (solvent A: 0.1 % FA in water, solvent B 0.1 % FA in ACN).

Alternatively, LC-MS analysis of full-length proteins was performed on a Bruker-Compact (Q-TOF MS) with chromatographic separation on an Acquity UPLC® Protein BEH C4 column (2.1 x 100 mm, 1.7 µm).

**Author Contributions**
K.L. and T.A.N. envisioned the ‘tag-and-modify’-approach to lysine acylations. T.A.N. synthesized initial carbamoylhydroxylamines and performed attempts to genetically incorporate them into proteins. P.K. synthesized the MeOK ncAAs, evolved the corresponding pNZ-MeOKRS and conducted PylRS screens to identify oAZ-MeOKRS. P.K. expressed and purified Ub, SUMO2, IDH, histone H2B and GAPDH bearing site-specific MeOK. P.K. synthesized MIDA acylboronates, optimized reaction conditions with MeOK-bearing POIs and installed site-specific acylations. P.K. performed nitroreductase experiments, IDH activity assays, GAPDH assays and deacylation assays. P.K. and K.L. analysed the data, and K.L. supervised the study. K.L. and P.K. wrote the paper. with input from T.A.N.

**Fully uncropped and unprocessed gels**

red boxes shown as cropped gels in Figure 1c and 3d

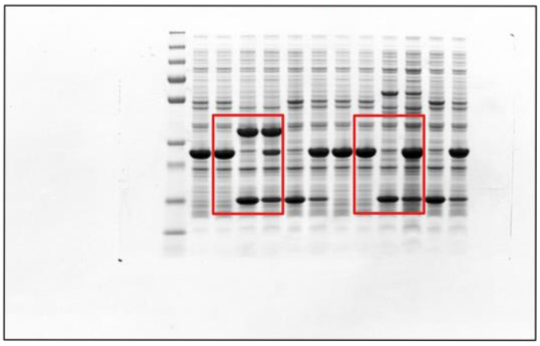

red box shown as cropped gel in Figure 4d

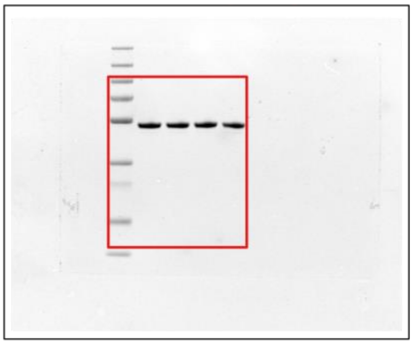

red box shown as cropped gel in Figure 5e

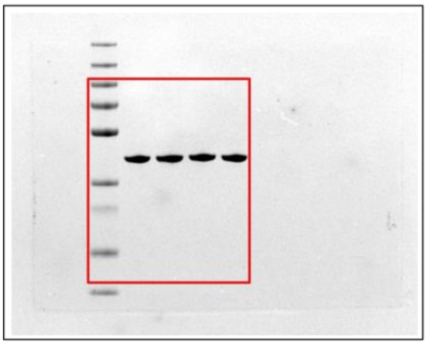

red box shown as cropped gel in Figure S2a

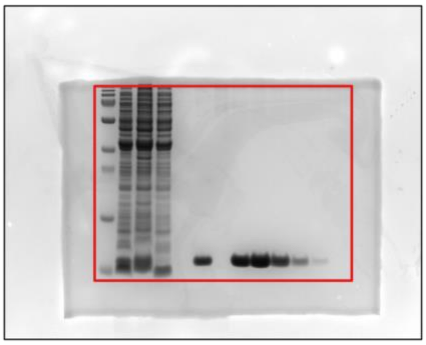

red box shown as cropped gel in Figure S2b

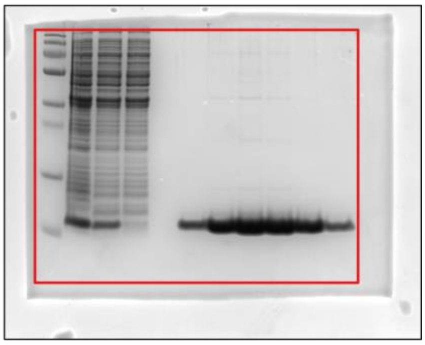

red boxes shown as cropped gels in Figure S2b

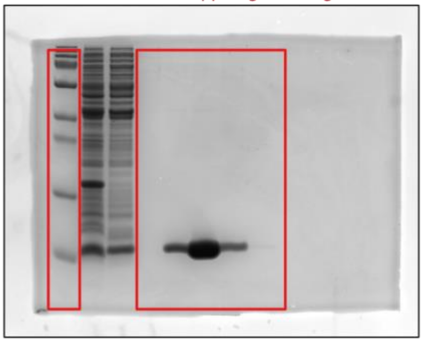

red boxes shown as cropped gels in Figure S2c and Figure S2d

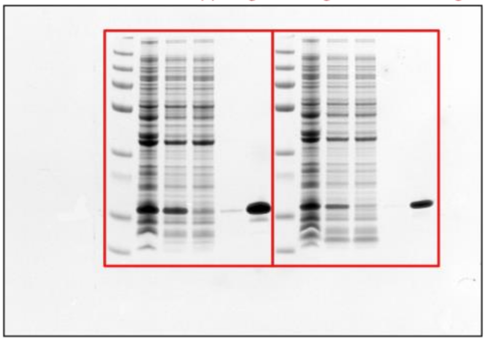

red box shown as cropped gel in Figure S2c

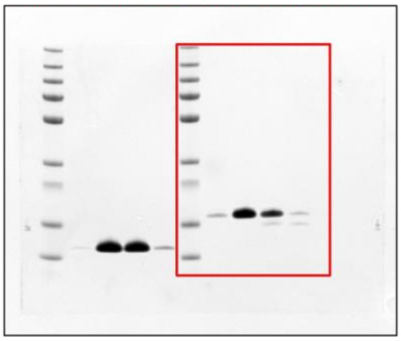

red box shown as cropped gel in Figure S2d

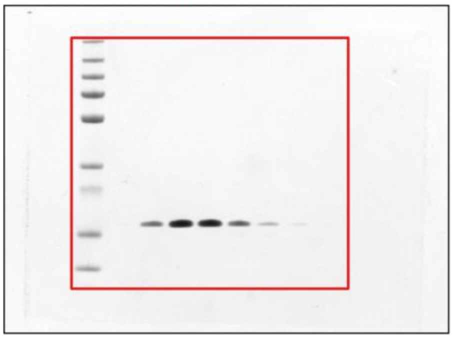
